# The genomic landscape of clonal hematopoiesis in Japan

**DOI:** 10.1101/653733

**Authors:** Chikashi Terao, Akari Suzuki, Yukihide Momozawa, Masato Akiyama, Kazuyoshi Ishigaki, Kazuhiko Yamamoto, Koichi Matsuda, Yoshinori Murakami, Steven A McCarroll, Michiaki Kubo, Po-Ru Loh, Yoichiro Kamatani

**Affiliations:** Laboratory for Statistical and Translational Genetics, RIKEN Center for Integrative Medical Sciences, Kanagawa, Japan; Clinical Research Center, Shizuoka General Hospital, Shizuoka, Japan; The Department of Applied Genetics, The School of Pharmaceutical Sciences, University of Shizuoka, Shizuoka, Japan; Laboratory for Autoimmune Diseases, RIKEN Center for Integrative Medical Sciences, Yokohama, Japan; Laboratory for Genotyping Development, RIKEN Center for Integrative Medical Sciences, Yokohama, Kanagawa, Japan; Department of Ophthalmology, Graduate School of Medical Sciences, Kyushu University, Fukuoka, Japan; Laboratory of Genome Technology, Human Genome Center, Institute of Medical Science, The University of Tokyo, Tokyo, Japan; Laboratory of Clinical Genome Sequencing, Department of Computational Biology and Medical Sciences, Graduate School of Frontier Sciences, The University of Tokyo, Tokyo, Japan; Division of Molecular Pathology, Institute of Medical Science, The University of Tokyo, Tokyo, Japan; Program in Medical and Population Genetics, Broad Institute of MIT and Harvard, Cambridge, MA, USA; Stanley Center for Psychiatric Research, Broad Institute of MIT and Harvard, Cambridge, Massachusetts, USA; Department of Genetics, Harvard Medical School, Boston, Massachusetts, USA; Division of Genetics, Department of Medicine, Brigham and Women’s Hospital and Harvard Medical School, Boston, MA, USA; Kyoto-McGill International Collaborative School in Genomic Medicine, Kyoto University Graduate School of Medicine, Kyoto, Japan

## Abstract

Cancer and aging are widely assumed to involve similar biological trajectories in different populations, though such biology has not been systematically characterized on a population scale. Clonal hematopoiesis with acquired mutations is a pre-cancerous condition that becomes common with advancing age. Here we describe shared and population-specific patterns of genomic mutation revealed by 33,250 autosomal mosaic chromosomal alterations (mCAs) we ascertained from 179,417 Japanese participants in the BioBank Japan cohort and compared to analogous analyses of UK Biobank. In this long-lived Japanese population, mCAs were detected in more than 35.0% (s.e. 1.4%) of individuals older than 90 years, suggesting that such clones trend toward inevitability with aging. Japanese and Europeans exhibited key differences in the genomic locations of mutations in their respective hematopoietic clones, which anticipated the populations’ relative rates of CLL (more common among Europeans) and T-cell lymphoma (more common among Japanese). Multiple mutational precursors of CLL (including trisomy 12, 13q loss, and 13q CNN-LOH) were also 2-6x less common among the Japanese, suggesting that these populations differ in selective pressures on clones long before the development of clinically apparent CLL. Japanese and UK populations also exhibited very different rates of clones arising from B- and T-cell lineages, anticipating the relative rates of B- and T-cell cancers in these populations. We identified six new loci at which inherited variants predispose to mCAs that duplicate or remove inherited risk alleles, including large-effect rare variants at *NBN, MRE11*, and *CTU2* (OR=28-91). Our results motivate further exploration of the global genomic landscape of clonal hematopoiesis and cancer, and point to population-specific selective pressures on hematopoietic clones.

---

Clonal hematopoiesis involving cells with genomic alterations commonly occurs in older individuals and confers risk of hematological malignancies and overall mortality^1-10^. Clones harbor diverse mutations on every chromosome. This spectrum of mosaic events exhibits a correspondingly wide range of etiologies—including inherited genetic components^10-13^ —and effects on subsequent health, motivating deeper study of specific events in large, well-powered cohorts.

Though populations can differ greatly in their rates of specific kinds of cancer, precancerous mosaicism in large cohorts of non-European ancestry is currently unexplored.

We searched for mosaic chromosomal alterations (mCAs) in blood DNA microarray data from 179,417 participants in the BioBank Japan (BBJ) cohort^14^ (Methods). We ascertained mCAs by analyzing allele-specific hybridization intensities for 515,355 genotyped autosomal variants (Table S1), analyzing these data with a recently developed approach that detects imbalanced abundance of an individual’s two inherited haplotypes by utilizing the long-range haplotype phase information that is present in large cohorts^10,15^ (Methods).

This analysis detected 33,250 autosomal mCAs (at FDR = 0.05) in 27,910 unique individuals (Fig. 1 and Fig. S1-S22). (The high rate of events, relative to a contemporaneous analysis of UK Biobank, reflects that BBJ participants were older (mean age at enrollment 62.8 years, s.d. 14.5 years, range 0-113 years) and a larger fraction were male (54.1%)^14^.) Of these mutations, 5,233 were confidently classified as mosaic deletions, 10,431 as copy-number neutral loss of heterozygosity (CNN-LOH) events, and 4,044 as duplications; the remaining 13,452 events were present at cell fractions too low to confidently determine copy number (Fig. 2a and Table S2). Some 4,156 individuals harbored two or more detectable mCAs (Table S3); analysis of mosaic cell fractions suggested that these events were usually present in distinct clones (Fig. S23). The mCA detection rate was broadly consistent across genotyping arrays (Table S4) and across cases of the 47 diseases systematically surveyed by BBJ (Table S5); mCAs were strongly enriched (OR=1.93 (1.66-2.24), p=2.0×10^-17^) only among individuals with hematological cancers at registry, as expected from earlier work.

**Figure 1.**
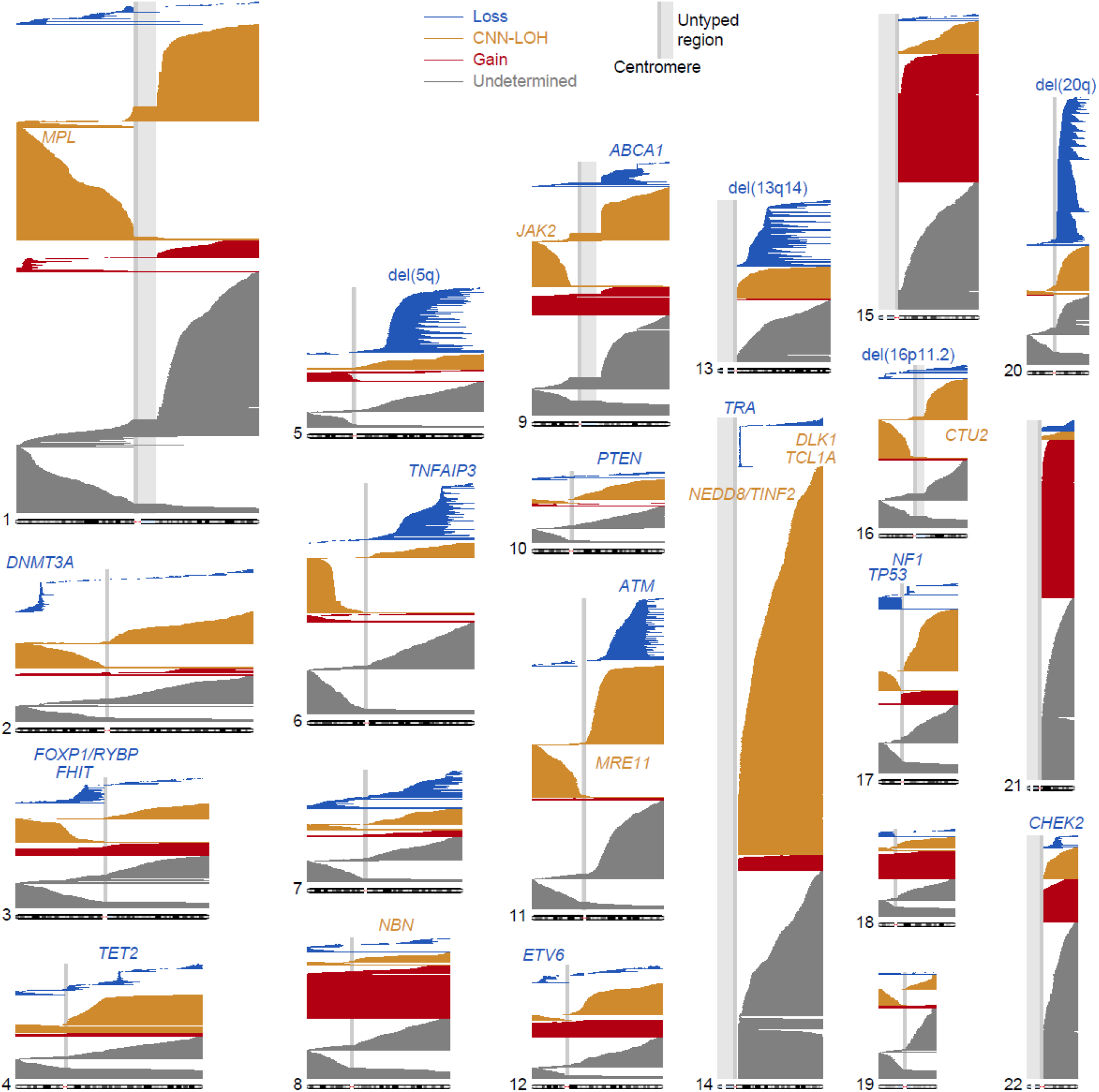
Genomic locations of 33,250 autosomal mCAs detected in 27,910 unique BioBank Japan participants. Loss, CNN-LOH, and gain events are plotted as blue, orange, and red horizontal lines, respectively. Events with undetermined copy number are plotted in grey. Commonly deleted regions are labeled in blue; loci associated with CNN-LOH mutations in *cis* are labeled in orange.

**Figure 2.**
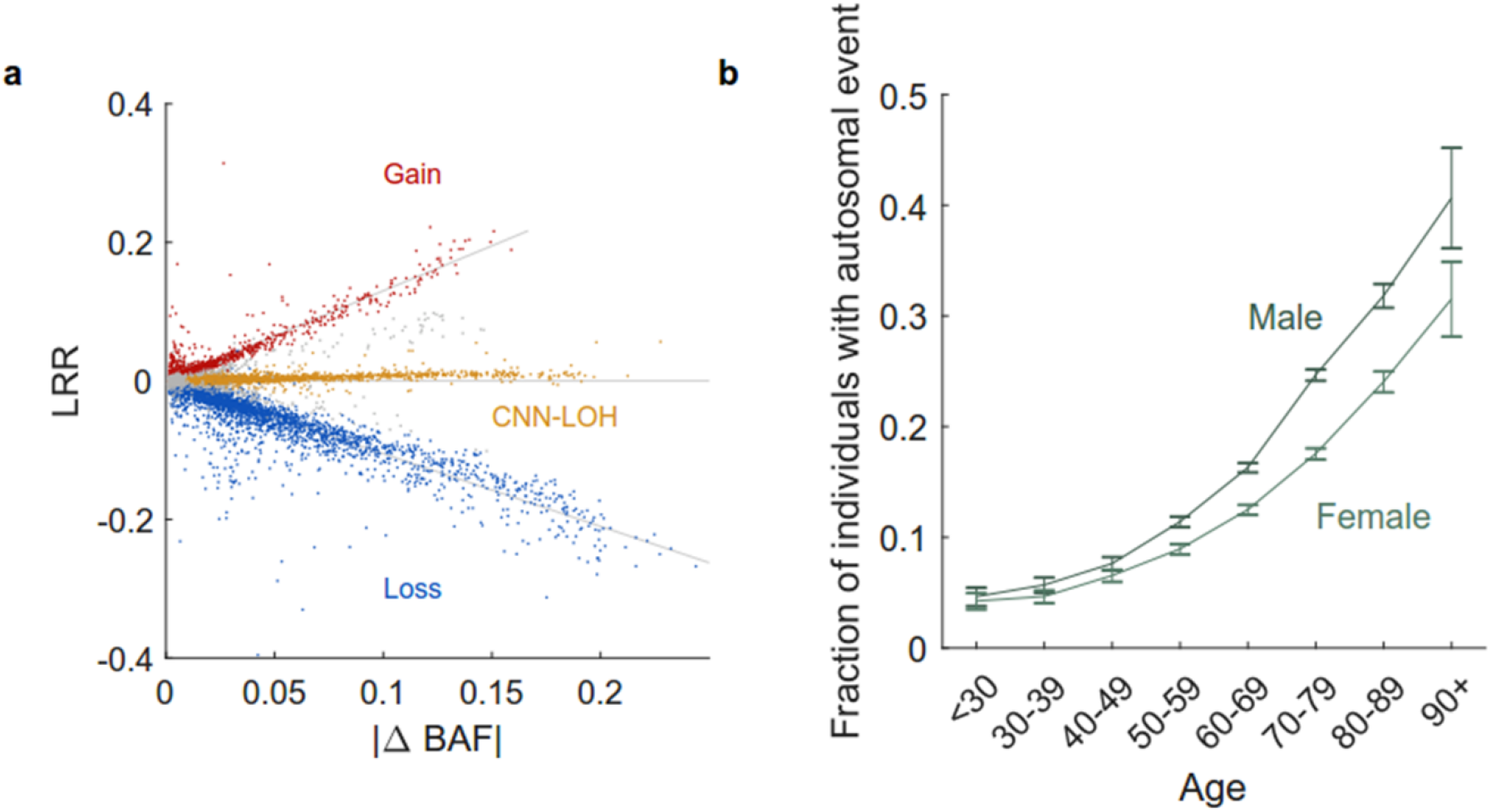
Classification of mCAs and enrichment in subsets of BBJ participants. **a.** Classification of mCAs as loss, CNN-LOH, or gain events using log R ratio (LRR, measuring total DNA abundance) and B allele frequency deviation from 0.5 (|ΔBAF|}, measuring allelic imbalance; Methods). Unclassified events are indicated in grey. **b.** Frequency of detectable mosaicism stratified by age and sex. Error bars, 95% CIs. Numeric data are provided in Tables S2 and S6.

The long-lived Japanese population revealed that clonal hematopoiesis with mCAs becomes extremely common in the very old: detectable mosaicism reached 40.7% (standard error 2.3%) in males and 31.5% (standard error 1.7%) in females over the age of 90 (Fig. 2b and Table S6). This trend toward inevitability in the elderly has not previously been observed for mosaic chromosomal alterations^1,2,5,6,10^ (Supplementary Note). Mosaic chromosomal alterations on different chromosomes and with different copy number changes exhibited various degrees of enrichment in males and in the elderly (Fig. S24, Tables S7-S8 and Supplementary Note) and in individuals with anomalous blood counts (Table S9), suggesting a spectrum of biological processes involved in the development of different clones.

To compare the genomic distributions of mCAs in the Japanese and UK populations, we co-analyzed BBJ mCAs together with 19,632 autosomal mCAs detected in parallel work^16^ on the UK Biobank (UKB) cohort^17,18^ (Fig 3 and Fig S25-46).

**Figure 3.**
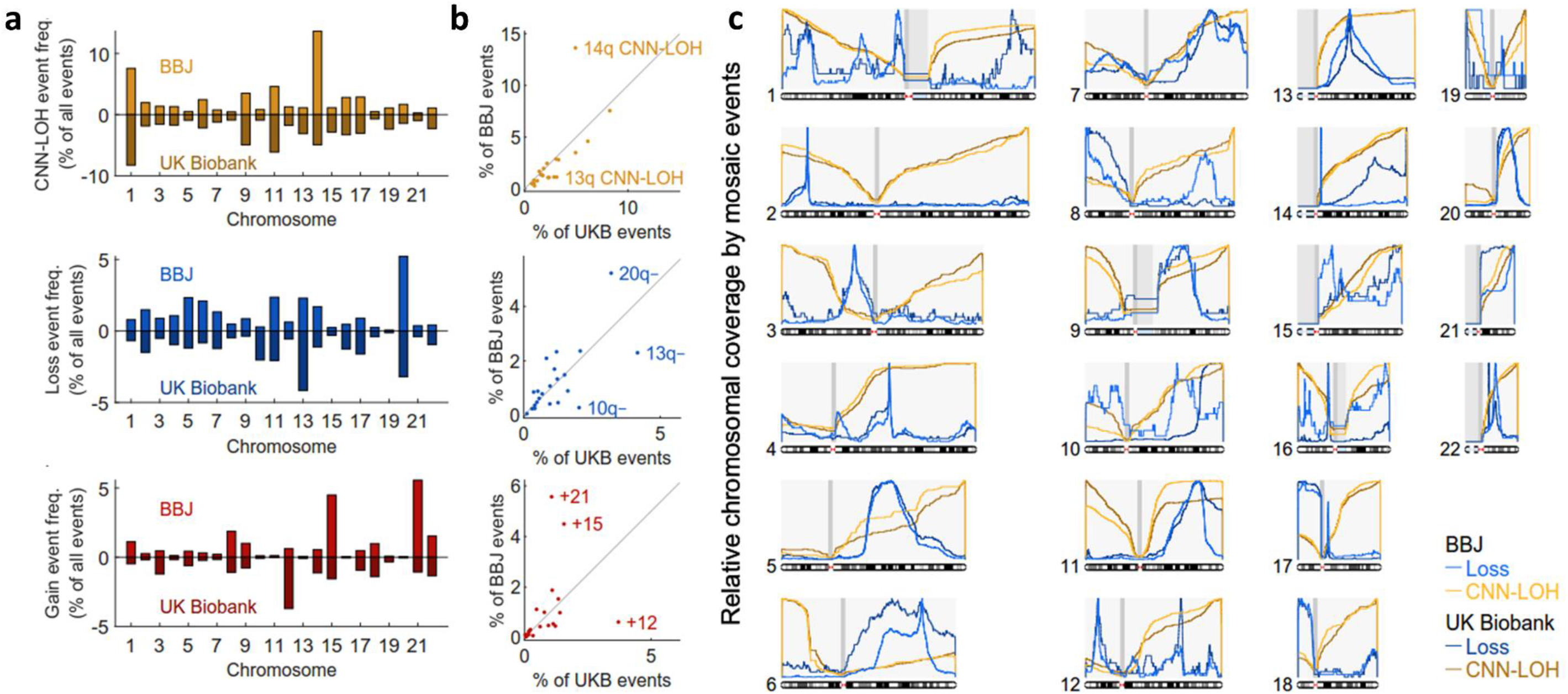
Comparison of genomic distributions of mCAs in BioBank Japan and UK Biobank. **a**,**b.** Distribution of mCAs by chromosome and copy number in BioBank Japan and UK Biobank. **c.** Chromosomal coverage of loss and CNN-LOH events in BioBank Japan and UK Biobank. Curves indicate frequencies at which each chromosomal position is contained in loss (resp. CNN-LOH) events, normalized to 1 on each chromosome. Numeric data are provided in Table S12.

The Japanese have a 10-fold higher incidence of adult T-cell leukemias ^19^ and 5-fold lower incidence of chronic lymphocytic leukemia (CLL, a B-cell malignancy) compared to Europeans^20,21^. Our analysis indicated that, even among people without cancer in both populations, Japanese and UK populations have dramatically different rates of hematopoietic clones arising from the B- and T-cell lineages, as evidenced by deletions that report on V(D)J recombination in developing T and B lymphocytes and thus identify clonal expansions in the T- and B-cell lineages. Mosaic deletions at the TCR alpha locus on 14q (indicating clonal expansion in the T-cell lineage; Supplementary Note) were common in BBJ but rare in UKB (82% vs 11% of chr14 loss events in BBJ and UKB, respectively); in contrast, deletions at the *IGH* and *IGL* immunoglobulin loci (indicating clonal expansion in the B-cell lineage) were common in UKB but rare in BBJ (5% vs 39% of chr14 loss events and 2% vs 58% of chr22 loss events in BBJ and UKB, respectively) (Fig3c, FigS38 and S46). (We verified that these differences did not arise from differences in genomic coverage by the genotyping arrays used by BBJ and UKB (Fig. S47).) Clones arising from the T-cell lineage (as evidence by deletions at *TRA*) were also associated with elevated lymphocyte counts (Table S10-S11) l. Thus, the differences in rates of B- and T-cell malignancies between Japanese and UK populations appear to be foreshadowed in rates of subclinical clonal expansions.

mCAs affect the various human chromosomes at different frequencies. Chromosome-specific CNN-LOH frequencies varied in a way that strongly correlated between BBJ and UKB (*R*=0.73, *p*=0.00013), with the exception of 14q (more common in BBJ) and 13q (more common in UKB). In contrast, the most common loss and gain events in each population (including 20q-, 13q-, 10q-, +21, +15, and +12 events) tended to be much more common in one population than the other (Fig. 3a,b and Table S12).

A clear pattern among the most strongly population-differentiated mutations involved the 2- to 6-fold lower frequency in BBJ of +12, 13q-, and 13q CNN-LOH events (Fig. 3b): all three mutations are commonly observed in chronic lymphocytic leukemia (CLL)^22,23^ and in individuals who later develop CLL^10^. Considering the 4-5 times lower incidence of CLL in East Asians, the parallel difference in population frequencies of clones with each of these CLL-associated mutations suggests that this population difference originates in reduced selective advantage for clones with (a variety of) CLL precursor mutations.

The sub-chromosomal distributions of mosaic deletion events were broadly similar between BBJ and UKB while exhibiting a few notable differences (Fig. 3c, Fig. S25-S48, Table S13 and Supplementary Note). Focal deletions frequently targeted *DNMT3A, TET2, ETV6, NF1*, and *CHEK2*, as in UK Biobank and previous studies^1,2,5,6,10^ (Fig. 1 and Fig. 3c). Interestingly, the CLL-related deletion region at 13q14 was less focal in BBJ than in UKB (Fig. 3c), involving longer deletions in a pattern more similar to the 20q, 5q, and 11q deletion regions. We observed new focal deletion regions in BBJ at *FHIT* on 3p, *TNFAIP3* on 6q, *ABCA1* on 9q, and *PTEN* on 10q (Fig. 3c, Table S14 and S15); *FHIT, TNFAIP3*, and *PTEN* are known tumor suppressor genes associated with blood cancers ^24-26^.

Recent work has established an inherited component of clonal hematopoiesis involving both common variants that slightly increase risk (of clones with any mutation)^11-13,27^ and rare variants that strongly predispose to developing clones with specific mCAs in *cis*^10^. The large number of mCAs detected among the Japanese, together with the presence of a distinct set of low-frequency alleles in Japan, could enable detection of additional risk loci. To identify inherited variants associated with mCAs, we first performed association tests aimed at detecting CNN-LOH events in *cis* that promoted clonal expansion by making risk alleles homozygous or removing them from the genome^10^. We tested variants imputed into BBJ using the 1000 Genomes Phase 3 reference panel^28^ together with 1,037 sequenced Japanese samples^29^, setting a significance threshold of *p*<5×10^-9^ (Methods). We further performed binomial tests to determine whether each risk allele was consistently duplicated or removed by CNN-LOH events (in individuals heterozygous for the risk allele, Methods).

We identified five new loci at which inherited variants associated with mosaic CNN-LOH events in *cis* (and replicated previously reported associations at *JAK2* ^30-32^ and *MPL*^10^) (Table 1, Fig. 4, Fig. S49-S52, and Supplementary Note). Three of the new loci—*NBN, MRE11*, and *CTU2*—involved rare variants with large effects. At *NBN*, the rare stop-gained variant rs756831345 on 8q associated strongly (OR=91 (52-159), *p*=9.8×10^-23^) with 8q CNN-LOH events, which consistently duplicated the *NBN* risk allele (*p*=0.00012; Table 1 and Fig. 4a). At *MRE11*, a very rare intronic variant (probably tagging a different causal variant; Supplementary Note) on 11q associated strongly (OR=37 (17-84), *p*=2.6×10^-9^) with 11q CNN-LOH events, which always duplicated the *MRE11* risk allele (*p*=0.016; Table 1 and Fig. 4b). *NBN, MRE11*, and *RAD50* (which did not exhibit a similar association; Table S16) encode the components of the MRN double strand break repair complex, which recruits ATM in response to DNA damage, leading to phosphorylation of p53 and CHK2 and initiation of cell cycle arrest, apoptosis, or DNA repair^33^. Together with the observations of focal deletions at *ATM, TP53*, and *CHEK2* (Fig. 1) and rare *ATM* risk alleles for CNN-LOH events in *cis*^10^, these results indicate a key role of DNA damage-response dysfunction in clonal hematopoiesis.

**Table 1.** Genome-wide significant associations between inherited variants and mosaic chromosomal alterations.

| Mosaic event | Locus | Chr | Pos | Variant | REF | ALT | AF | *P* | OR (95%CI) | Allelic imbalance in hets ($N_{\text{REF}}:N_{\text{ALT}}$) | Allelic imbalance *P* |
| --- | --- | --- | --- | --- | --- | --- | --- | --- | --- | --- | --- |
| Novel associations with mCAs in *cis* |  |  |  |  |  |  |  |  |  |  |  |
| 8q CNN-LOH | *NBN* | 8 | 90949282 | rs756831345 | C | A | 0.00061 | $9.8 \times 10^{-23}$ | 91 (52-159) | 0:14 | 0.00012 |
| 11q CNN-LOH | *MRE11* | 11 | 94160189 | 11:94160189 | G | A | 0.00011 | $2.6 \times 10^{-9}$ | 37 (17-84) | 0:7 | 0.016 |
| 14q CNN-LOH | *NEDD8/TINF2* | 14 | 24711798 | rs28372734 | C | G | 0.073 | $1.0 \times 10^{-11}$ | 1.62 (1.42-1.85) | 58:176 | $5.2 \times 10^{-15}$ |
| 14q CNN-LOH | *TCL1A* | 14 | 96180242 | rs1122138 | C | A | 0.05 | 0.015 | 0.88 (0.79-0.98) | 231:107 | $1.3 \times 10^{-11}$ |
| 14q CNN-LOH | *DLK1* | 14 | 101175967 | rs10873520 | G | A | 0.30 | $7.1 \times 10^{-39}$ | 1.38 (1.31-1.44) | 1121:689 | $2.5 \times 10^{-24}$ |
| 16q CNN-LOH | *CTU2* | 16 | 88781475 | rs200779411 | C | T | 0.00065 | $7.3 \times 10^{-20}$ | 28 (17-45) | 2:11 | 0.022 |
| Previously reported associations with mCAs in *cis* |  |  |  |  |  |  |  |  |  |  |  |
| 1p CNN-LOH | *MPL* | 1 | 45444734 | 1:45444734 | G | A | 0.00016 | $5.3 \times 10^{-18}$ | 54 (30-100) | 14:0 | 0.00012 |
| 9p CNN-LOH | *JAK2* | 9 | 5026293 | rs2183137 | A | G | 0.24 | $1.1 \times 10^{-11}$ | 1.68 (1.46-1.95) | 54:155 | $1.7 \times 10^{-12}$ |
| Novel associations with mCAs in *trans* |  |  |  |  |  |  |  |  |  |  |  |
| 14q CNN-LOH | *TERT* | 5 | 1287194 | rs2853677 | A | G | 0.31 | $1.5 \times 10^{-22}$ | 1.27 (1.21-1.33) | - | - |
| chr15 gain | *MAD1L1* | 7 | 1975624 | rs12699483 | C | G | 0.42 | $6.9 \times 10^{-23}$ | 1.61 (1.46-1.77) | - | - |

Chr, chromosome. Pos, base pair position (hg19). REF, reference allele. ALT, alternate allele. AF, allele frequency in controls. *P*, association *P*-value. OR, odds ratio (using ALT as effect allele). $N_{\text{REF}}$ (resp. $N_{\text{ALT}}$ ), number of individuals heterozygous for the variant in which a mosaic CNN-LOH event in *cis* made the REF allele (resp. ALT allele) homozygous and removed the other allele.

**Figure 4.**
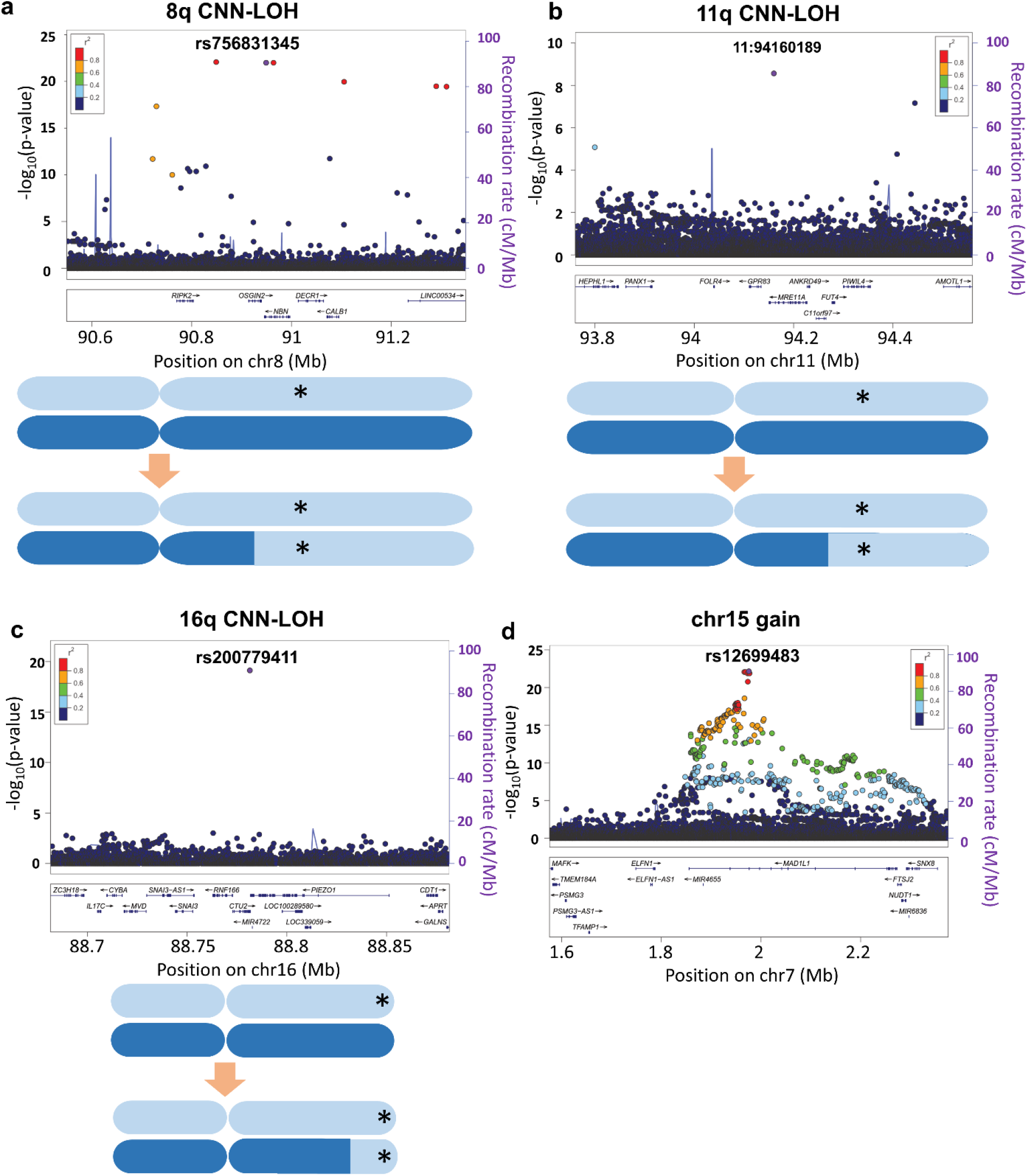
Association signals at the *NBN, MRE11, CTU2*, and *MAD1L1* loci. Rare variants in *NBN, MRE11*, and *CTU2* associate with clonal expansions of CNN-LOH events in *cis* that make the risk alleles homozygous. Common variants at *MAD1L1* associate with mosaic gains of chromosome 15 in *trans*. Lead associated variants are indicated in purple; other variants are color-coded according to LD with lead variants.

At *CTU2*, the rare missense variant rs200779411 associated strongly (*p*=7.3 x10^-20^, OR=28 (17-45)) with 16q CNN-LOH events, which consistently duplicated the *CTU2* risk allele (*p*=0.022; Table 1 and Fig. 4c). *CTU2* encodes a component of the cytosolic thiouridylase complex, which is required for maintenance of genome integrity^34^. The missense variant rs200779411 was predicted to be probably damaging by PolyPhen-2^35^ and deleterious by SIFT^36^, suggesting that impaired *CTU2* function may promote clonal expansion by reducing genome stability.

To additionally detect inherited variants associated with mCAs in *trans*, we performed genome-wide association tests on each mCA type (classifying events by chromosome and copy number), setting a genome-wide significance threshold of *p*<5×10^-11^ to account for multiple hypotheses tested (Methods). Two *trans* associations reached significance: common variants in *MAD1L1* associated with gains on chromosome 15, and common variants in *TERT* (previously associated with mosaic *JAK2* V617F mutation^12^) associated with 14q CNN-LOH (Table 1, Fig. 4d, Fig. S52, and Supplementary Note). At *MAD1L1*, a cluster of five SNPs in near-perfect linkage disequilibrium (including the missense variant rs1801368) associated (*p*=6.9 ×10^-23^, OR=1.61 (1.46-1.77); Table 1 and Fig. 4d) with chromosome 15 gain events (mostly full trisomies; Fig. 1). We replicated this association in the UK population with a slightly reduced effect size (*p*=5.1×10^-4^, OR=1.40 (1.16-1.69) for rs1801368). *MAD1L1* encodes a component of the mitotic spindle assembly checkpoint (SAC) that ensures proper chromosome segregation^37^. The *MAD1L1* risk allele was also previously observed to increase risk of mosaic Y chromosome loss^13^, consistent with a mechanism involving mis-segregation of chromosomes during mitosis due to impaired SAC function. Lending further support to this hypothesis, the risk haplotype was estimated to also increase risk for large (arm-level or whole-chromosome) gain events in 9 out of 10 chromosomes with at least 50 such events (binomial *p*=0.02; Table S17).

A comparison of the mCA risk loci detected in BBJ with previously-reported loci from UK Biobank^10^ revealed ways in which genetic background can differentially shape clonal hematopoiesis in different populations. Four of the risk variants we found in BBJ (at *NBN, MRE11, NEDD8*/*TINF2*, and *CTU2*) were present at much lower allele frequencies in Europeans^38^ (Table S18). Conversely, all rare variants previously associated with mCAs in UKB (at *MPL, FRA10B, ATM*, and *TM2D3*/*TARSL2*)^10^ were absent from the 1000 Genomes and Japanese WGS imputation panels, with the absence of *FRA10B* fragile alleles explaining the lack of 10q25.2-qter deletions in BBJ (Fig. 1). Notably, *MPL* variants were associated with 1p CNN-LOH events in both BBJ and UKB despite risk alleles in each cohort being population-private, demonstrating a shared path to mosaicism initiated by independent variation.

Clonal hematopoiesis has previously been linked to poorer health outcomes, with various types of mosaic events observed to increase risk of future blood cancers, mortality, and cardiovascular disease^1-4,10,39^. To dissect the link between mCAs and mortality, we analyzed mortality outcomes (including cause of death) available for ∼72% of the cohort^40^ (Supplementary Methods).

We observed a nearly five-fold increase in the risk of death due to leukemia (HR=4.70 (3.26--6.78); Fig. S53a, Table S19 and Supplementary Note). The increased risk of leukemia mortality did not appear to extend to other hematological malignancies (malignant lymphoma and multiple myeloma; Fig. S53a and Table S19). We also did not observe significantly increased risk of cardiovascular mortality, suggesting that previous associations of clonal hematopoiesis (primarily involving point mutations in *DNMT3A, TET2, JAK2*, and *ASXL1*) with cardiovascular outcomes^4,39^ may be limited to specific mosaic events (Fig. S53a and Table S19). To refine the association between mosaic status and leukemia mortality, we partitioned mosaic events by chromosome and copy-number change (Methods), identifying seven mCAs with significant (*p*<0.05/120, Cochran-Mantel-Haenszel test), large effects on leukemia mortality risk (Fig. S53b and Table S20). Mosaic cell fraction and the number of mosaic events carried by an individual each associated with further increases in leukemia mortality risk (Fig. S53c,d and Table S21-S22). Mosaic status increased risk of overall mortality (HR=1.10 (1.05-1.16), p=2.7×10^-5^; Table S19), an association that was driven by mCAs in chromosomes 9 and 14 (HR>1.3 and p<0.00078; Table S23).

Our study of clonal hematopoiesis in Japan enabled the first detailed comparison of the genomic landscape of clonal hematopoiesis between populations, revealing broad overall similarities as well as intriguing population differences. A clear pattern among these results was that population differences in cancer rates are anticipated by population differences in sub-clinical clonal expansions, at multiple levels: (i) in specific cell lineages (including B- and T-cell lineages), and (ii) with specific cancer-associated mutations (the +12, 13q-, and 13q CNN-LOH mutations that are hallmarks of CLL). These results point toward population differences in the clonal advantages that are gained by specific mutations in specific genetic and environmental contexts.

The opportunity to probe both acquired and inherited genetic variation in Japan at large sample size enabled further insights into the influences of inherited variation on clonal hematopoiesis: several risk loci pointed to a key role of maintenance of genomic integrity, with corroborating evidence from loci implicated by focal deletions. We anticipate that future studies of even larger, more diverse cohorts will lead to more insights into the genetic architecture of clonal hematopoiesis and inform clinical management of specific pre-cancerous genomic alterations.

## Methods

### The BioBank Japan cohort

All the subjects analyzed in this study were participants of the BioBank Japan Project (BBJ). The BBJ is a multi-hospital-based registry that collected clinical information, DNA, and serum samples from approximately 200,000 patients with one or more of 47 target diseases at a total of 66 hospitals between fiscal years 2003 and 2007^14^. The case proportions correlated well with prevalence in the Japanese population and all of the study participants were diagnosed by medical doctors as described elsewhere^14^. This project was approved by the ethics committees of RIKEN Center for Integrative Medical Sciences and the Institute of Medical Sciences, the University of Tokyo. Written informed consent was obtained from all of the participants.

### Genotyping subjects in the BBJ

Subjects were genotyped in three batches using different arrays or set of arrays, namely: (1) a combination of Illumina Infinium Omni Express and Human Exome, (2) Infinium Omni Express Exome v1.0, and (3) Infinium Omni Express Exome v1.2 (Table S1). The SNP content of the three methods was quite similar. DNA was obtained from blood samples for all but one individual (for which DNA was obtained from oral mucosa; this sample was negative for mosaic events).

We excluded outliers from East Asian clusters in a plot in which we projected BBJ subjects in combination with 1000 Genomes Project samples^41^ in the principal component (PC)1 and PC2 space. We also excluded samples genetically identical to another sample, samples with call rates less than 0.98, and samples whose reported sex information was not supported by genotypes in the X chromosome. We further excluded three samples with evidence of potential contamination (as suggested by low-cell-fraction mosaic events called on many chromosomes^6,10^), leaving 179,417 samples for analysis. We used plink1.9 software^42^ to handle the genotyping data.

### Genotyping intensity data used for calling mosaic events

To call mosaic events, we analyzed genotyping intensity data for variants in the intersection of the three arrays used for BBJ genotyping (namely, Illumina Infinium Omni Express and Infinium Omni Express Exome v1.0 and v1.2) to allow analysis of the same set of variants in all individuals. We did not include variants contained in Human Exome array (see above) when calling mosaic events to avoid batch effects arising from different arrays. We used the variants in Human Exome array for association studies (see below).

### Phasing of genotype data for calling mosaic events

We phased the filtered genotypes mentioned above with the use of Eagle2 software^15^ which enabled us to conduct accurate long-range phasing.

### Calculation of BAF and LRR from genotype intensity

We computed B allele frequency (BAF) and log2 R ratio (LRR) values with the use of the BBJ genotype intensity data (X and Y)^43^. We modified previous methods used by Jacobs et al^1^ and Loh et al^10^ to fit the current data set. We computed LRR and BAF values in an array-basis in which all of the subjects genotyped in the same arrays were clustered together. Details are provided in the Supplementary Note.

### Filtering possible non-mosaic trisomy or monosomy events

We excluded chromosomes with mean LRR more than 0.2 or mean LRR less than −0.5 (possible trisomy and monosomy, respectively, Supplementary Note).

### Calling mosaic events with the use of BAF and LRR

We used the same method to call mosaic events as Loh et al^10^. This calling method is composed of the following steps: (1) filtering constitutional duplications; (2) evaluating phased BAF for variants on each chromosome using a parameterized hidden Markov model; (3) calling existence of events using a likelihood ratio test; (4) calling event boundaries; (5) calling copy number; (6) filtering remaining possible constitutional duplications; (7) estimating cell fraction of mosaic events. Details of each step are provided in the Supplementary Note.

### Analyses of associations between mCA detection rate and array batches or disease status at registry

We conducted logistic regression analyses to evaluate associations between mCA detection rate and 1) array batches or 2) disease status at registry. For array batches, we put mosaic detection as a dependent variable and age, sex, smoking and arrays as independent variables. For disease status at registry, we put disease presence as a dependent variable and presence of mosaic events, age, sex and genotyping array as independent variables.

### Comparison of mosaic frequency between BBJ and UKB

We co-analyzed BBJ mosaic calls with mosaic calls in UKB from 482,857 individuals^16^. We calculated frequencies of mosaic events subdivided by chromosome arm and copy number in both data sets. We assessed correlation of event frequencies in the two data sets using Spearman’s correlation coefficient.

### Associations between mosaic events and hematological traits

We extracted data from the BBJ of 13 hematological traits, namely, red blood cell count, hemoglobin, hematocrit, mean corpuscular volume, mean corpuscular hemoglobin, mean corpuscular cell hemoglobin, white blood cell count, neutrophil count, lymphocyte count, monocyte count, eosinophil count, basophil count and platelet count. Associations between 13 hematological quantitative traits and mosaic events were analyzed in a logistic regression model. The traits were standardized separately for males and females. We used disease status at registry, age, sex, smoking and genotyping arrays as covariates. We took this approach to avoid effects of disease status on the association results. We subdivided mosaic events by p vs. q arm for loss and CNN-LOH events. To reduce multiple testing burden, we restricted analyses to mosaic events with more than 20 carriers. As a result, 88 mosaic events were analyzed in association with 13 hematologic traits. Residuals of quantitative traits after regressing covariates were normalized and used as independent variables. The statistical significance threshold was set at *p*<0.05/88/13 (4.4×10^-5^).

### Relative coverage of the genome by mosaic events

We determined mosaic coverage as follows. We divided chromosomes into 0.1Mb bins and calculated the fraction of loss (respectively, CNN-LOH) events covering each bin to compute mosaic coverage. We compared mosaic coverage in BBJ and UKB using Pearson’s correlation coefficient.

### Genetic association studies

We excluded subjects showing high degree of kinship (more than 1^st^ degree relatives detected by plink^42^) with other subjects, leaving 173,599 subjects for genetic association studies. Among related pairs, we retained subjects having mosaic events. We also integrated the genotyping data used for calling mosaic events with genotyping data from additional variants on the Human Exome Array when available (Table S1) to maximize the number of variants used for imputation. (We did not integrate these data at the stage of calling mosaic events to avoid batch effects.) We phased the integrated data using Eagle2 software^15^. The phased genotypes were imputed using a reference panel containing 2,504 1000 Genomes Phase 3 samples and 1,037 Japanese high-depth (30x) whole-genome sequencing samples (Data set 1 of ref. ^29^) using Minimac3 software^44^. Variants imputed with *R*^2^>0.3 were used for the association studies. We filtered variants with minor allele count less than 5. Best guess data was used to conduct Fisher’s exact tests using plink software (plink --fisher --ci 0.95). We used GWAS array genotyping data if available to rescue rare variants not included in the reference panel, not well-imputed or with low allele frequency. As a result, 26.6 million variants were used for association studies.

We analyzed mosaic events in each chromosome as distinct phenotypes, treating loss, CNN-LOH and gain separately. In order to maximize power of this study to identify significant associations with CNN-LOH, we included to CNN-LOH unclassified events not involving whole chromosome but involving telomere with |LRR|<0.02. We further divided loss and CNN-LOH events in each chromosome into p-arm and q-arm events. We set a threshold of at least 20 event carriers to consider an event in genetic association studies. This led to a total of 88 copy number-chromosome pairs analyzed (Table S2). We tested each of these phenotypes for association with variants in *cis* (i.e., on the same chromosome and contained within a mosaic event) or *trans* (i.e., on any chromosome). For *cis* associations, we also conducted allelic imbalance analyses to assess whether one of the alleles at each variant was preferentially duplicated by mosaic CNN-LOH events. Details of each test are described in the Supplementary Note.

### Associations between mosaic events and mortality

The BBJ project has follow-up data to survey mortality and cause of death^40^. A total of 141,612 BBJ subjects who have one of 32 out of 47 diseases were prospectively followed up (∼10 years) after DNA collection. For subjects who died, further detailed surveillance was made to identify causes of death coded by ICD10 by accessing national vital registration system used for input survey of medical and social welfare at Ministry of Health, Labor, and Welfare Japanese Government.

We restricted subjects to those who were followed at least one year after registry and free from malignancy at blood collection. We found 86,546 subjects in the current study were included in the follow-up data for mortality. Among them, 16,812 deaths were recorded during the follow-up period. The average follow-up period was 7.6 years (median 8.3 years, s.d. 2.8 years).

We compared subjects with mosaic events (loss, CNN-LOH, or gain) at cell fraction ≥1% to subjects without mosaic events on any chromosomes.

Cox regression analysis was used for the analyses conditioning for age, age^2, sex, disease status, genotyping array and smoking. We used follow-up period as censoring factor. When we analyzed specific causes of death (e.g., non-hematopoietic malignancy), we used subjects whose deaths were not reported during follow-up as controls to use consistent control samples across analyses.

Associations between mortality (overall or specific causes) and presence of mosaic events (regardless of mosaic types) were analyzed as an initial evaluation. We analyzed overall mortality, hematopoietic malignancy mortality and non-hematopoietic malignancy mortality. We used significance thresholds based on Bonferroni’s correction (depending on tested mortality phenotypes).

After evaluating associations between mortality phenotypes and presence of any mosaic event in any chromosome, we searched for associations between specific mosaic event types and mortality. Mosaic types of any mosaic type (CNN-LOH, loss and gain), loss, CNN-LOH, gain were analyzed in each chromosome. CNN-LOH were subdivided into p and q arm events. We restricted to mosaic events present in at least 20 carriers in the initial data set (Table S1). A total of 120 specific mosaic types were analyzed. In the analysis of associations of specific mosaic types, we divided subjects based on age, sex and smoking and computed associations with the use of Cochran-Mantel-Haenszel method to avoid inflation of statistics arising from limited number of subjects carrying mosaic events. We set a significance level based on Bonferroni’s correction (p<0.00042, 0.05/120).

We also analyzed cardiovascular mortality (defined as ischemic heart diseases and ischemic stroke) as previous studies have reported associations between mosaic point mutations and cardiovascular outcomes.

### Definition of cancers

We categorized causes of death to decrease multiple testing burden. Hematopoietic malignancy was defined by ICD10 codes of C81-C96 and D45, 46 and D47. Leukemic diseases were defined by ICD10 codes of C91-C96, D45 and D46. Malignant lymphoma was defined by ICD10 codes of C81-C88. Multiple myeloma was defined by C90. Cancers were defined as ICD10 codes starting with “C” together with hematopoietic malignancies defined not starting with “C”. We did not regard other ICD10 codes starting with “D” as cancer since most of them are benign tumors.

### Associations between multiple mosaic events and mortality

We extended the mortality analyses to investigate the effect of multiple mosaic events within a single individual. We limited analyses to subjects with at most three mosaic events. We divided subjects into three groups: (1) subjects without mosaic events, (2) subjects with a single mosaic event, (3) subjects with multiple mosaic events (2 or 3 mosaic events in different chromosomes). We analyzed an association of presence of multiple mosaic events with leukemia mortality in comparison with presence of single mosaic event. The analyses were conducted by conditioning on age, sex, disease status, genotyping array and smoking.

### Associations between cell fraction of mosaic events and mortality

We also extended the mortality analyses to investigate the effect of mosaic cell fraction. For subjects with multiple mosaic events, we took the highest cell fraction. We divided subjects into categories according to cell fraction of mosaic events and analyzed associations between cell fractions and outcomes with which presence of mosaic events were significantly associated.

## Acknowledgments

This research was conducted using the UK Biobank Resource under Application #19808.

P.-R.L. was supported by NIH grant DP2 ES030554, a Burroughs Wellcome Fund Career Award at the Scientific Interfaces, the Next Generation Fund at the Broad Institute of MIT and Harvard, and a Glenn Foundation for Medical Research and AFAR Grants for Junior Faculty award.

## References

1. Jacobs, K. B. et al. Detectable clonal mosaicism and its relationship to aging and cancer. Nat. Genet. 44, 651–658, doi:10.1038/ng.2270 (2012).

2. Laurie, C. C. et al. Detectable clonal mosaicism from birth to old age and its relationship to cancer. Nat. Genet. 44, 642–650, doi:10.1038/ng.2271 (2012).

3. Genovese, G. et al. Clonal hematopoiesis and blood-cancer risk inferred from blood DNA sequence. N. Engl. J. Med. 371, 2477–2487, doi:10.1056/NEJMoa1409405 (2014).

4. Jaiswal, S. et al. Age-related clonal hematopoiesis associated with adverse outcomes. N. Engl. J. Med. 371, 2488–2498, doi:10.1056/NEJMoa1408617 (2014).

5. Machiela, M. J. et al. Characterization of large structural genetic mosaicism in human autosomes. Am. J. Hum. Genet. 96, 487–497, doi:10.1016/j.ajhg.2015.01.011 (2015).

6. Vattathil, S. & Scheet, P. Extensive Hidden Genomic Mosaicism Revealed in Normal Tissue. Am. J. Hum. Genet. 98, 571–578, doi:10.1016/j.ajhg.2016.02.003 (2016).

7. Young, A. L., Challen, G. A., Birmann, B. M. & Druley, T. E. Clonal haematopoiesis harbouring AML-associated mutations is ubiquitous in healthy adults. Nature communications 7, 12484, doi:10.1038/ncomms12484 (2016).

8. Forsberg, L. A., Gisselsson, D. & Dumanski, J. P. Mosaicism in health and disease - clones picking up speed. Nat Rev Genet 18, 128–142, doi:10.1038/nrg.2016.145 (2017).

9. Abelson, S. et al. Prediction of acute myeloid leukaemia risk in healthy individuals. Nature 559, 400–404, doi:10.1038/s41586-018-0317-6 (2018).

10. Loh, P. R. et al. Insights into clonal haematopoiesis from 8,342 mosaic chromosomal alterations. Nature 559, 350–355, doi:10.1038/s41586-018-0321-x (2018).

11. Zhou, W. et al. Mosaic loss of chromosome Y is associated with common variation near TCL1A. Nat. Genet. 48, 563–568, doi:10.1038/ng.3545 (2016).

12. Hinds, D. A. et al. Germ line variants predispose to both JAK2 V617F clonal hematopoiesis and myeloproliferative neoplasms. Blood 128, 1121–1128, doi:10.1182/blood-2015-06-652941 (2016).

13. Wright, D. J. et al. Genetic variants associated with mosaic Y chromosome loss highlight cell cycle genes and overlap with cancer susceptibility. Nat. Genet. 49, 674–679, doi:10.1038/ng.3821 (2017).

14. Nagai, A. et al. Overview of the BioBank Japan Project: Study design and profile. J. Epidemiol. 27, S2–S8, doi:10.1016/j.je.2016.12.005 (2017).

15. Loh, P. R., Palamara, P. F. & Price, A. L. Fast and accurate long-range phasing in a UK Biobank cohort. Nat. Genet. 48, 811–816, doi:10.1038/ng.3571 (2016).

16. Loh, P.-R., Genovese, G. & McCarroll, S. A. Monogenic and polygenic inherited causes of clonal hematopoiesis. bioRxiv (2019).

17. Sudlow, C. et al. UK biobank: an open access resource for identifying the causes of a wide range of complex diseases of middle and old age. PLoS Med. 12, e1001779, doi:10.1371/journal.pmed.1001779 (2015).

18. Bycroft, C. et al. The UK Biobank resource with deep phenotyping and genomic data. Nature 562, 203–209, doi:10.1038/s41586-018-0579-z (2018).

19. Iwanaga, M., Watanabe, T. & Yamaguchi, K. Adult T-cell leukemia: a review of epidemiological evidence. Front Microbiol 3, 322, doi:10.3389/fmicb.2012.00322 (2012).

20. Tamura, K. et al. Chronic lymphocytic leukemia (CLL) is rare, but the proportion of T-CLL is high in Japan. Eur. J. Haematol. 67, 152–157 (2001).

21. Li, Y., Wang, Y., Wang, Z., Yi, D. & Ma, S. Racial differences in three major NHL subtypes: descriptive epidemiology. Cancer Epidemiol. 39, 8–13, doi:10.1016/j.canep.2014.12.001 (2015).

22. Landau, D. A. et al. Mutations driving CLL and their evolution in progression and relapse. Nature 526, 525–530, doi:10.1038/nature15395 (2015).

23. Puente, X. S. et al. Non-coding recurrent mutations in chronic lymphocytic leukaemia. Nature 526, 519–524, doi:10.1038/nature14666 (2015).

24. Iwai, M. et al. Expression and methylation status of the FHIT gene in acute myeloid leukemia and myelodysplastic syndrome. Leukemia 19, 1367–1375, doi:10.1038/sj.leu.2403805 (2005).

25. Schmitz, R. et al. TNFAIP3 (A20) is a tumor suppressor gene in Hodgkin lymphoma and primary mediastinal B cell lymphoma. J. Exp. Med. 206, 981–989, doi:10.1084/jem.20090528 (2009).

26. Liu, Y. et al. The genomic landscape of pediatric and young adult T-lineage acute lymphoblastic leukemia. Nat. Genet. 49, 1211–1218, doi:10.1038/ng.3909 (2017).

27. Zink, F. et al. Clonal hematopoiesis, with and without candidate driver mutations, is common in the elderly. Blood 130, 742–752, doi:10.1182/blood-2017-02-769869 (2017).

28. Genomes Project, C. et al. A global reference for human genetic variation. Nature 526, 68–74, doi:10.1038/nature15393 (2015).

29. Okada, Y. et al. Deep whole-genome sequencing reveals recent selection signatures linked to evolution and disease risk of Japanese. Nature communications 9, 1631, doi:10.1038/s41467-018-03274-0 (2018).

30. Kilpivaara, O. et al. A germline JAK2 SNP is associated with predisposition to the development of JAK2(V617F)-positive myeloproliferative neoplasms. Nat. Genet. 41, 455–459, doi:10.1038/ng.342 (2009).

31. Jones, A. V. et al. JAK2 haplotype is a major risk factor for the development of myeloproliferative neoplasms. Nat. Genet. 41, 446–449, doi:10.1038/ng.334 (2009).

32. Olcaydu, D. et al. A common JAK2 haplotype confers susceptibility to myeloproliferative neoplasms. Nat. Genet. 41, 450–454, doi:10.1038/ng.341 (2009).

33. Lee, J. H. & Paull, T. T. ATM activation by DNA double-strand breaks through the Mre11-Rad50-Nbs1 complex. Science 308, 551–554, doi:10.1126/science.1108297 (2005).

34. Dewez, M. et al. The conserved Wobble uridine tRNA thiolase Ctu1-Ctu2 is required to maintain genome integrity. Proc. Natl. Acad. Sci. U. S. A. 105, 5459–5464, doi:10.1073/pnas.0709404105 (2008).

35. Adzhubei, I., Jordan, D. M. & Sunyaev, S. R. Predicting functional effect of human missense mutations using PolyPhen-2. Curr Protoc Hum Genet Chapter 7, Unit7 20, doi:10.1002/0471142905.hg0720s76 (2013).

36. Sim, N. L. et al. SIFT web server: predicting effects of amino acid substitutions on proteins. Nucleic Acids Res. 40, W452–457, doi:10.1093/nar/gks539 (2012).

37. DeAntoni, A., Sala, V. & Musacchio, A. Explaining the oligomerization properties of the spindle assembly checkpoint protein Mad2. Philos. Trans. R. Soc. Lond. B Biol. Sci. 360, 637-647, discussion 447-638, doi:10.1098/rstb.2004.1618 (2005).

38. Karczewski, K. J. et al. Variation across 141,456 human exomes and genomes reveals the spectrum of loss-of-function intolerance across human protein-coding genes. bioRxiv (2019).

39. Jaiswal, S. et al. Clonal Hematopoiesis and Risk of Atherosclerotic Cardiovascular Disease. N. Engl. J. Med. 377, 111–121, doi:10.1056/NEJMoa1701719 (2017).

40. Hirata, M. et al. Overview of BioBank Japan follow-up data in 32 diseases. J. Epidemiol. 27, S22–S28, doi:10.1016/j.je.2016.12.006 (2017).

41. Consortium, G. P. A map of human genome variation from population-scale sequencing. Nature 467, 1061–1073, doi:nature09534[pii]10.1038/nature09534 (2010).

42. Purcell, S. et al. PLINK: a tool set for whole-genome association and population-based linkage analyses. Am. J. Hum. Genet. 81, 559–575, doi:S0002-9297(07)61352-4[pii]10.1086/519795 (2007).

43. Staaf, J. et al. Normalization of Illumina Infinium whole-genome SNP data improves copy number estimates and allelic intensity ratios. BMC Bioinformatics 9, 409, doi:10.1186/1471-2105-9-409 (2008).

44. Das, S. et al. Next-generation genotype imputation service and methods. Nat. Genet. 48, 1284–1287, doi:10.1038/ng.3656 (2016).

