## Supplementary Materials for "The genomic landscape of clonal hematopoiesis in Japan"

Supplementary Method page 3-10

Supplementary Note page 11-15

Fig. S1-S55 page 16-73

Tables S1-S28 page 74-108

Supplementary Reference page 109

### Supplementary Methods

**Calculation of BAF and LRR from genotype intensity**

1. Computation of cluster median of X and Y signal intensities.

We calculated median of X and Y intensities in each genotype. If a cluster contained calls less than 10, we set its median to missing.

2. Affine-normalization and correction by GC and CpG contents.

We took a similar approach to Jacobs et al[1], and our method is the same as Loh et al[10] regarding this calculation. A pair of multiple variate linear regression was carried out to correct effects of GC and CpG contents. Median of centers (X, Y) were set as two dependent variables to be expected (X, Y) values and modeled with the use of X and Y values in each subject of the genotype, GC and CpG contents in 9 windows of 50, 0.1k, 0.5k, 1k, 10k, 50k, 100k, 250k and 1Mbp at the center of $SNP_m$ as covariates as follows.

$$X_{m,i,exp} = e_{x,i} + X_{m,i}\alpha_{x,i} + Y_{m,i}\alpha_{y,i} + \sum_{k=1}^{9}\sum_{p=1}^{2}[(f_{m,k}^{GC})^p \cdot \alpha_{i,k,p}^{GC} + (f_{m,k}^{CpG})^p \cdot \alpha_{i,k,p}^{CpG}\;]$$

$$Y_{m,i,exp} = e_{y,i} + X_{m,i}\beta_{x,i} + Y_{m,i}\beta_{y,i} + \sum_{k=1}^{9}\sum_{p=1}^{2}[(f_{m,k}^{GC})^p \cdot \beta_{i,k,p}^{GC} + (f_{m,k}^{CpG})^p \cdot \beta_{i,k,p}^{CpG}\;]$$

where $X_{m,i,exp}$ and $Y_{m,i,exp}$ are median of X and Y intensity in a cluster of a genotype of i-th individual in m-th variant, respectively, $X_{m,i}$ and $Y_{m,i}$ are X and Y values in i-th individual for m-th variant, respectively, $\alpha_{x,i}$ , $\beta_{y,i}$ are effect sizes of calculated X and Y for i-th individual, respectively, $f_{m,k}^{GC}$ or and $f_{m,k}^{CpG}$ are fraction of GC and CpG in k-th window, $\alpha_{i,k,p}^{CpG}$ $and$ $\beta_{i,k,p}^{GC}$, are effect size of GC and CpG in i-th individual for k-th window, respectively and $e_{x,i}$ and $e_{y,i}$are the error terms of X and Y for i-th individual, respectively.

The multi-variate linear regression analyses were conducted per individual (~179k sets of models), assuming fixed effects of X, Y intensities, GC and CpG waves across variants.

The GC content was computed with the use of bedtools on the hg19 reference. The CpG content was calculated with the use of EpiGRAPH CpG annotation[45]. Corrected X, Y values were calculated by expected X and Y values minus residuals of X and Y.

3.Computation of means of corrected X and Y values.

Means of corrected X and Y calculated above in the three cluster centers for each genotype were computed.

4.Transformation of corrected X and Y intensities to θ and R values.

We transformed corrected X and Y to θ and R as follows;

$$\theta = \frac{2}{\pi} \cdot \arctan(\frac{Y}{X})$$

$$log_2 R = X + Y$$

This follows the method by Staaf et al[43] and differs from the method by Loh et al[10].

5. Transformation of θ and R to LRR and BAF values.

We computed cluster centers of each genotype to obtain θ and R in three cluster centers. We conducted linear interpolation between the cluster centers to estimate expected $\log_2 R$ based on θ in each individual. If cluster centers of homozygotes were missing, we used reflection of the cluster center in the opposite homozygote genotype across the vertical line on the center of heterozygote genotype.

6. Mean shift of LRR.

We noticed that LRR in our dataset had slightly downward bias. To avoid noise in mosaic calls due to this bias, we shifted LRR values in each genotype for each variant to have mean 0 (mean shift).

**Calling mosaic events with the use of BAF and LRR**

1.Filtering constitutional duplications

The 25 states corresponding to phased BAF deviation (from -0.24 to 0.24 with interval of 0.02) were used in HMM. We assume events in the state revealed mean BAF deviation equal to state value (-0.24 – 0.24 with interval of 0.02) with empirical standard deviation computed across genotyping results. We capped z-score of 4. Transition probability was determined 0.003 from zero to non-zero state, 0.001 from a state to its negative value (phase switch error). Regions to mask were selected by computing maximum likelihood (Viterbi path) and examining contiguous non-zero states. We masked regions of likely constitutional duplications with <2Mb with |ΔBAF|>0.1 and LRR>0.1 and their 2Mb nearby regions.

2.Parameterized hidden Markov model for event detection

We used a family of 3 states HMMs parametrized by deviation of BAF, θ, namely, {-θ,0, θ} taking phase switch errors into account with transition matrix slightly different from 1st step (from±θ to 0, 0.0003, from 0 to ±θ, 0.004 * 0.0003 and switch error of 0.001). For acrocentric chromosomes (no p-arm genotypes), starting probability of non-zero state was decreased by a factor of 0.2. Like the 1st step, we assume events in the state revealed mean BAF deviation equal to state value with empirical standard deviation computed across genotyping results. We capped z-score of 2.

3.Calling existence of an event: likelihood ratio test statistics

Based on a sequence of phased BAF deviation (ΔBAF) on each chromosome, we can analyze whether the sequence of observed ΔBAF can be explained by presence of mosaic events with the states defined above. We compute the likelihood ratio statistics with the use of a family of HMM paths parameterized by θ and modeled a total probability observing the sequence ofΔBAF. Likelihood ratio for θ can be modeled by $\mathrm{f}(\Delta\mathrm{BAF}) = \frac{\mathrm{L}(0 \mid \Delta\mathrm{BAF})}{sup_{\theta}\{L(\theta \mid \Delta BAF)\}}$

0 of θ indicates no mosaic events. We discretized θ to run from 0.001 to 0.25 in 50 multiplicative steps in practice.

4.Calling event boundaries

Boundaries of called events were determined based on the consensus of mosaic calls in five samples taken from the posterior of the HMM using the likelihood-maximizing value for θ.

5. Calling copy number

Since we noticed that the steepness of slopes in BAF-LRR space to distinguish possible gains and losses from CNN-LOH in the current data set were quite different from the previous study (this is not limited to raw or transformed BAF and LRR values in the current study), we did not use the previously-defined threshold of slopes. We defined 1.3 and -1.05 to distinguish gains and losses from CNN-LOH, respectively. We called copy number only if the most likely call was at least 10 times more likely than the next-most likely call (~90% confidence).

6. Filtering possible constitutional duplications after calling mosaic events.
We excluded possible constitutional duplications with length >10 Mb with LRR > 0.35 or LRR >0.2 and |ΔBAF|>0.16. We also filtered events with length <10Mb with LRR >0.2 or LRR>0.1 and |ΔBAF|>0.1. These threshold was defined by the previous study[10].

We further excluded events classified as gain or unknown and satisfying any of the following criteria: (1) less than 5Mbp length and contained in segmental duplications reported in 1000 genome projects with extension of 0.1Mbp in both directions (2) length <5Mb with LRR>0.13, or (3) length<2Mbp and LRR>0.02.

We excluded events with LRR>-0.1 and observed heterozygosity within events less than one third of expected heterozygosity.

7. Estimating fractions of cells carrying mosaic events.

Cell fraction with mosaic events was calculated by the method previously developed[1] as follows.

muDiff = 2 x 0.01 x BAF

$AF_{loss}$ = 2*muDiff / (1+muDiff)

$AF_{CNN}$ = muDiff

$AF_{gain}$ = 2*muDiff / (1-muDiff)

where $AF_{mosaic\ type}$ indicates cell fraction of the corresponding mosaic types.

**Exclusion of samples of possible contamination.**
We excluded three samples with mosaic events in more than 7 chromosomes among which they did not carry loss or gain events but many unknown events strongly suggesting contamination.

**Exclusion of events of possible non-mosaic whole chromosomal trisomy.**
We additionally filtered mosaic events on chromosomes with chromosome-wide mean LRR more than 0.175.

**Co-occurrence of mosaic events.**

We assessed co-occurrence of classified mosaic events in single individual. We excluded co-occurrence of mosaic events in the same chromosome (for instance 1. multiple LOSS in the same chromosome, 2. LOSS in chromosome 17p and GAIN in chromosome 17q). We evaluated odds ratio (OR) of co-occurrence by Fisher's exact test based on 2x2 tables (rows and columns corresponding to individuals with and without the two mosaic events). We excluded subjects having mosaic events more than 5.

**Analysis of focal deletions**

We focused on mosaic deletion events in each chromosome. Focal mosaic events are determined based on plots of mosaic coverage in the two populations. We excluded chromosome 19 because deletions on chr19 are rare in both populations. We also evaluated importance of genes by taking numbers of gene involved in loss events into account. We counted the number of genes involved in each loss event and defined a score of each loss event as one divided by the number (when a loss event contained only one gene, the gene received a score of one). We summed up all scores across all of loss events in each gene. To pick up genes frequently involved with focal deletions only in Japanese, we picked up genes covered by at least 5% of loss events in a chromosome, more than 10 times of scores than UKB, and scores more than 0.5.

**Cis-association**

Cis-1. Evaluation of cis-variants' associations with mosaic events

We restricted variants with minor allele count more than five to use in genetic studies. Since the previous UKB study strongly suggested that associated variants are associated with mosaic events in cis-acting manner, e.g., variants in p-arm are associated with p-arm mosaic events, we conducted cis-associations at first. For instance, when we analyzed chr 2p CNN-LOH, we extracted variants in chromosome 2p with MAC more than five and rsq more than 0.3. We excluded subjects not carrying the mosaic event being studied carrying any other mosaic event on the chromosome from each analysis. While the variants in each chromosome or chromosome arm are not mutually independent, we set a stringent cut-off of significance of $p<5x10^{-9}$. Since the previous UKB study reported associations between CNN-LOH events that extended to telomeres and did not span whole chromosomes and variants contained within these events, we divided chromosomes into bins with 1Mbp and restricted cases to carriers of events reaching at least the border of the 1Mbp window and extending to telomeres (but not spanning whole chromosomes) when performing association studies for CNN-LOH events. For association analyses for CNN-LOH, we allowed to include unclassified events as long as the events satisfied the conditions mentioned above to maximize the statistical power.

Cis-2. Allelic imbalance study in cis associations for CNN-LOH.

We conducted allelic imbalance analyses in mosaic events[10] (analogous to allele-specific expression in gene expression) to assess whether one of the alleles at each variant was preferentially duplicated by mosaic CNN-LOH events. We took advantage of phase information of each individual and assessed allelic imbalance in each site by extracting subjects heterozygous for risk variants and carrying mosaic events. P-values of allelic imbalance were calculated based on a binomial test under the null hypothesis that the allelic shift of mosaic events at heterozygote sites is at random.

Cis-3. Evaluation of variants reported in the previous UKB study.

We evaluated whether variants which were significantly associated with mosaic events in the previous UKB study were present in our data and associated with mosaic events. We extracted from the current results a total of six variants, namely, three variants in *MPL* (rs144279563, rs182971382 and rs369156948 (nonsense mutation)), rs118137427 in *FRA10B*, rs532198118 in *ATM*, and rs182643535, tagging 70kb deletion of *TM2D3*/*TARSL2* region. We also analyzed whether these variants were included in the Japanese WGS used in the reference panel in the current study.

**Trans-association**

Trans-1.Genome-wide search for associated variants

After evaluating cis-associations, we conducted trans-associations (between chromosome and arm-specific mosaic events and variants outside the chromosome and arm of the mosaic events). We conducted association analyses by Fisher's exact test. We set a very conservative threshold of significance at p-values less than $5.0x10^{-11}$ (considering 88 tested mosaic events and 26.6 million variants).

Trans-2. Candidate analyses of associations between mosaic events and variants associated with MPN, CLL or mLOY.

We combined the 86 variants in the previous study[10] which were associated with MPN, CLL or mLOY in the European population with 4 variants we recently found to be associated with mLOY as the 2nd hit in the known genes (Terao et al, submitted). We excluded variants in *TERT*, *DLK1*, *JAK2* and *TCL1A* since these genes were shown to be significantly associated with mosaic events in Table 1. As a result, we analyzed a total of 63 variants. We evaluated whether variants showed associations with loss, CNN-LOH, or gain in any chromosomes or any mosaic types in any chromosomes. We set a significance level using Bonferroni's correction. When we found a variant satisfying the significance level, we further analyzed detailed mosaic types and detailed chromosomes (we tested an additional 118 associations with mosaic types with more than 20 carriers).

**Functional analyses of gene expression in significant variants**

We regarded the *NBN* and *CTU2* variants as likely to be causal since they showed stop-gain and amino acid alteration predicted as deleterious by multiple prediction methods, respectively. We conducted functional analyses for significant variants whose functions were not easily interpreted, namely, the *MRE11* and *MPL* variants. We evaluated alteration of gene expression in risk alleles in contrast to reference alleles by the following methods.

**Vector construction and luciferase reporter assay**

The section containing intron in *MRE11* gene with polymorphism and upstream region of *MPL* gene with polymorphism were generated by synthetic oligonucleotides. Annealed oligonucleotides were digested by NheI and HindIII, and linked into the vector pGL4.24 minP vector and pGL4.11 basic vector (Promega, Madison, WI), respectively. As the result, pGL4minP-G and pGL4minP-A which encodes the C/T variant located on 94160189 in *MRE11* gene intron (chr11), and pGL4basic-G and pGL4basic-A wihich encodes the G/A variant located on 43799207 in upstream of *MPL* gene (chr1) were constructed. These vectors were transformed into Escherichia coli strain DH5α and recovered using the EndoFree Plasmid Maxi Kit (Qiagen, West Sussex, UK). The presences of polymorphisms were verified by sequencing. The recombinant plasmids were used for luciferase assay with pGL4minP and pGL4basic as mock vectors. The Jurkat E6.1 cell line and THP-1 cell line (purchased from ATCC) were maintained in RPMI1640 with 10%FBS at 37°C in a 5% $CO_2$ incubator. Transfection of Jurkat cells with the reporter vectors carrying each variants was carried out using Neon® Transfection System (Invitrogen, Carlsbad, CA). 0.2 Million cells were resuspended in 100 µl of electroporation buffer R that contained 0.5µg of pGL4.74 (Renilla luciferase-TK control reporter vector, Promega) and 2.5 µg of each test vector with polymorphism or empty vector for control. The procedure was conducted according to the manufacturer’s protocol with electroporation options recommended for each cell line (three 10 ms, 1350 mV pulses for Jurkat E6.1 and THP-1). Luciferase activity was measured 24 h after transfection using Dual-Luciferase Reporter Assay System (Promega) according to the manufacturer’s protocol. Luminescence was detected by TriStar LB941 (Berthold, Bad Wildbad, Germany). Information of oligo nucleotide sequence used for the current study was provided in Table S24.

**Electrophoretic mobility shift assay (EMSA)**

The following protocol has been used to make nuclear extracts from Jurkat E6-1.1x 107 Jurkat E6-1, washed with PBS and used for nuclear extraction as described previously[46].Two double-stranded 51-nucleotide biotin-labeled DNA probes were prepared by annealing (Table S24). EMSA experiments were carried out using the Lightshift Chemiluminescent EMSA kit (Thermo Fisher), as recommended by the supplier. In brief, 2 µL Binding buffer was mixed with 6.4 µg nuclear extract, then 50 fmol biotin-labeled probe was added, and hybridization was carried out for 20 min at room temperature. The mixtures were then loaded into a 6% Polyacrylamide gel, separated by electrophoresis at 4°C, and transferred onto a nylon membrane. As competitors, non-labeled oligo nucleotides were incubated with nuclear extracts before adding the labeled probe.

#### Supplementary Note.

##### Filtering possible monosomy and trisomy based on BAF and LRR.

We noticed that long-range phasing and phasing itself did not always produce a clear sequence of deviation of BAF across chromosomal positions in subjects having possible trisomy. This is compatible with previous findings of trisomy 21 composed of one paternal and two maternal chromosomes without a duplicated chromosome. In addition to excess departure of BAF and LRR in a single chromosome, this type of signal characterizes the existence of a trisomy originating from three different chromosomes.

##### Inevitable development of mosaic events in the elderly.

The trend toward inevitability in the elderly has not previously been observed for mosaic of point mutations[7,27]; however, previous studies of large chromosomal alterations have detected autosomal mCAs in at most 13% of very elderly individuals[1,2,5,6,10]. Here, the observation of mCA rates reaching 40% was driven by the combination of sensitive statistical methods[10], low noise in measuring allele-specific copy number, and representation of very elderly individuals in the BBJ cohort.

##### Age and sex associations with mCA

The BBJ data revealed that chr15 gain strongly skewed towards males and the elderly, which was also observed in the UKB[10]. Chr20q loss also showed the same pattern in the both populations. In the BBJ data, we did not observe any mosaic types with female dominance which was different from that in the UKB.

##### Co-occurrence of mosaic events

We observed 30 combinations of mosaics which significantly co-occur in the BBJ (Fig. S54 and Table S25). Five out of the 30 combinations, including a combination of chromosome 3 gain and chromosome 18 gain with the strongest association, were also reported in the UKB. The common combinations include a combination of chromosome 12 gain and chromosome 13q loss, both of which were associated with CLL. These suggest common and population-specific mechanisms underlying co-occurrence of mosaic events.

##### Cell fractions in multiple mosaic events in subjects.

We extracted subjects having two mosaic events in different chromosomes. We compared cell fractions of two mosaic events in subjects (Fig. S23). At least 54.5% subjects were estimated to have different cell fractions, suggesting multiple clones with mosaic events. We further analyzed whether specific combinations of chromosomes showed skewness of lack of difference in cell fractions, but we did not find specific chromosomal combinations. These results suggest that mosaic events do not usually occur at the same time in a single cell in spite of findings of specific mosaic combinations frequently observed in a single subject.

**Associations between quantitative hematologic traits and mosaic events.**

We found significant associations of multiple loss and gain events with hematologic traits (Table S9). Since no CNN-LOH reached statistical significance in spite of the CNN-LOH events accounting for most mosaic events, this results suggests larger influences of loss or gain, which decrease or increase copy number, on hematopoiesis.

**Consistent breakpoint and coverage of CNN-LOH between BBJ and UKB.**

CNN-LOH events, which typically extend from an interstitial breakpoint to a telomere, exhibited similar quantitative distributions across the autosomes in BBJ and UKB (Fig. 3c and Fig. S48). Correlations of CNN-LOH chromosomal coverage between BBJ and UKB ranged from 0.80–1.00 (Table S13), suggesting cross-population consistency in the mutational process (mitotic recombination) and selective pressures that lead to CNN-LOH events in clonal hematopoiesis.

**Genes frequently involved in focal deletions in Japanese but not in UK population.**

We found Japanese-specific focal deletions (Tables S14-S15). A total of 37 regions (defined by 1Mbp margin of each region) across 15 chromosomes were identified. *TNFAIP3* showed the highest scores (see Methods) among Japanese specific genes. Other genes which draw our attention were *FHIT* in chromosome 3 (which encompasses the fragile site *FRA3B*) and *VEGFC* in chromosome 4. Note that in this analysis, we focus on absolute coverage of genes by focal deletions and coverage was not scaled (different from S25-46).

**Focal deletions of TCR genes**

Since focal deletions at the TCR alpha locus (*TRA*) in chromosome 14 were frequently found in the BBJ, we analyzed whether this trend was consistent in *TRB* in chromosome 7. We observed more frequent focal deletions in *TRB* in the BBJ than the UKB (Fig. S7 and S31).

Since genetic recombination in TCR occurs in all of lymphocytes, focal deletions in TCR genes may not necessarily indicate clonal expansion (possibly reflecting common recombination of adjacent regions among TCR genes in majority of lymphocytes). To address this point, we analyzed overlap of focal deletions in *TRA* and *TRB*. We found significant co-occurrence of these two deletions (OR 305, $p=3.5 \times 10^{-11}$), indicating clonal expansion.

We confirmed that these two events were not enriched for subjects with cancer at registry.

**Significant cis loci associated with mCA in the BBJ.**

We identified five new loci showing cis-associations (with presence of mCA), namely, *NBN*, *MRE11*, *CTU2*, *NEDD8*/*TINF2* and *DLK1*. We also found *TCL1A* showing significant cis association with allele selection in mCA. The associations of *MPL* and *JAK2* were replicated in the BBJ. All of the eight associations were observed in CNN-LOH, compatible with the previous study[10] and in line with maximizing the number of events of CNN-LOH by including fraction of unclassified events.

*DLK1* encodes a noncanonical NOTCH ligand and is an imprinted gene[47] associated with both hematopoietic[48] and non-hematopoietic malignancies[49]. *NEDD8* encodes a ubiquitin-like protein also reported in the context of both hematopoietic and non-hematopoietic malignancy[50] and a therapeutic target for AML[51]. *TINF2* encodes a protein of a member of telosome complex which protects telomere. A mutation of TINF2 is known to cause congentical bone marrow failure[52]. *TCL1A* (T-cell leukemia/lymphoma 1A) is a susceptibility gene to mosaic of chromosome Y and hematopoietic and somatic cancer[13] and encodes a protein which expresses in fetal tissues and early stage of lymphocytes, interacts with many partners including ATM and is involved in multiple signaling including NFkB[53].

The association between a *JAK2* variant and presence of clonal expansion of V617F was previously reported. We found consistent associations between chr9p CNN-LOH and variants associated with *JAK2* V617F, indicating that chr9p CNN-LOH involving *JAK2* seem equivalent to *JAK2* V617F and that *JAK2* V617F involve other regions outside *JAK2* by duplicating a mutated haplotype.

A total of four associations (three cis and one trans) were identified in chr14q CNN-LOH. This is compatible with the largest number of chr14q CNN-LOH among autosomes and may suggest high susceptibility of mCA in chr 14 and high heritability of chr14q CNN-LOH.

**An enhanced strong association of NBN**

At *NBN*, we observed a very penetrating association between the rare stop gain variant rs756831345 and chr8q CNN-LOH (OR=240 (129-472), $p=1.1\times10^{-22}$) when we called mosaic events at FDR of 0.025 and restricted to events confidently called and not include unclassified events (possible CNN-LOH) in the association study. This restriction apparently reduces false-positive signals (at the cost of possible weaker p-values). This trend of higher OR was observed in the other two novel rare variant's associations (*MRE11*: OR=52 (22-124) and *CTU2*: OR=43 (25-73)), but quite prominent in the association of chr8q suggesting that the *NBN* association is associated with mosaic events with higher cell fraction.

**Pleiotropic associations of TERT**

At *TERT*, a SNP previously associated with mosaic *JAK2* V617F mutation[12] associated with 14q CNN-LOH events ($p=1.5\times10^{-22}$, OR =1.27 (1.21-1.33); Table 1 and Fig
. S51). Risk alleles at *TERT* have previously been observed to associate with clonal hematopoiesis involving a variety of mosaic mutations[10,12,27]; consistent with this finding, we observed that the *TERT* SNP also exhibited nominal association (after Bonferroni correction) with mosaic 20q- events ($p$=1.8 $\times10^{-7}$) and gains on chromosome 15 ($p$=0.00011) (curiously with the opposite allele increasing risk of +15). Candidate variant association tests of previously-reported risk variants for clonal hematopoiesis and hematological malignancies revealed additional, weaker *trans* associations with mCAs that will likely reach genome-wide significance in future studies (Table S26-S28).

**Gene expression changes of the variants in *MPL* and *MRE11* indicated by the functional analyses**

The luciferase assay revealed that risk variants of 1:43799207 and 11:94160189 at the *MPL* and *MRE11*, respectively, were associated with slight decrease and increase in gene expressions, respectively (Fig. S55). 1:43799207 is a lead SNP in the *MPL* region. EMSA assay suggested binding of transcription factor with 11:94160189 at *MRE11* (Fig. S55). However, previous eQTL studies revealed common variants with much stronger effects on *MRE11*. For instance, the GTEx project reported rs509744 with minor allele frequency of 0.30 associated with alteration of *MRE11* in whole blood ($p=1.0 \times 10^{-30}$ for 369 subjects).

Considering the high penetrance of the associations between the risk variants and CNN-LOH, the causal relationship between alteration of the gene expression via these rare variants and mechanism underlying CNN-LOH is inconclusive. Alternatively, it may be more likely that rare causal variants introducing functional impairment of proteins are present but not well imputed in the current study.

Fig. S1 A landscape of mosaic events in chromosome 1.

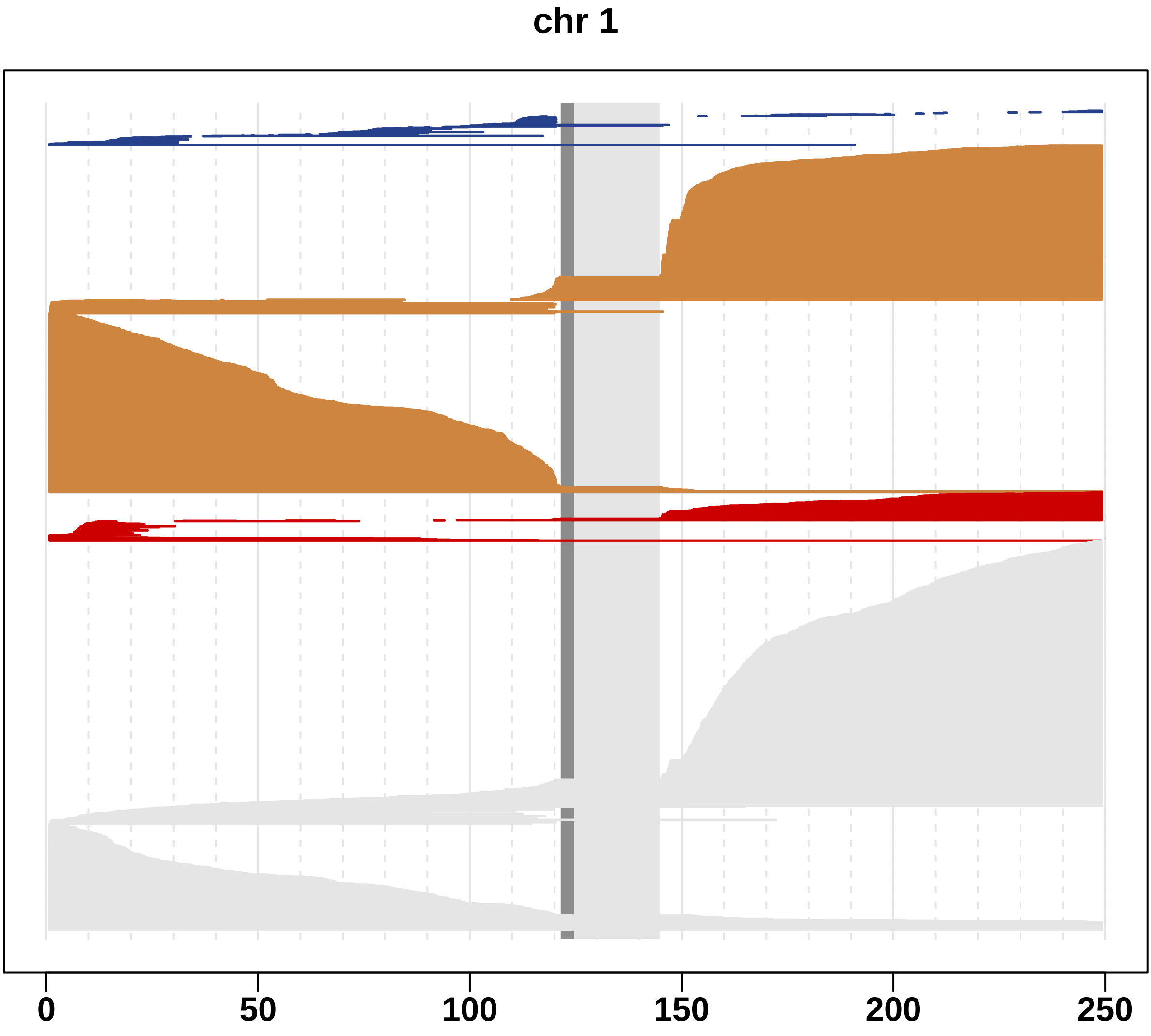

Fig. S2 A landscape of mosaic events in chromosome 2.

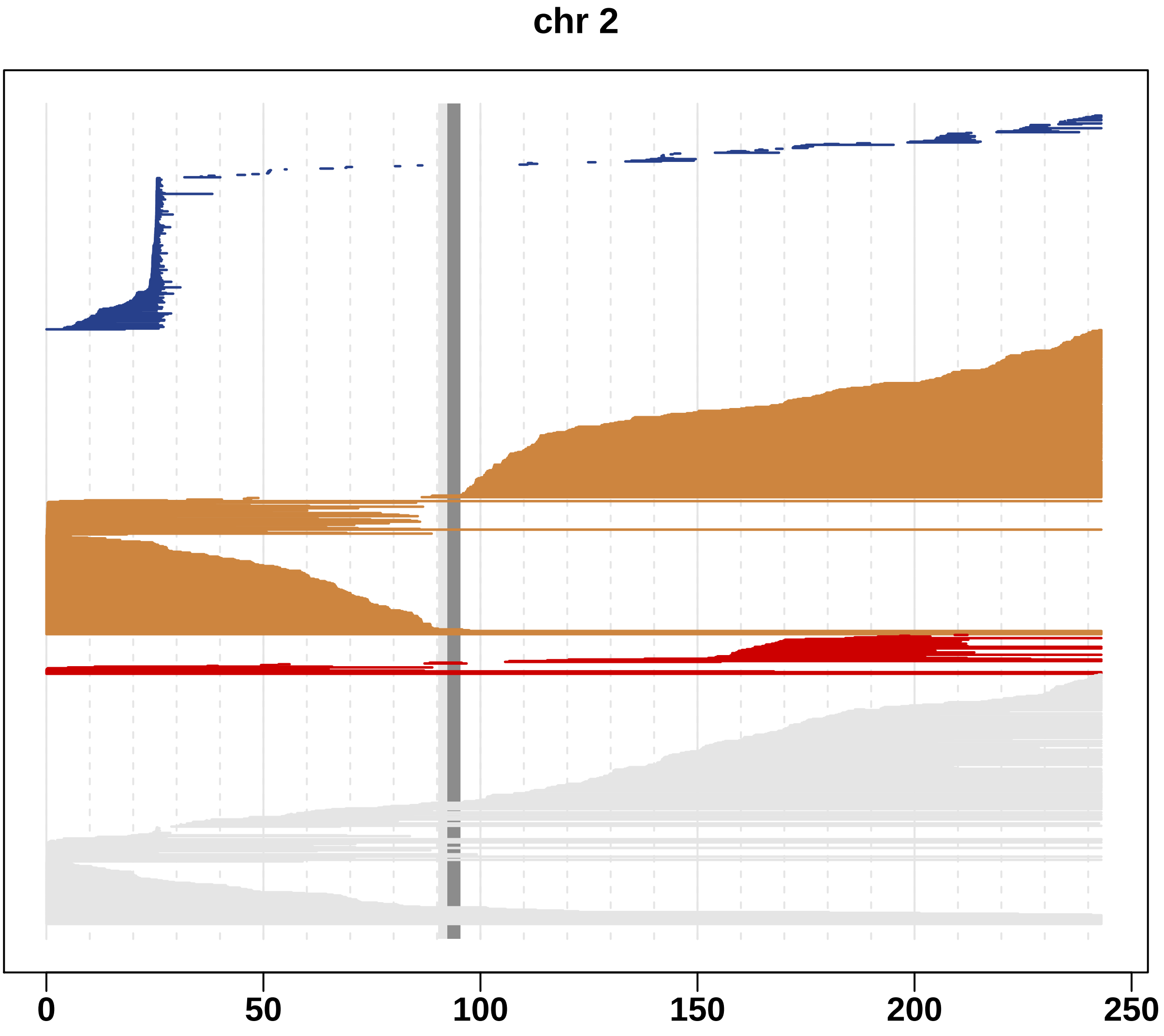

Fig. S3 A landscape of mosaic events in chromosome 3.

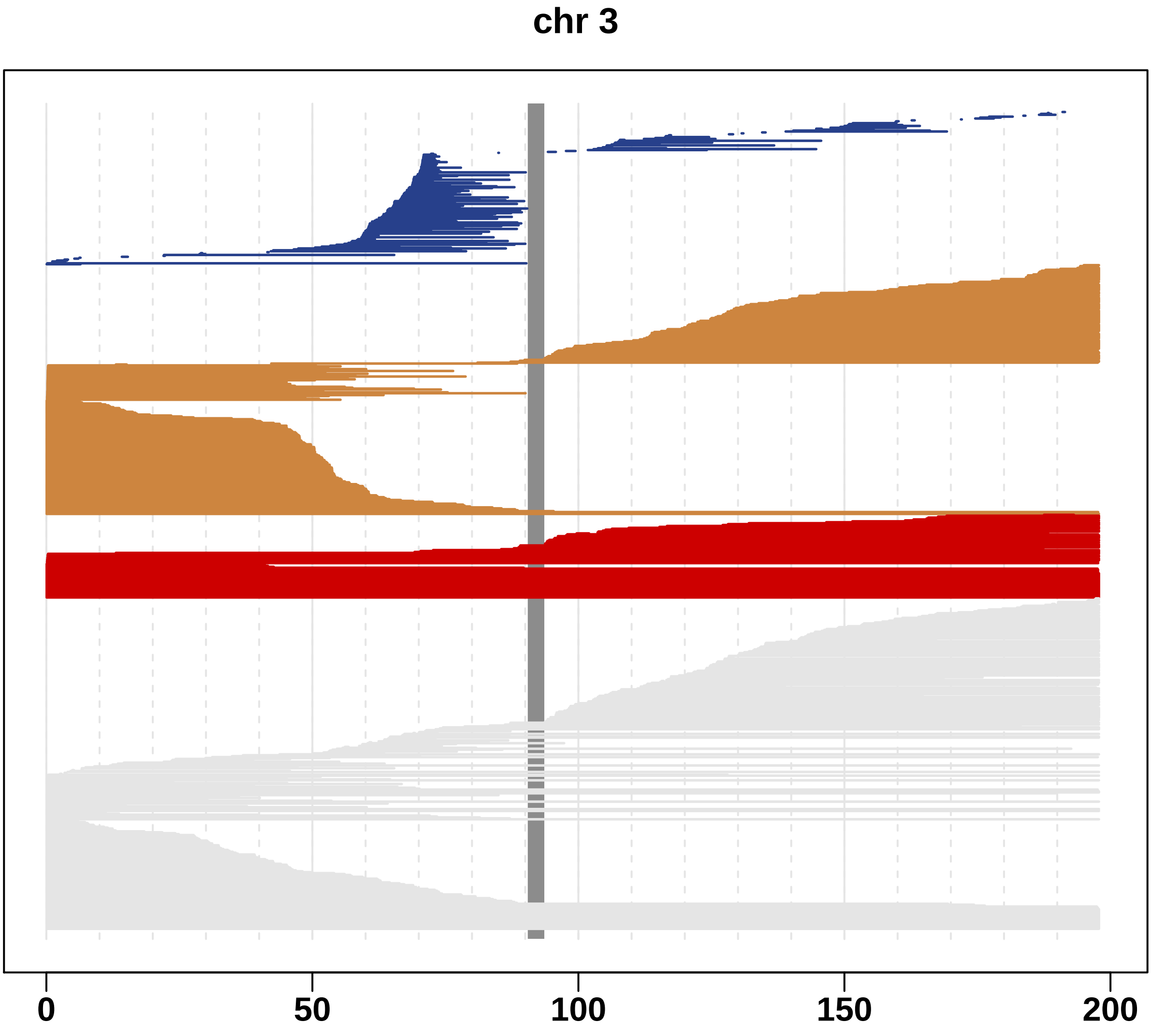

Fig. S4 A landscape of mosaic events in chromosome 4.

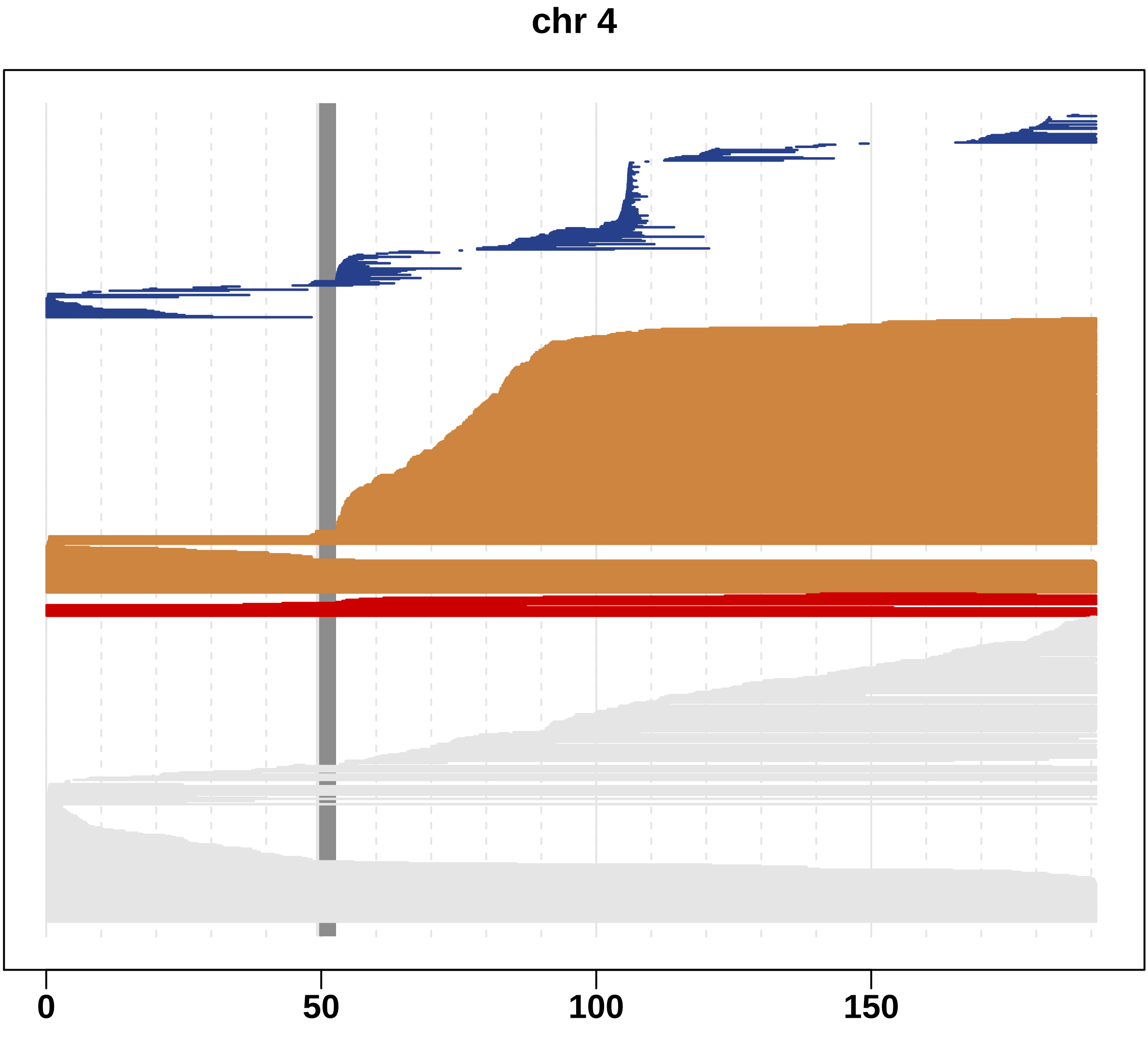

Fig. S5 A landscape of mosaic events in chromosome 5.

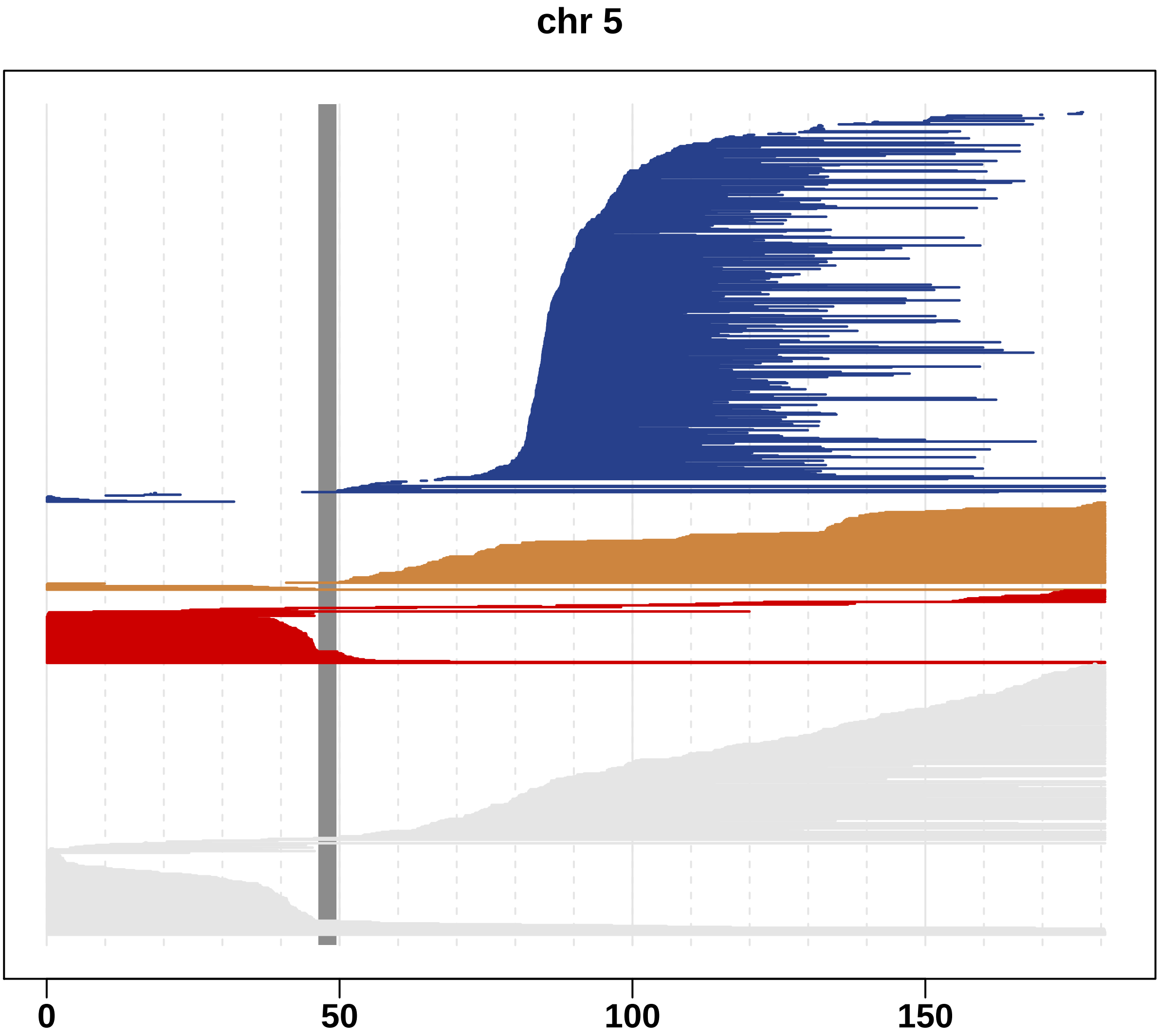

Fig. S6 A landscape of mosaic events in chromosome 6.

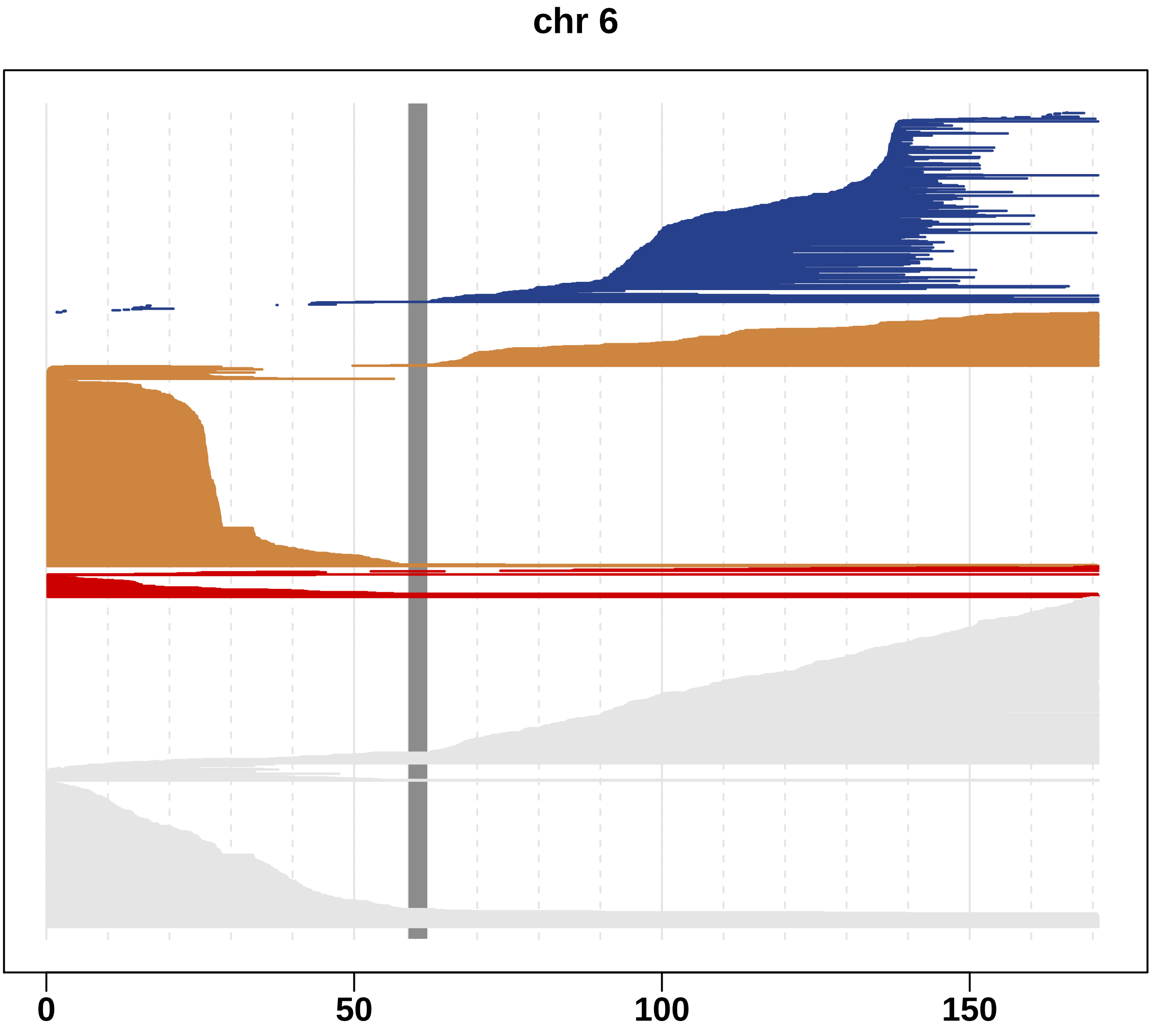

Fig. S7 A landscape of mosaic events in chromosome 7.

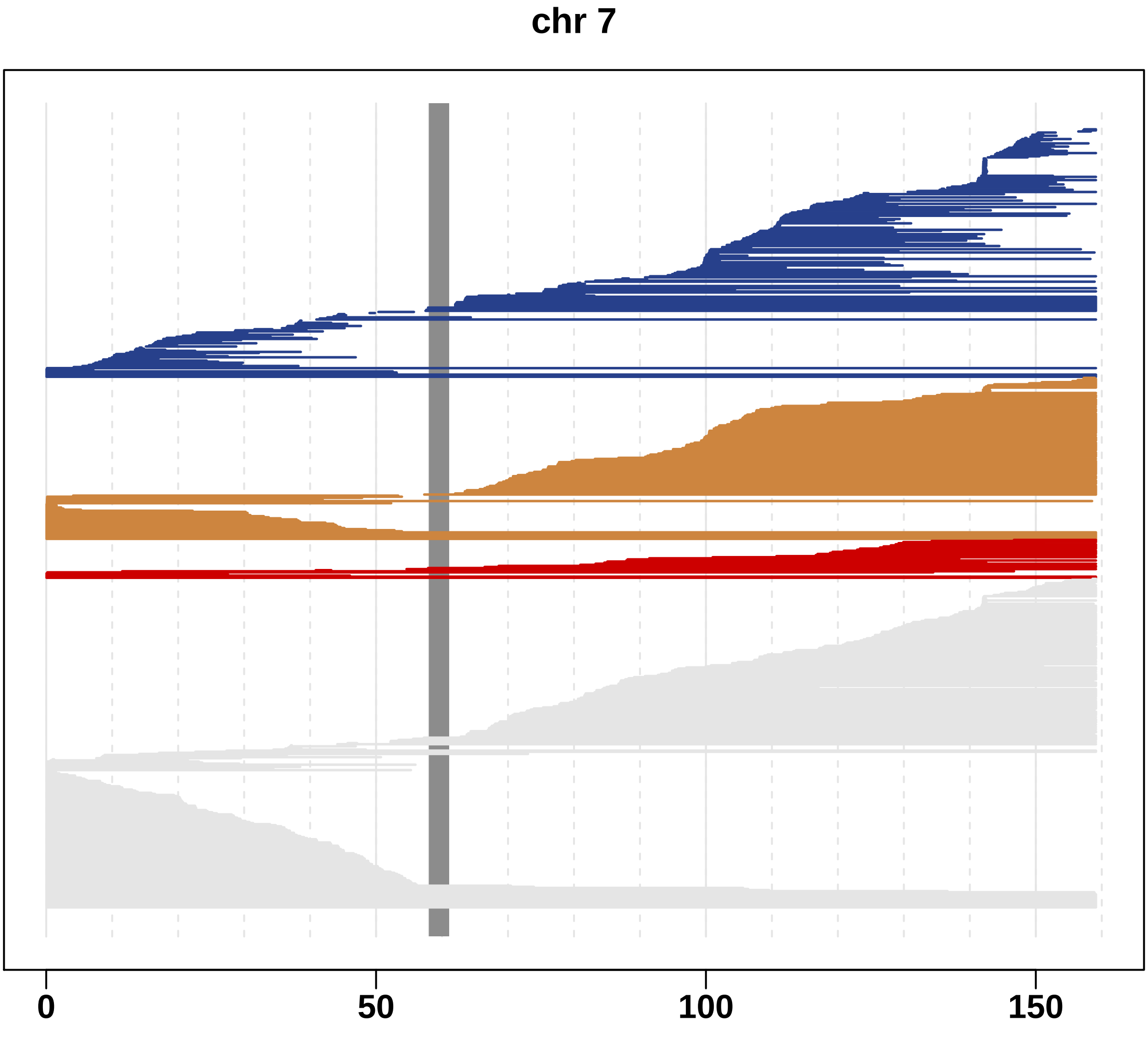

Fig. S8 A landscape of mosaic events in chromosome 8.

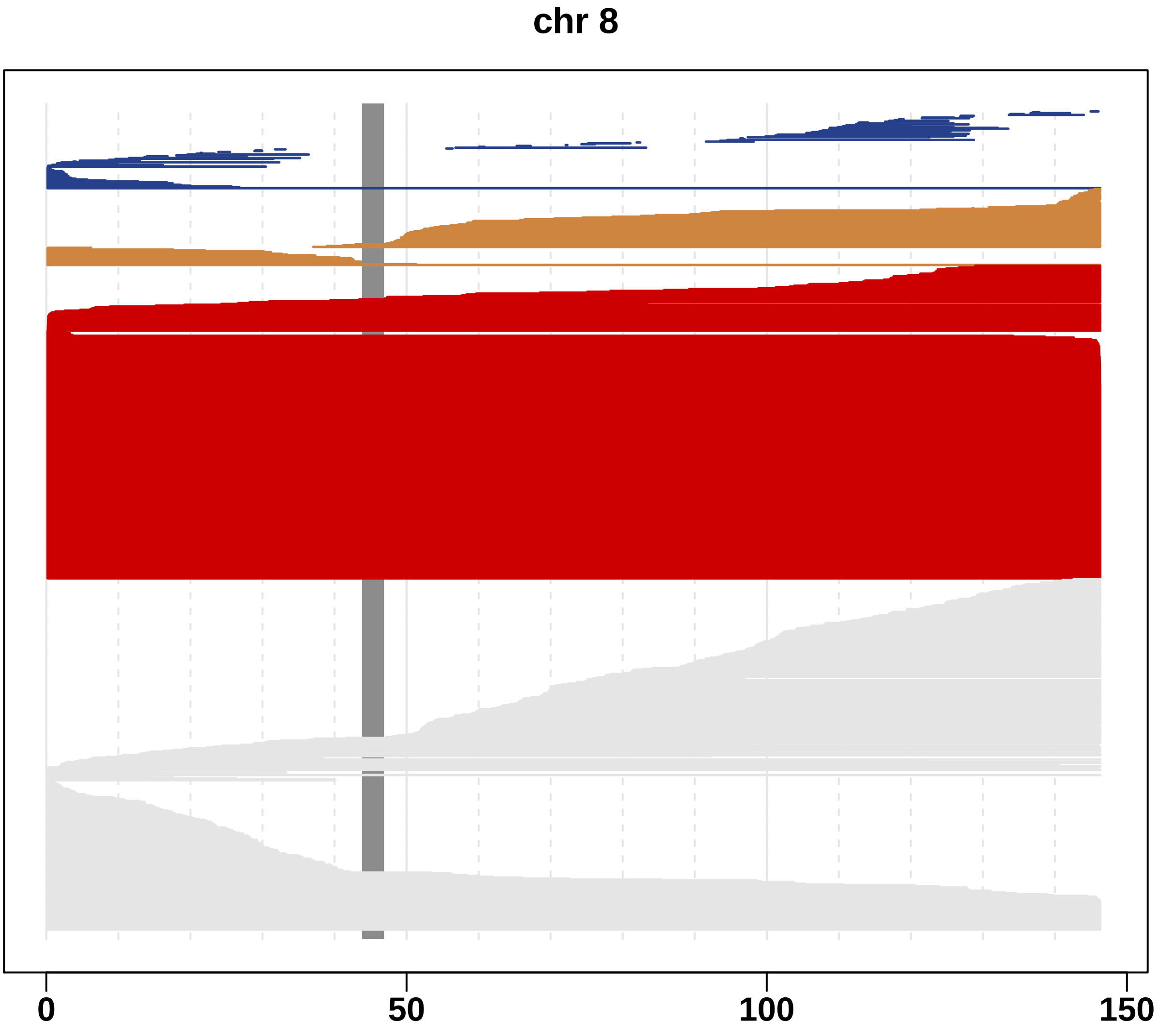

Fig. S9 A landscape of mosaic events in chromosome 9.

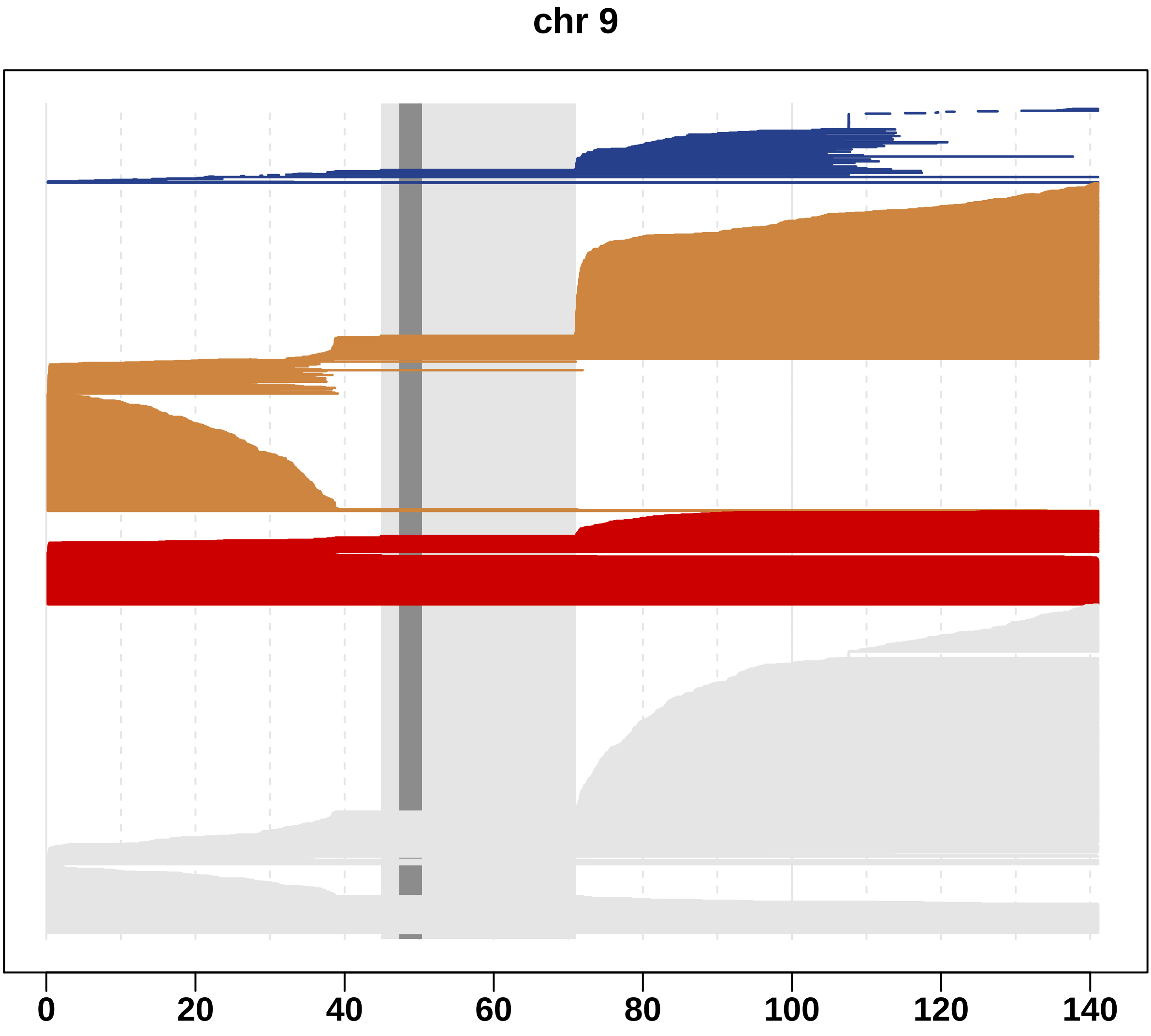

Fig. S10 A landscape of mosaic events in chromosome 10.

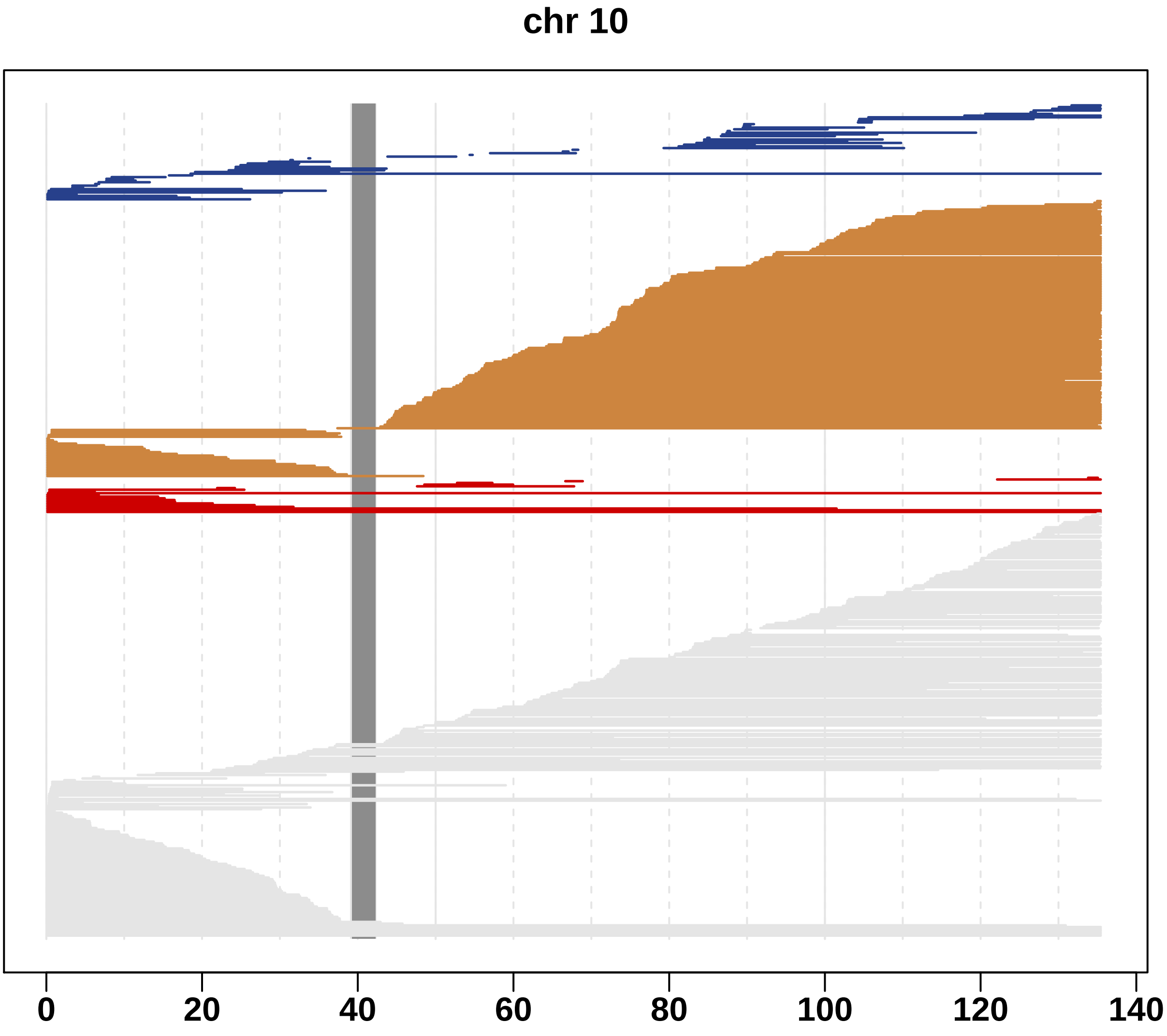

Fig. S11 A landscape of mosaic events in chromosome 11.

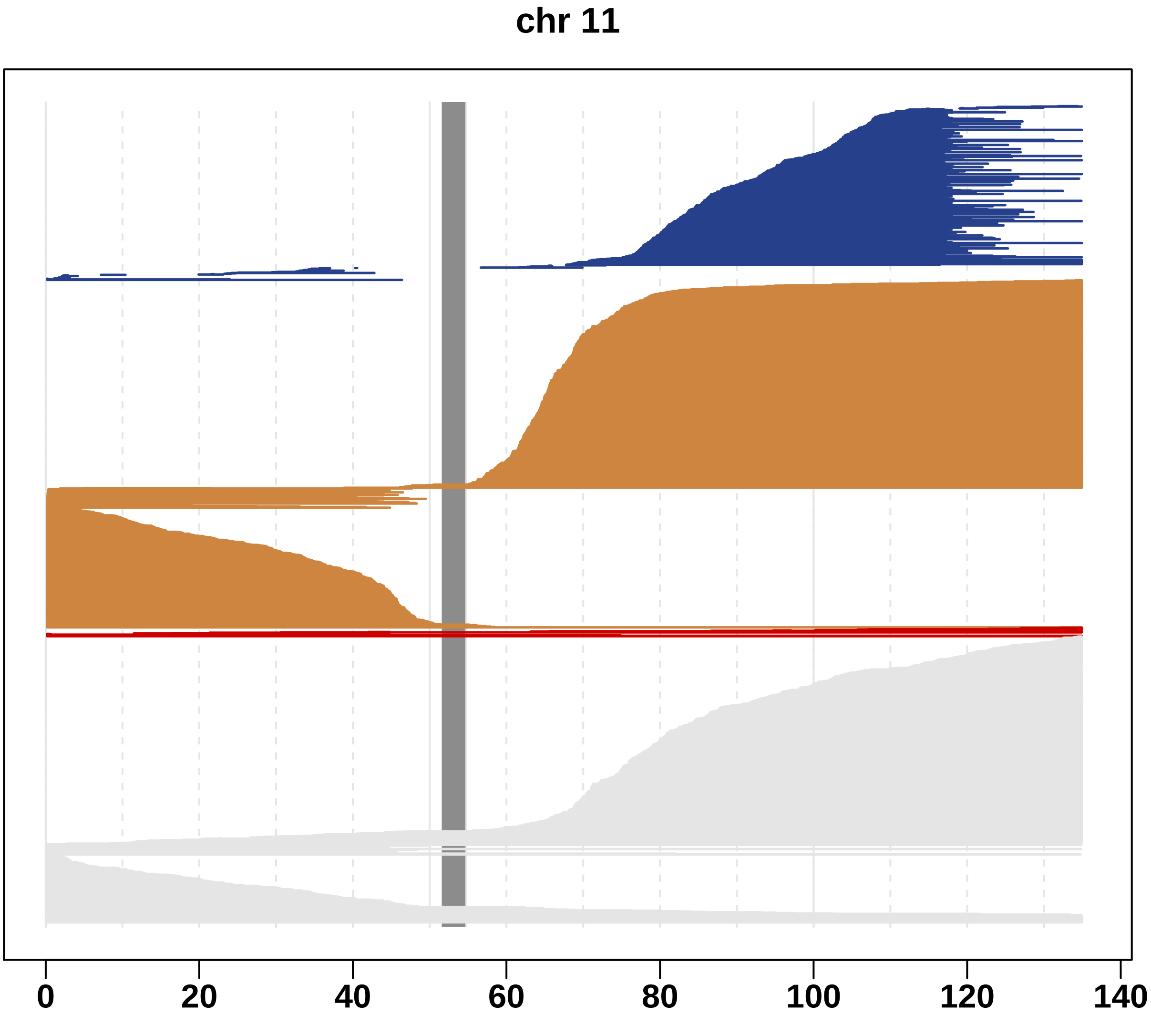

Fig. S12 A landscape of mosaic events in chromosome 12.

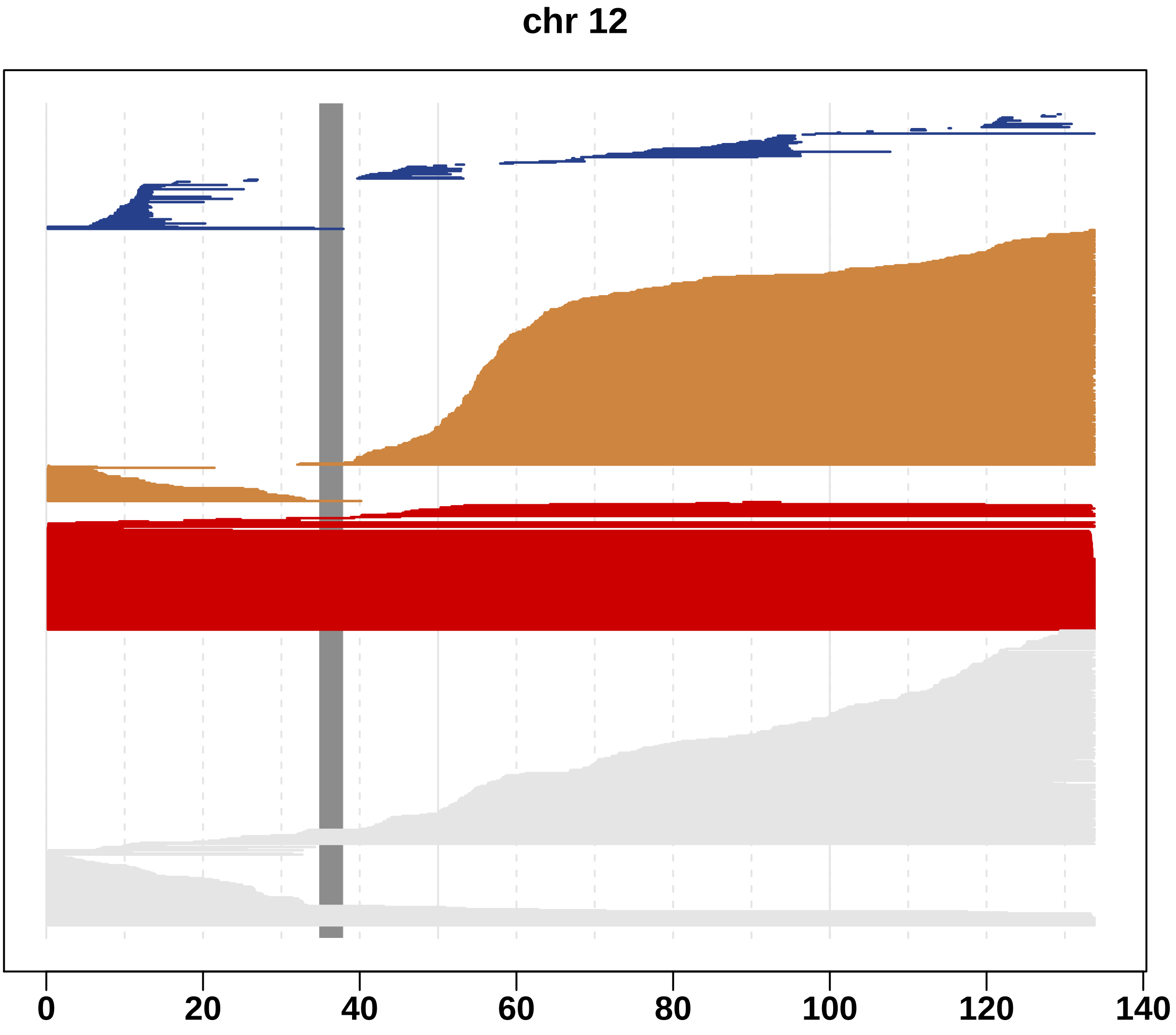

Fig. S13 A landscape of mosaic events in chromosome 13.

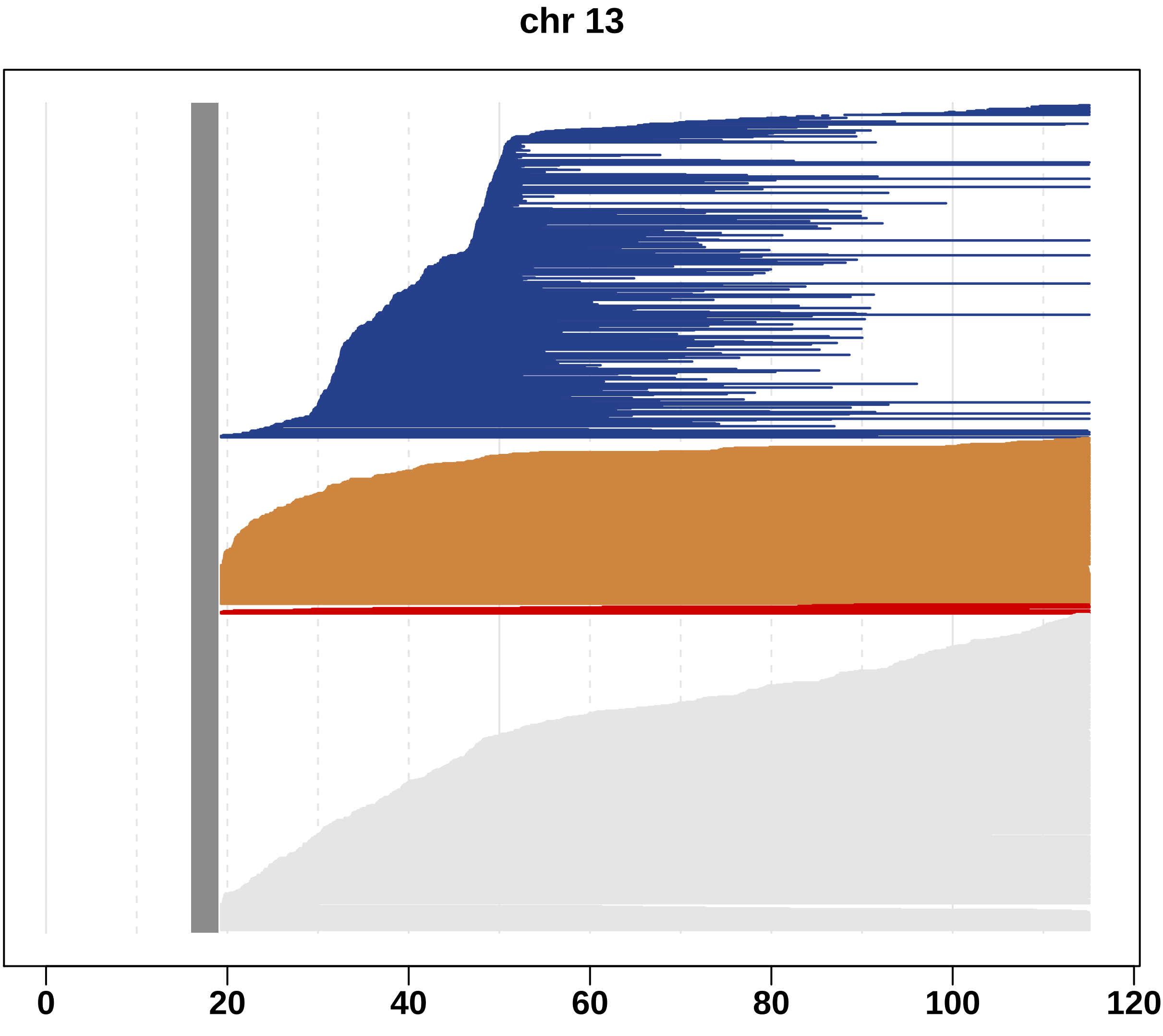

Fig. S14 A landscape of mosaic events in chromosome 14.

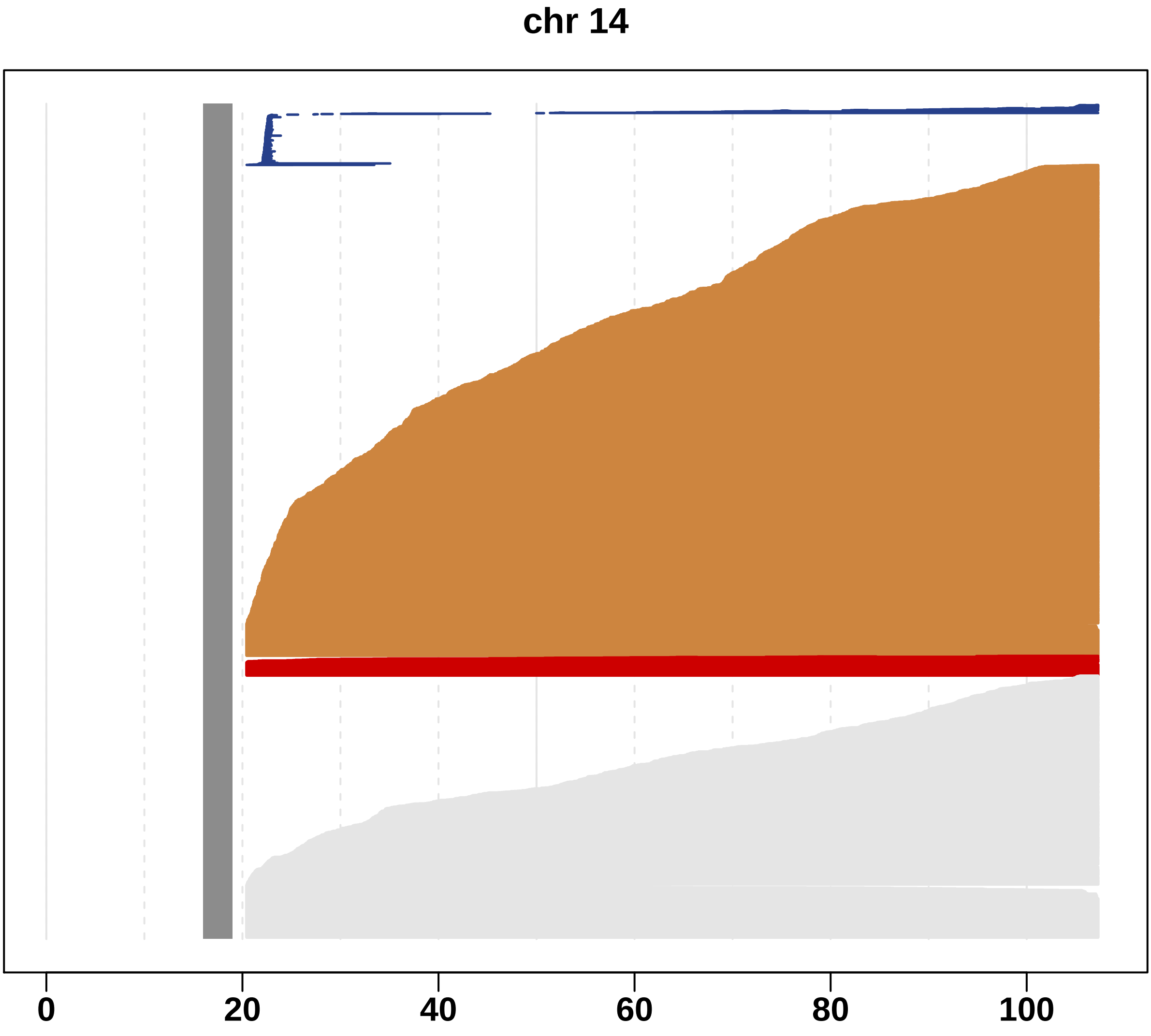

Fig. S15 A landscape of mosaic events in chromosome 15.

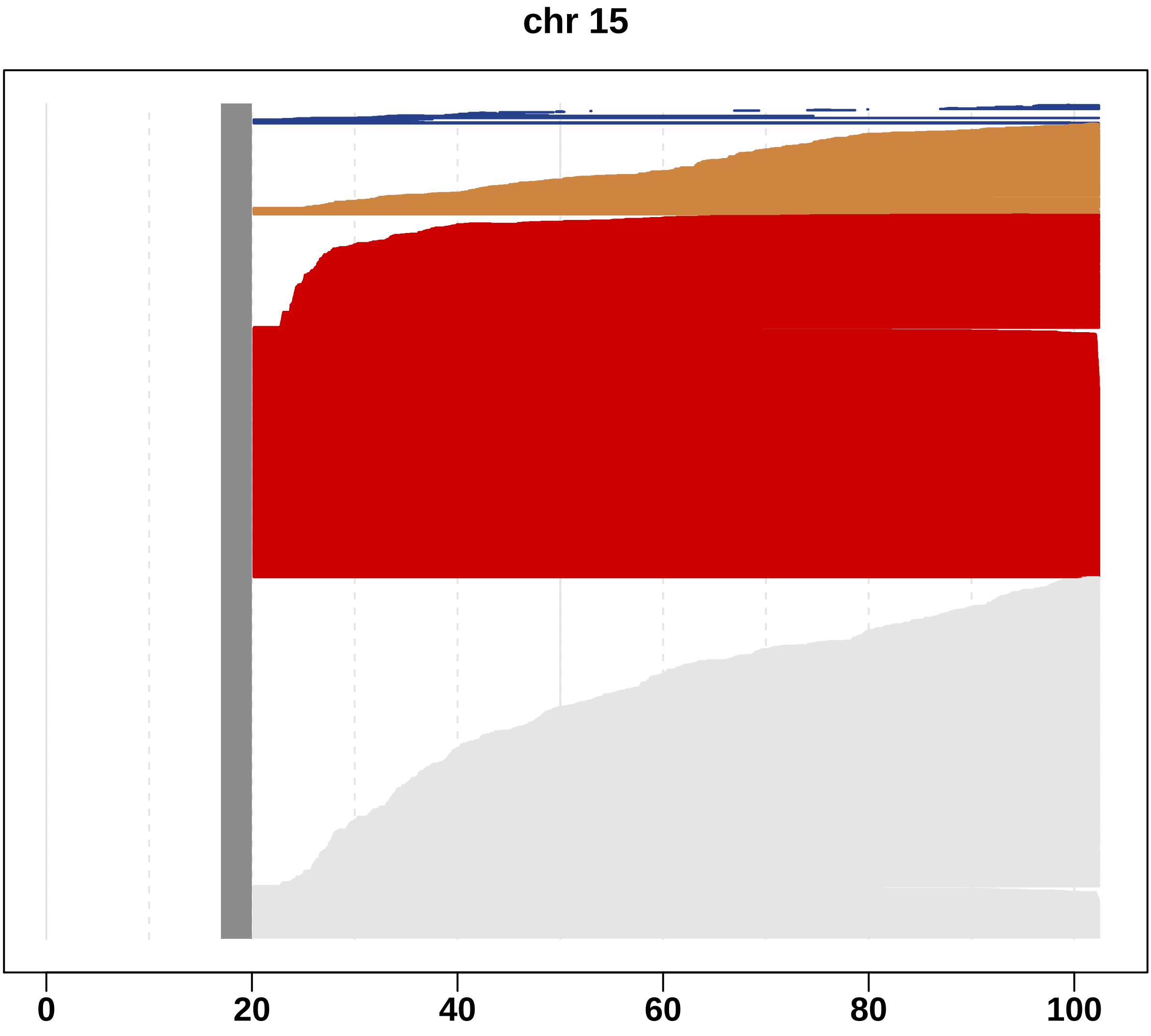

Fig. S16 A landscape of mosaic events in chromosome 16.

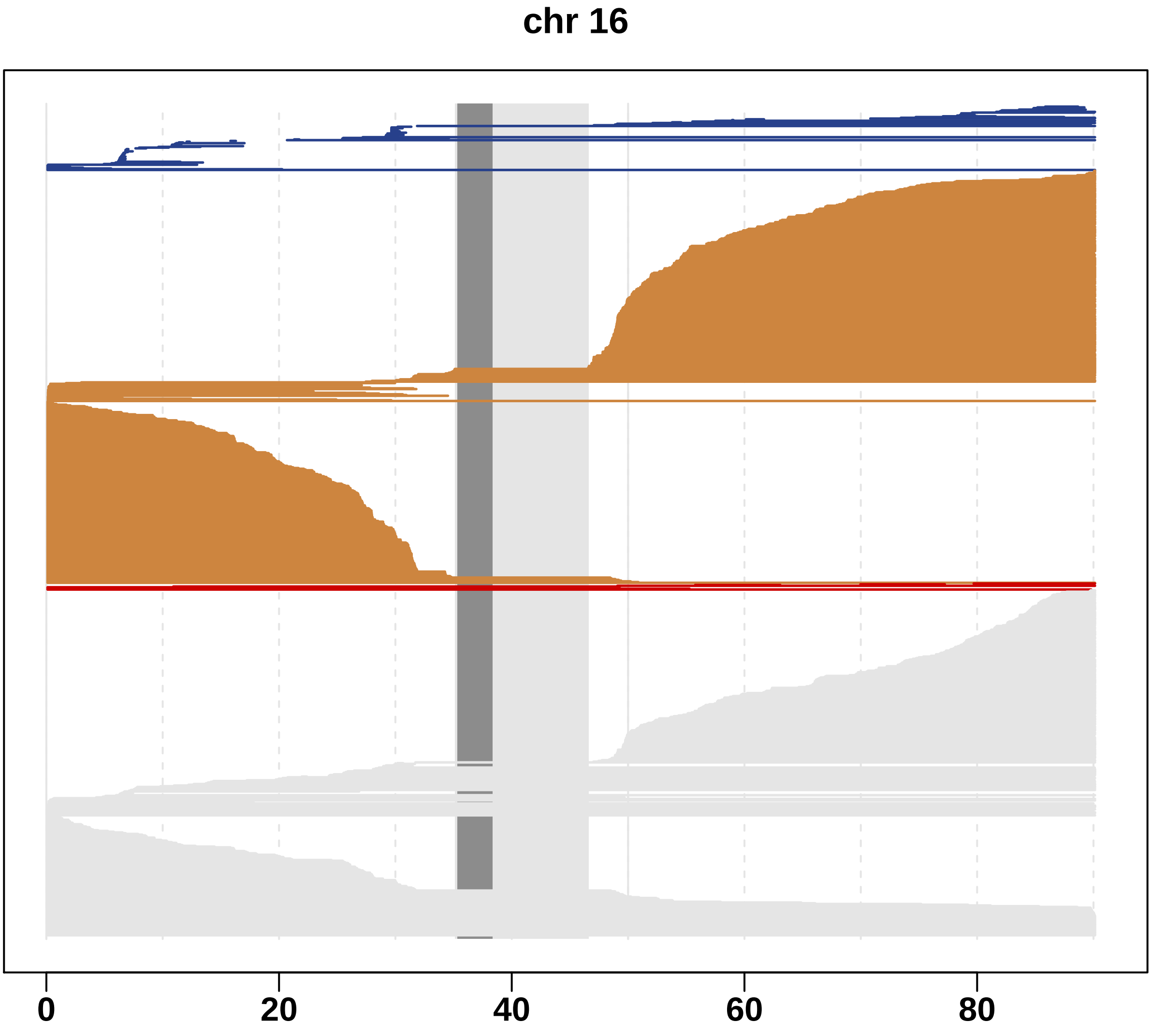

Fig. S17 A landscape of mosaic events in chromosome 17.

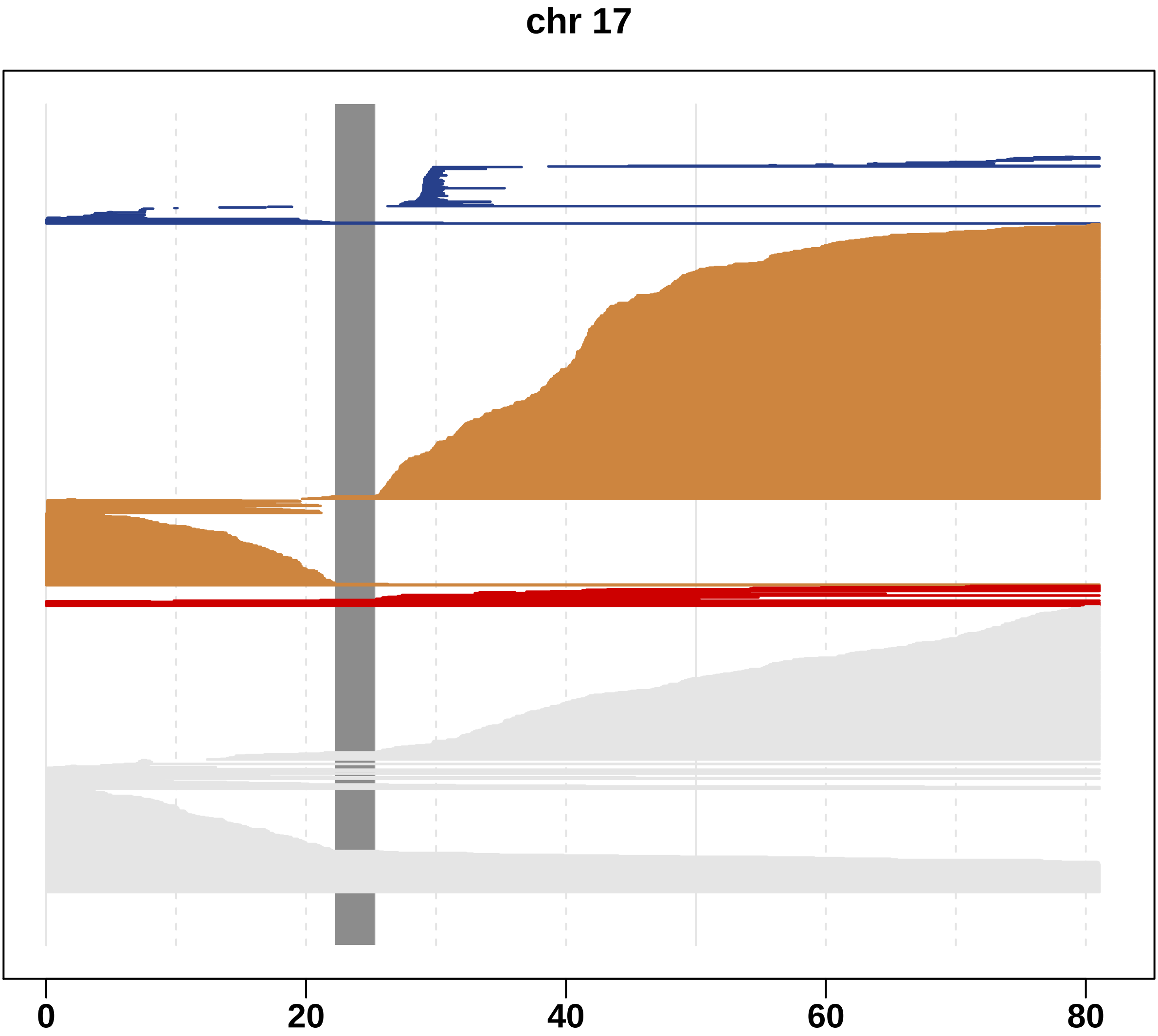

Fig. S18 A landscape of mosaic events in chromosome 18.

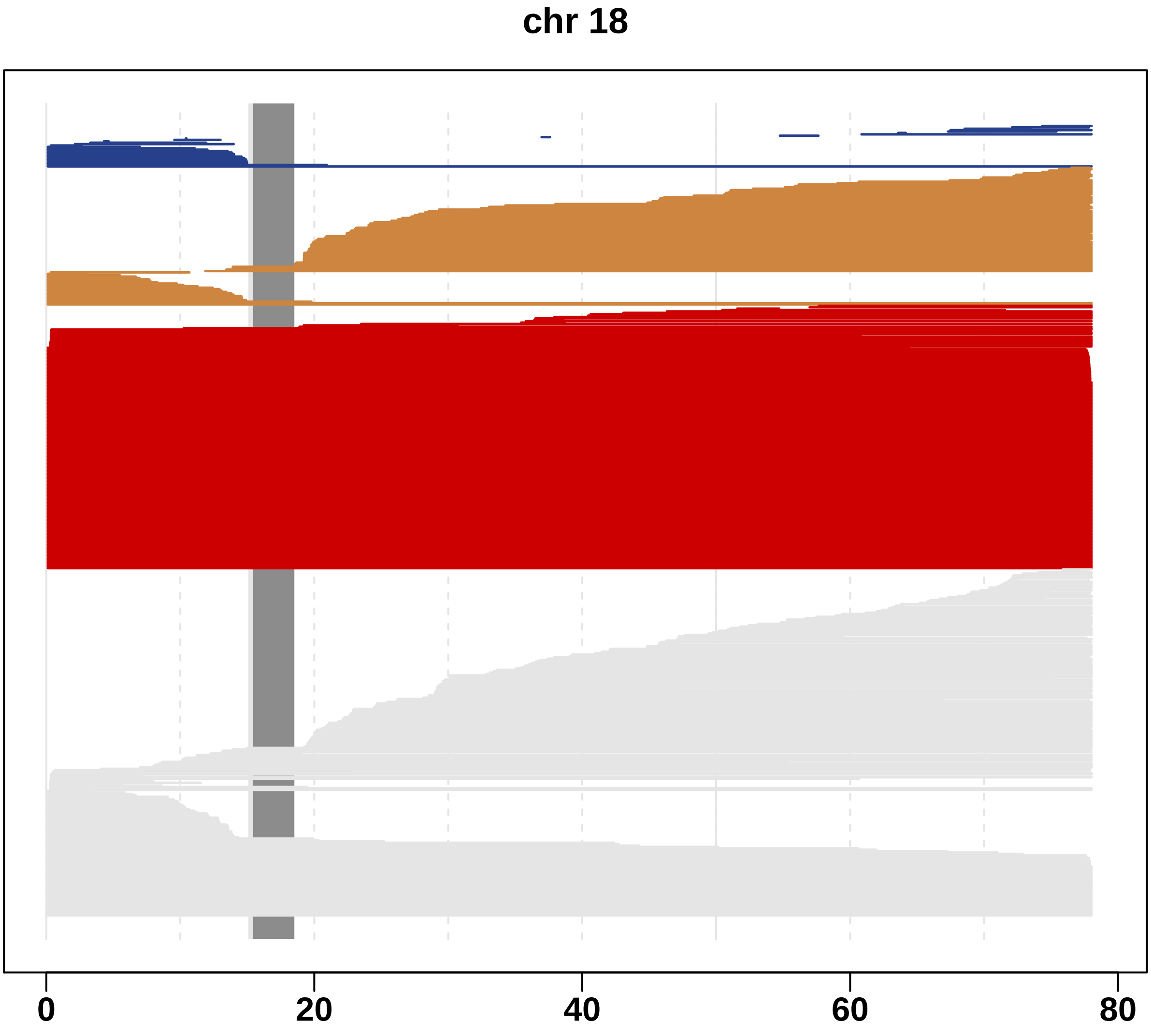

Fig. S19 A landscape of mosaic events in chromosome 19.

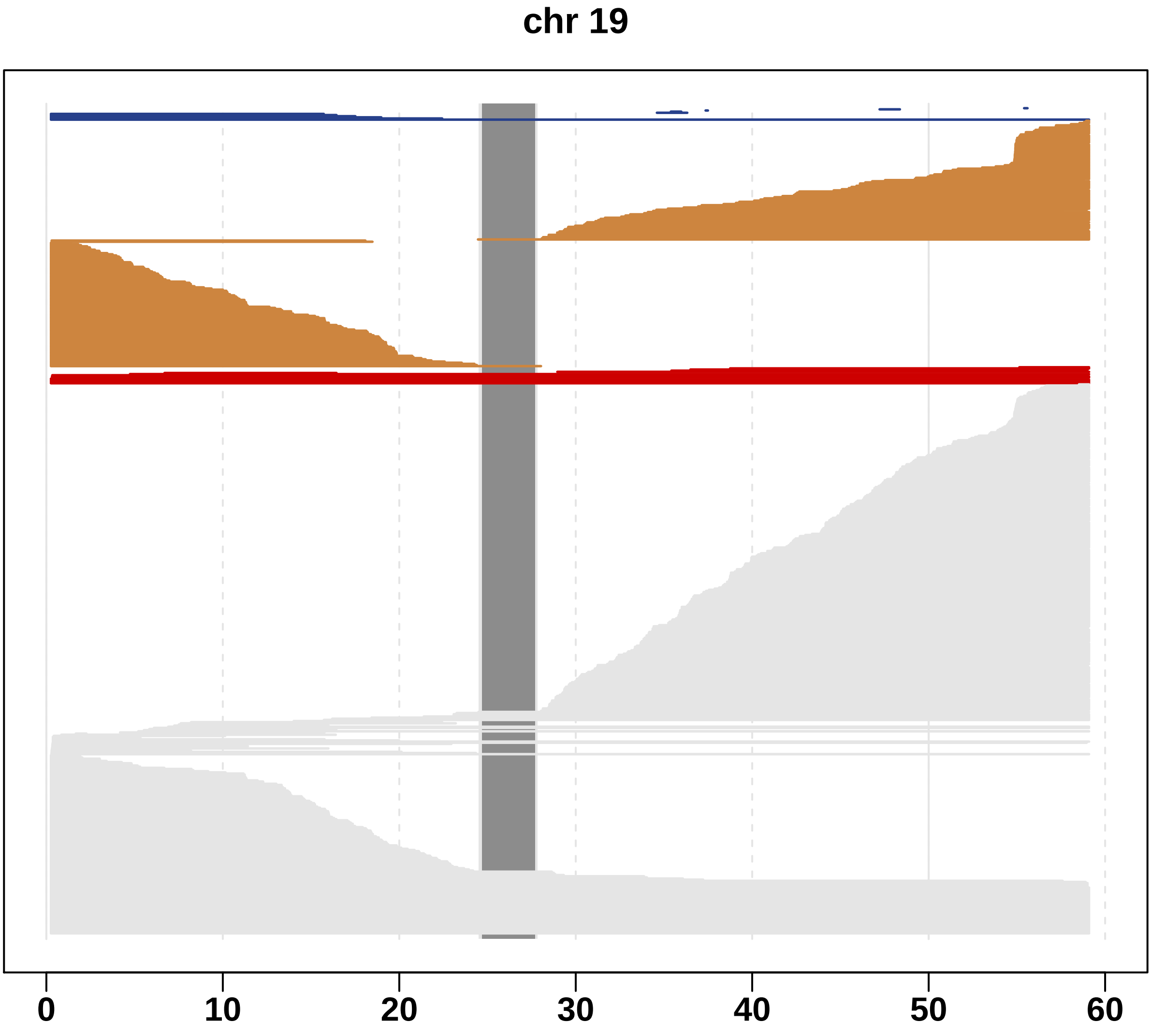

Fig. S20 A landscape of mosaic events in chromosome 20.

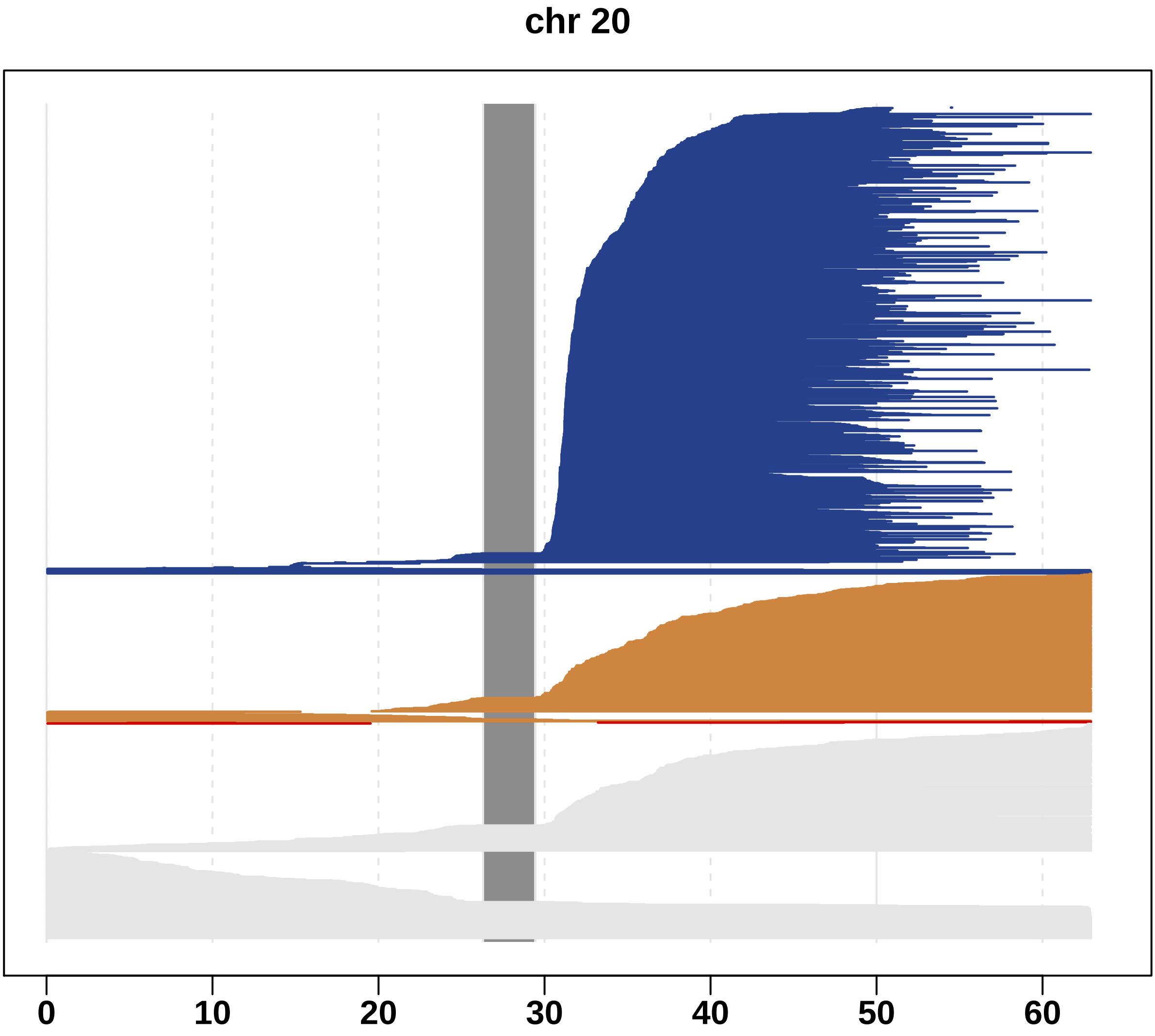

Fig. S21 A landscape of mosaic events in chromosome 21.

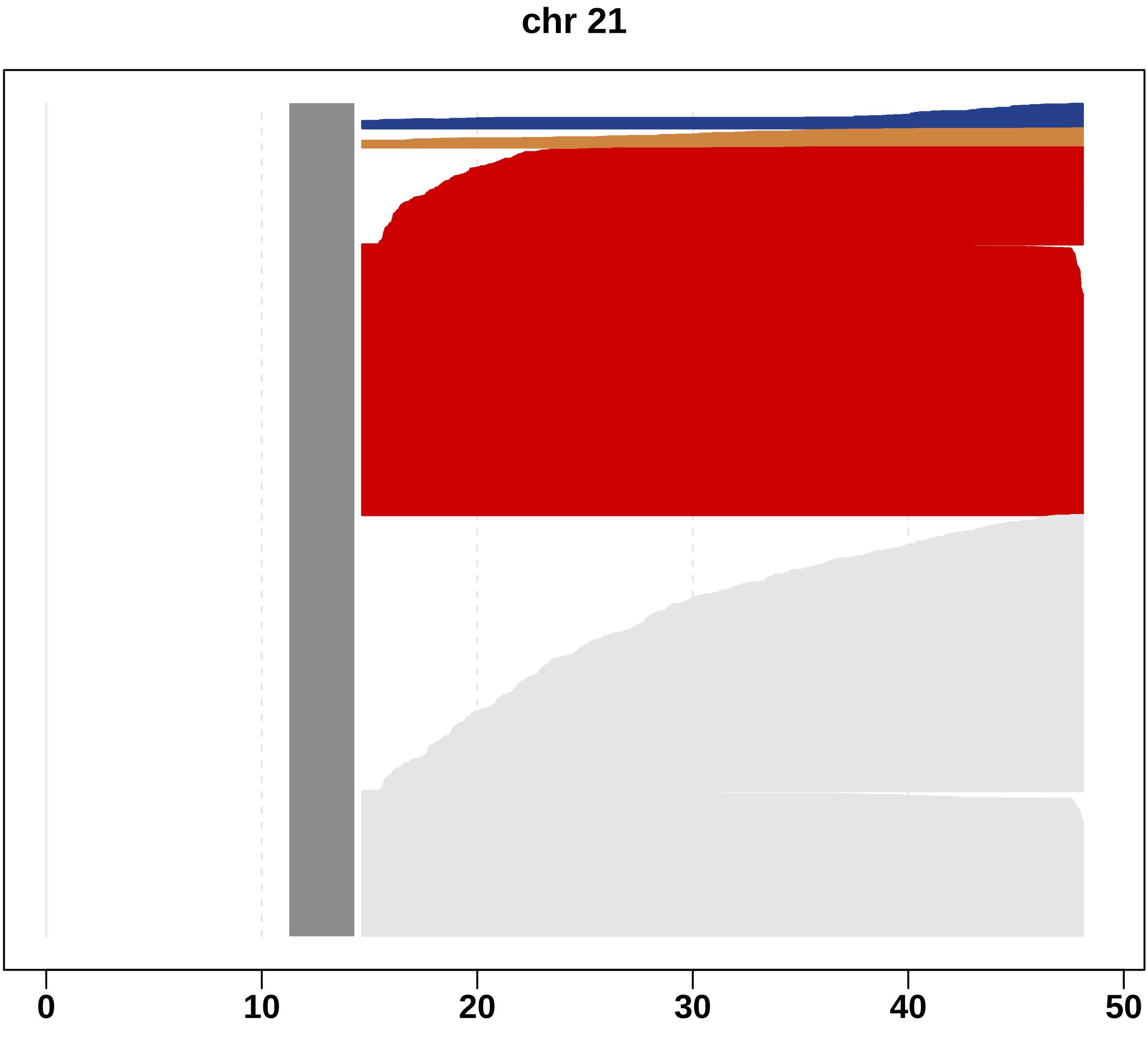

Fig. S22 A landscape of mosaic events in chromosome 22.

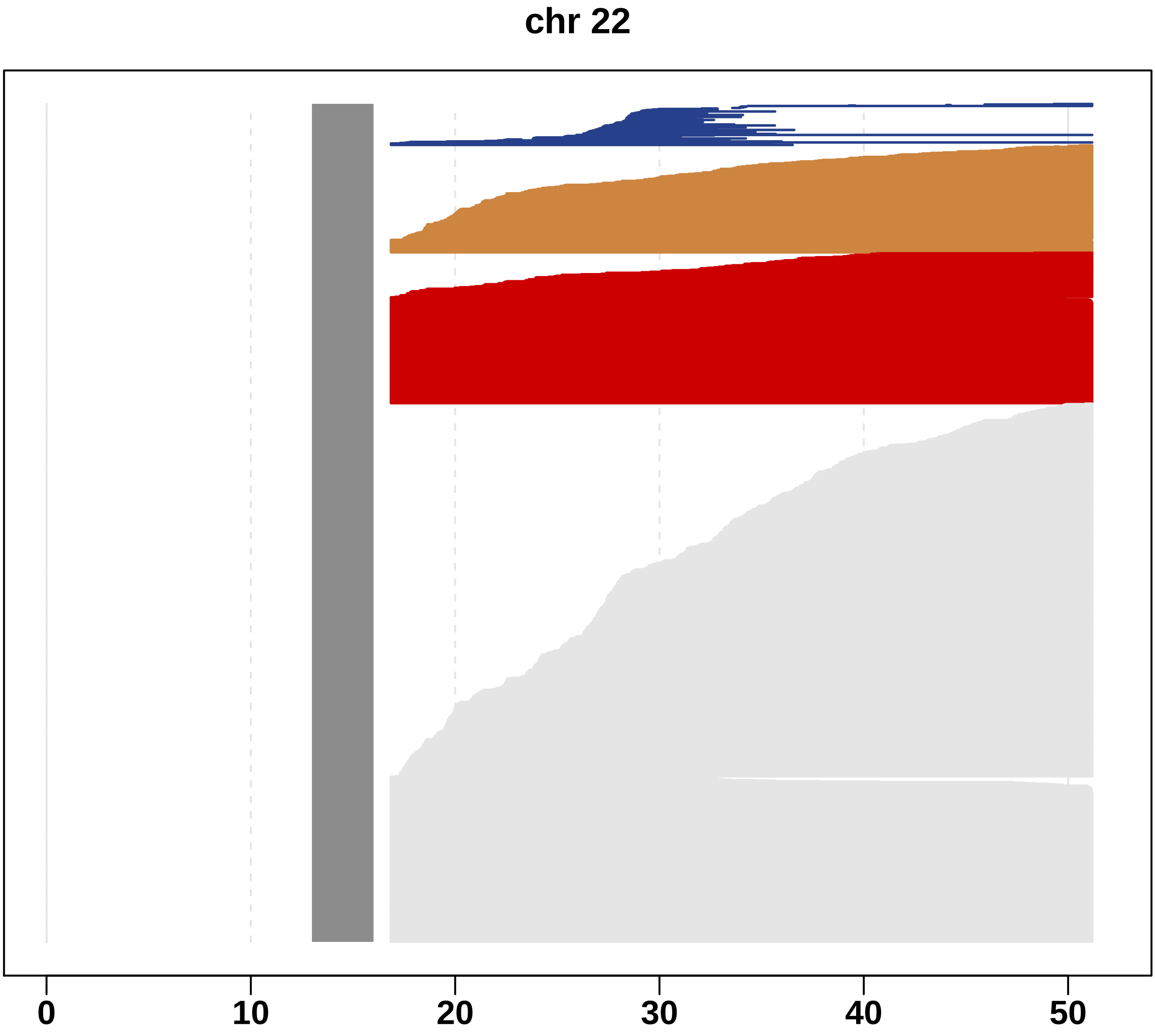

Fig. S23. Cell fractions of multiple mosaic events occurring in the same subject.

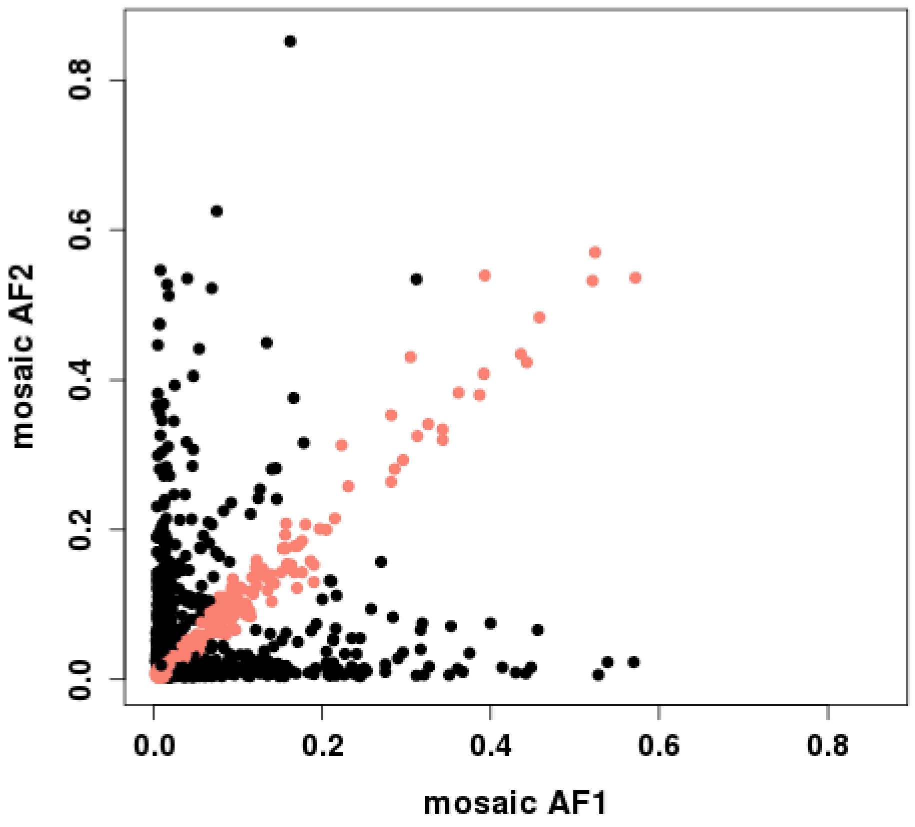

Mosaic allelic fractions (AF) are plotted in subjects with 2 mosaic events in different chromosomes. We computed difference in cell fractions between two mosaic events and regarded cell fractions as different if the difference in fractions satisfied the two conditions; 1) difference is more than 50% of the smaller AF and 2) difference is more than 0.01. As a result, 54.5% of subjects were estimated to have different cell fractions for mosaic events, suggesting multiple clones. Since it is hard to distinguish different clones with similar fractions in the remaining subjects, 54.5% should be considered as a minimum number.

Fig S24. Age and sex of carriers of mosaic event types.

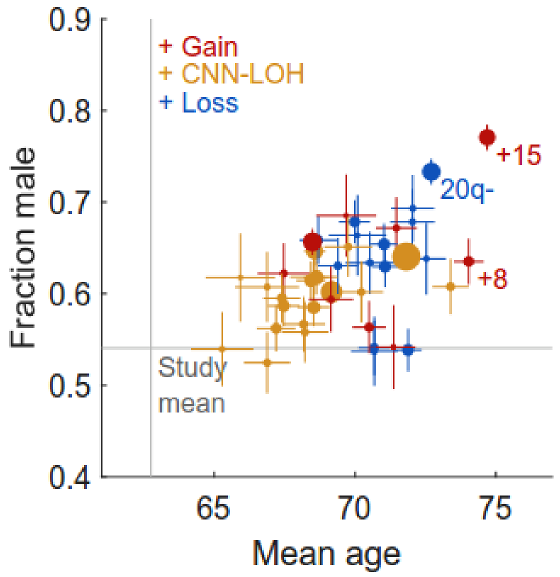

Mean age and sex of carriers of specific mCA types (defined by chromosome and copy number) with at least 100 carriers. Marker sizes are proportional to mCA frequencies. Error bars, s.e.m. Numeric data are provided in Table S7.

#### Fig. S25 Coverage of mosaic loss in chromosome 1.

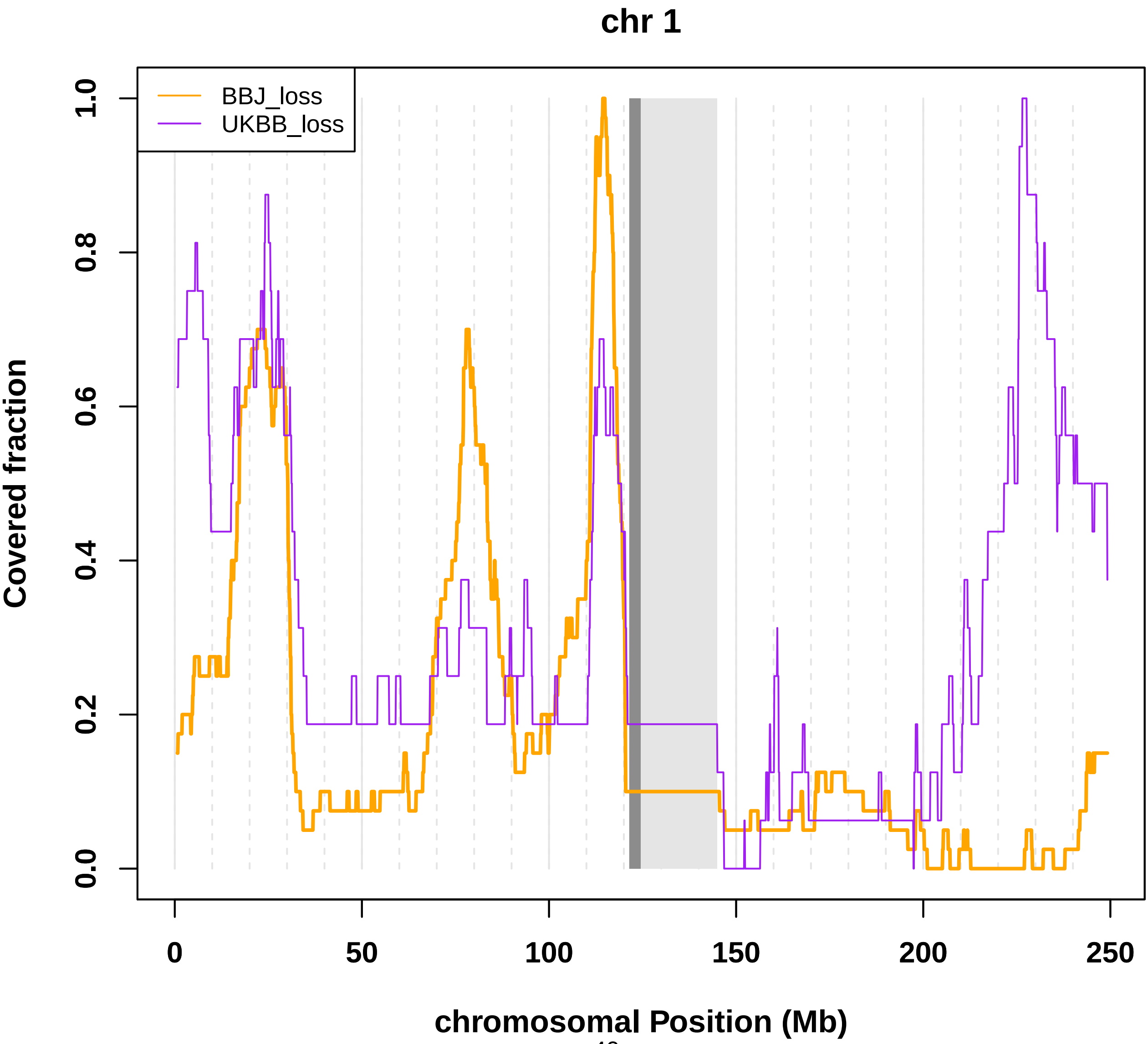

**Fig. S26 Coverage of mosaic loss in chromosome 2.**

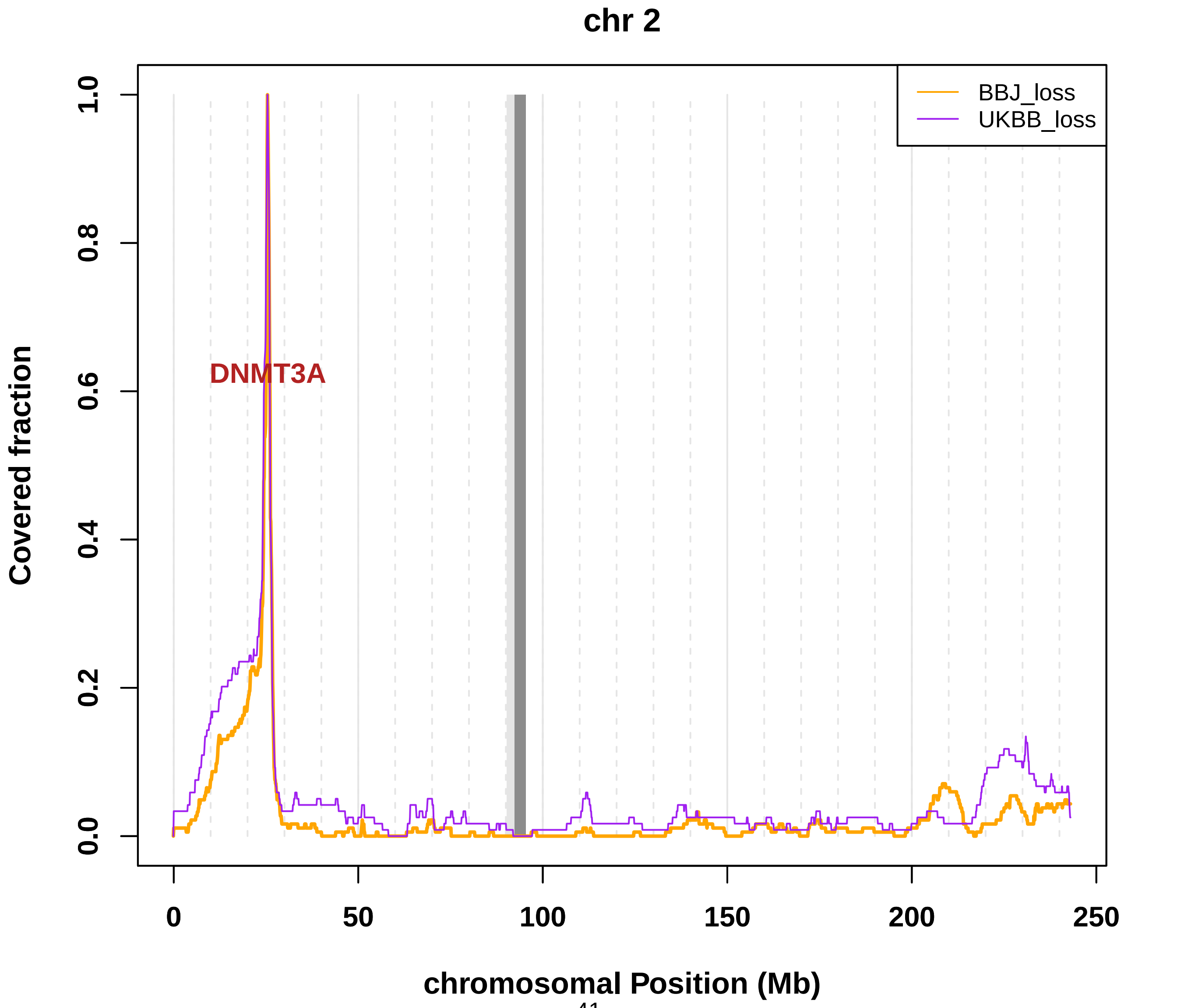

**Fig. S27 Coverage of mosaic loss in chromosome 3.**

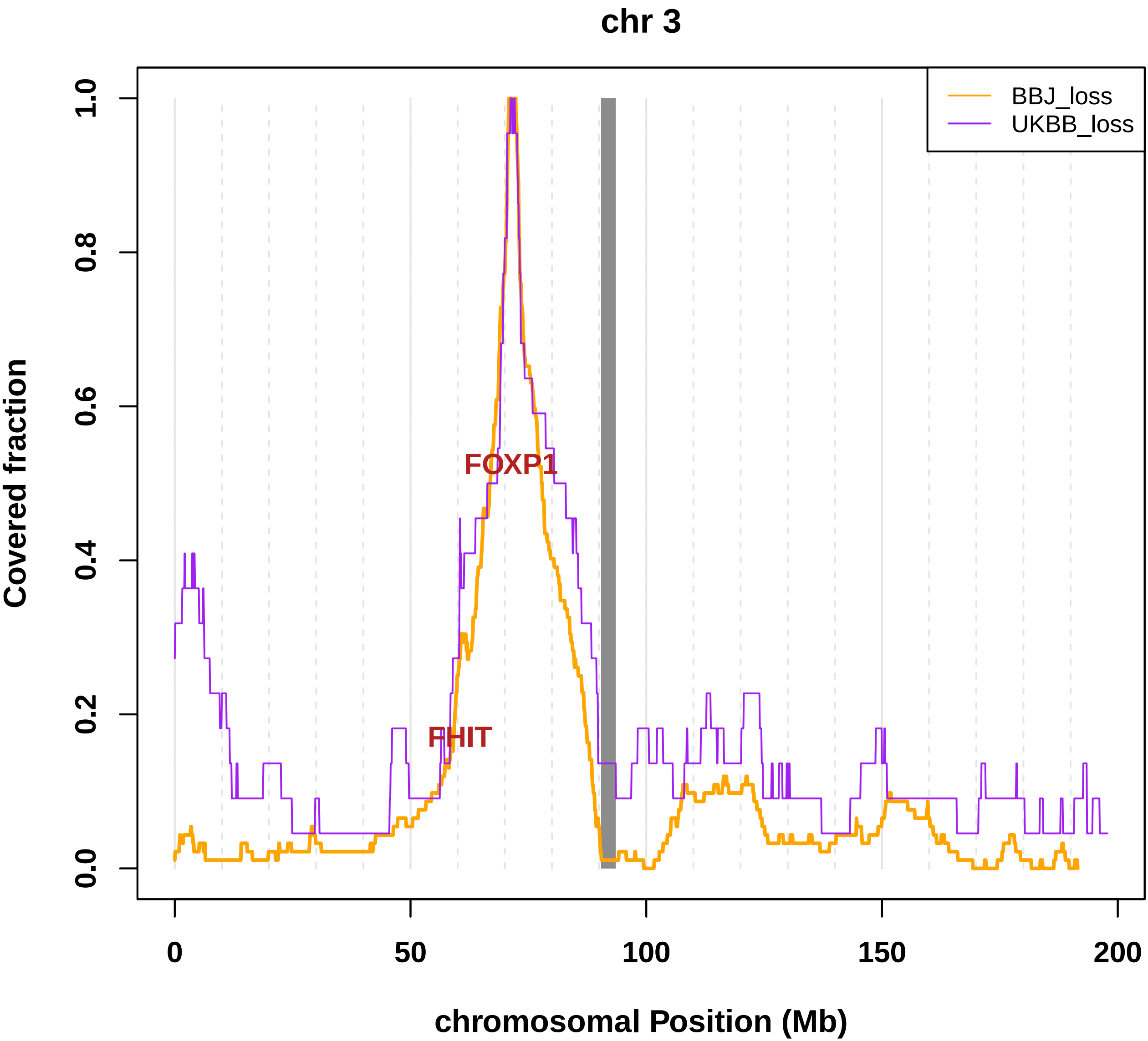

**Fig. S28 Coverage of mosaic loss in chromosome 4.**

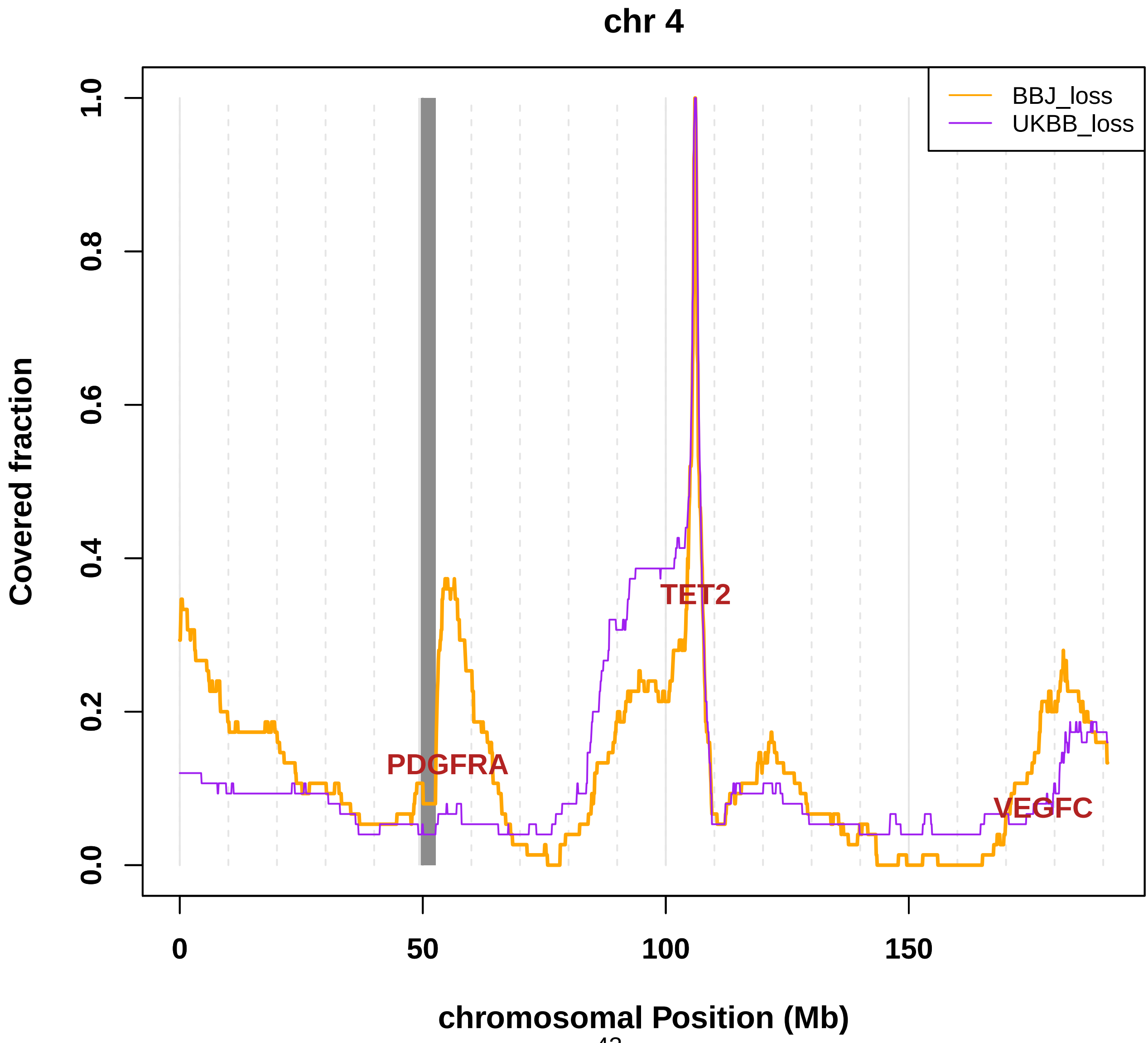

### Fig. S29 Coverage of mosaic loss in chromosome 5.

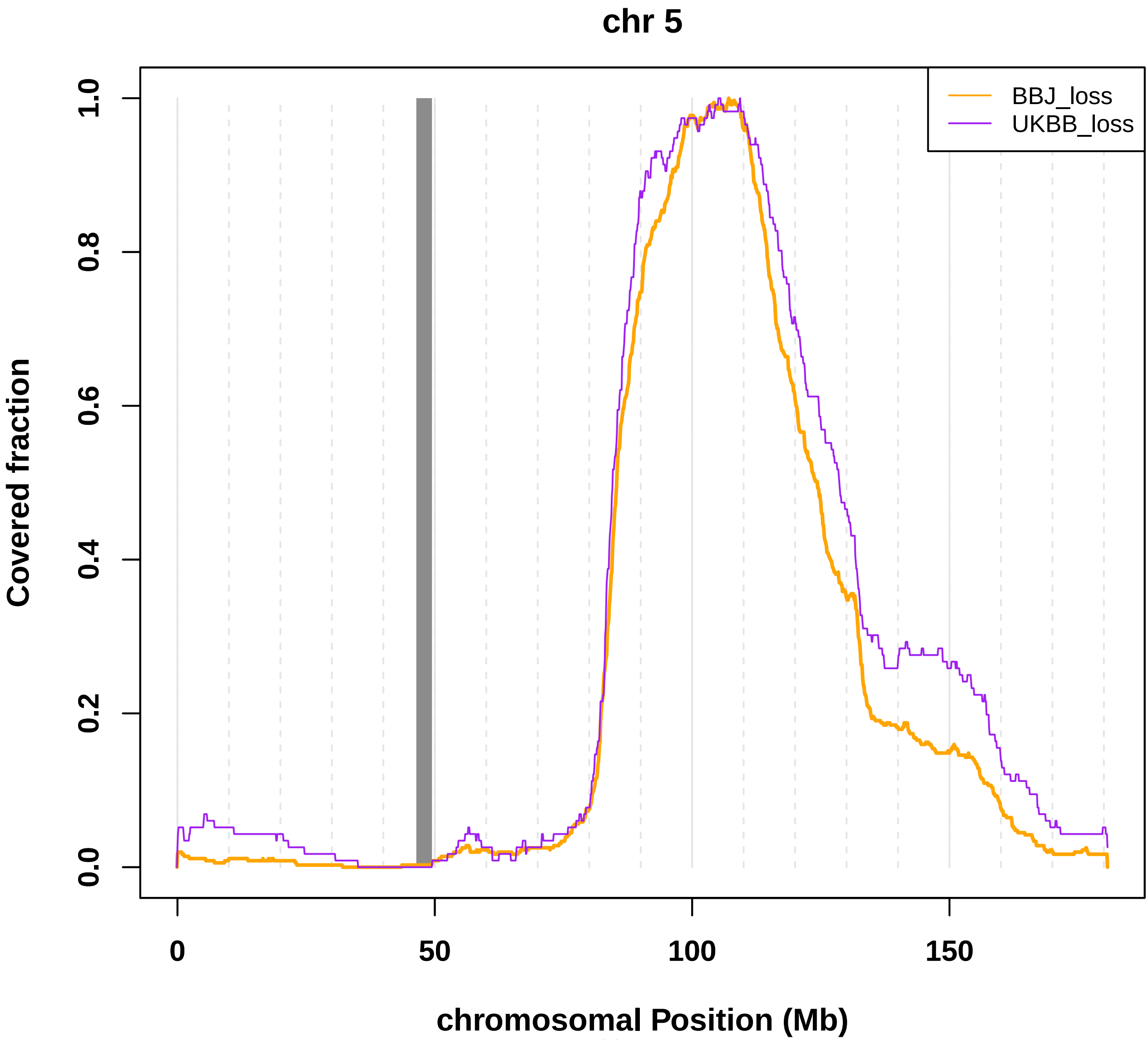

**Fig. S30 Coverage of mosaic loss in chromosome 6.**

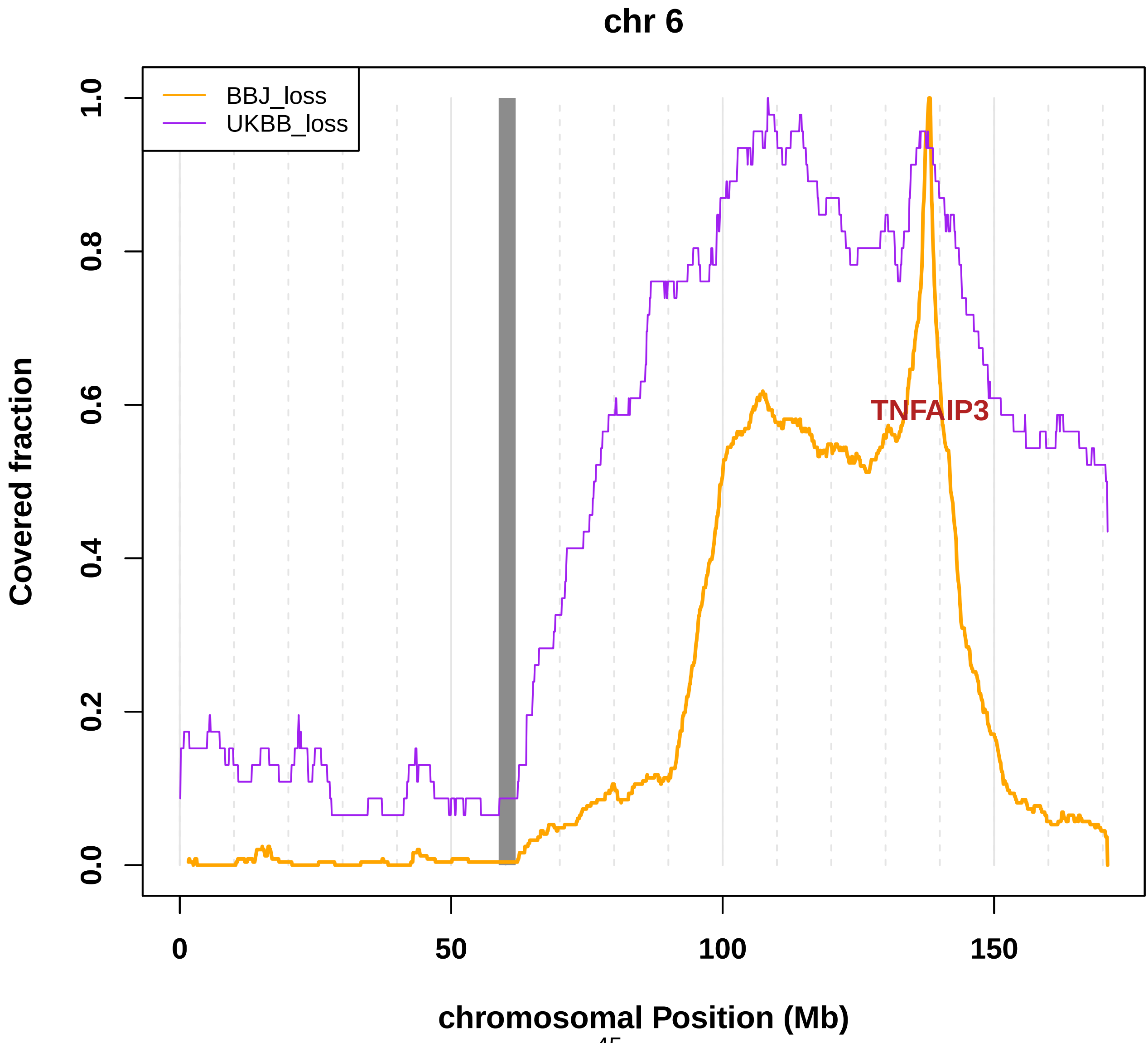

**Fig. S31 Coverage of mosaic loss in chromosome 7.**

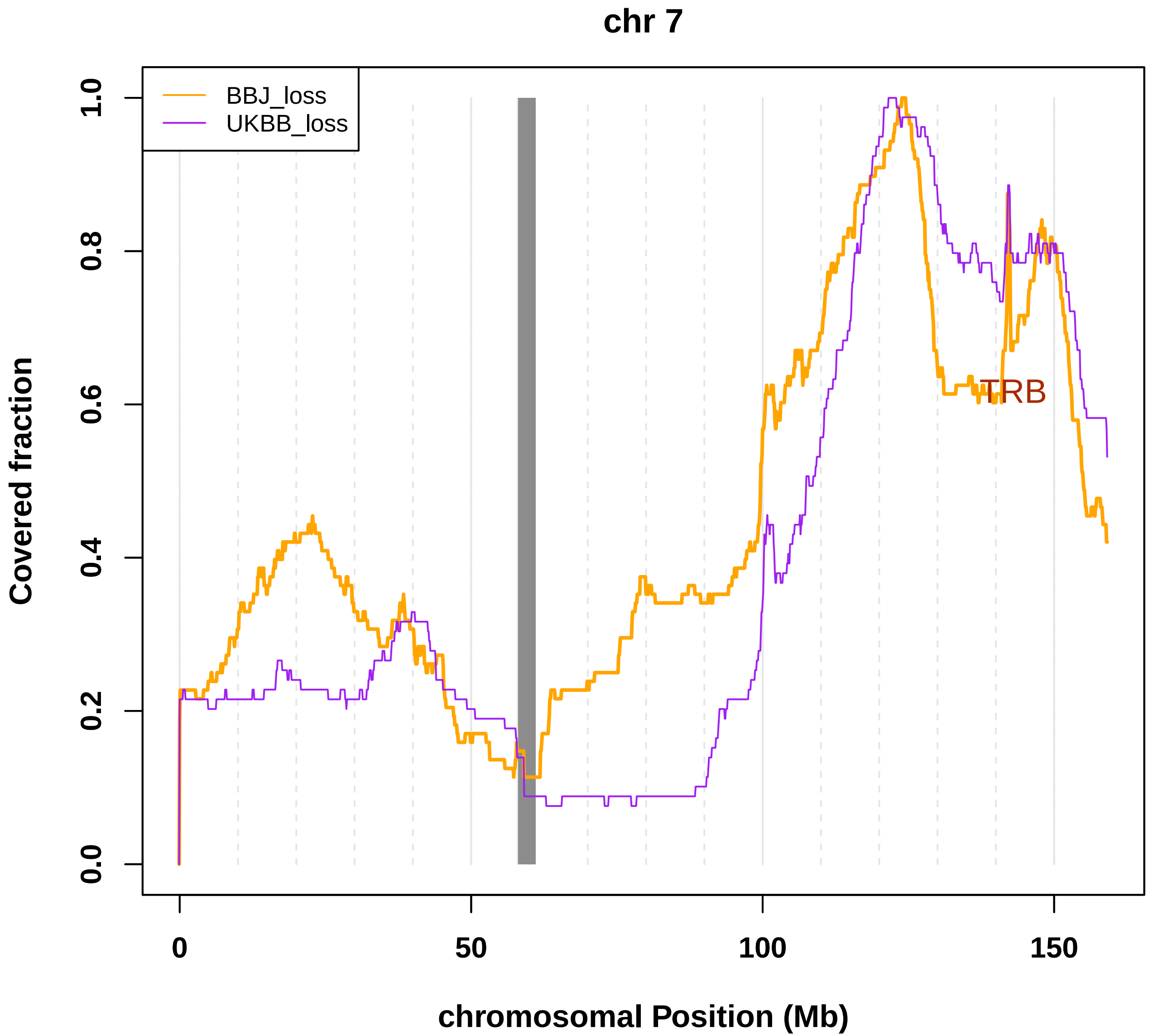

**Fig. S32 Coverage of mosaic loss in chromosome 8.**

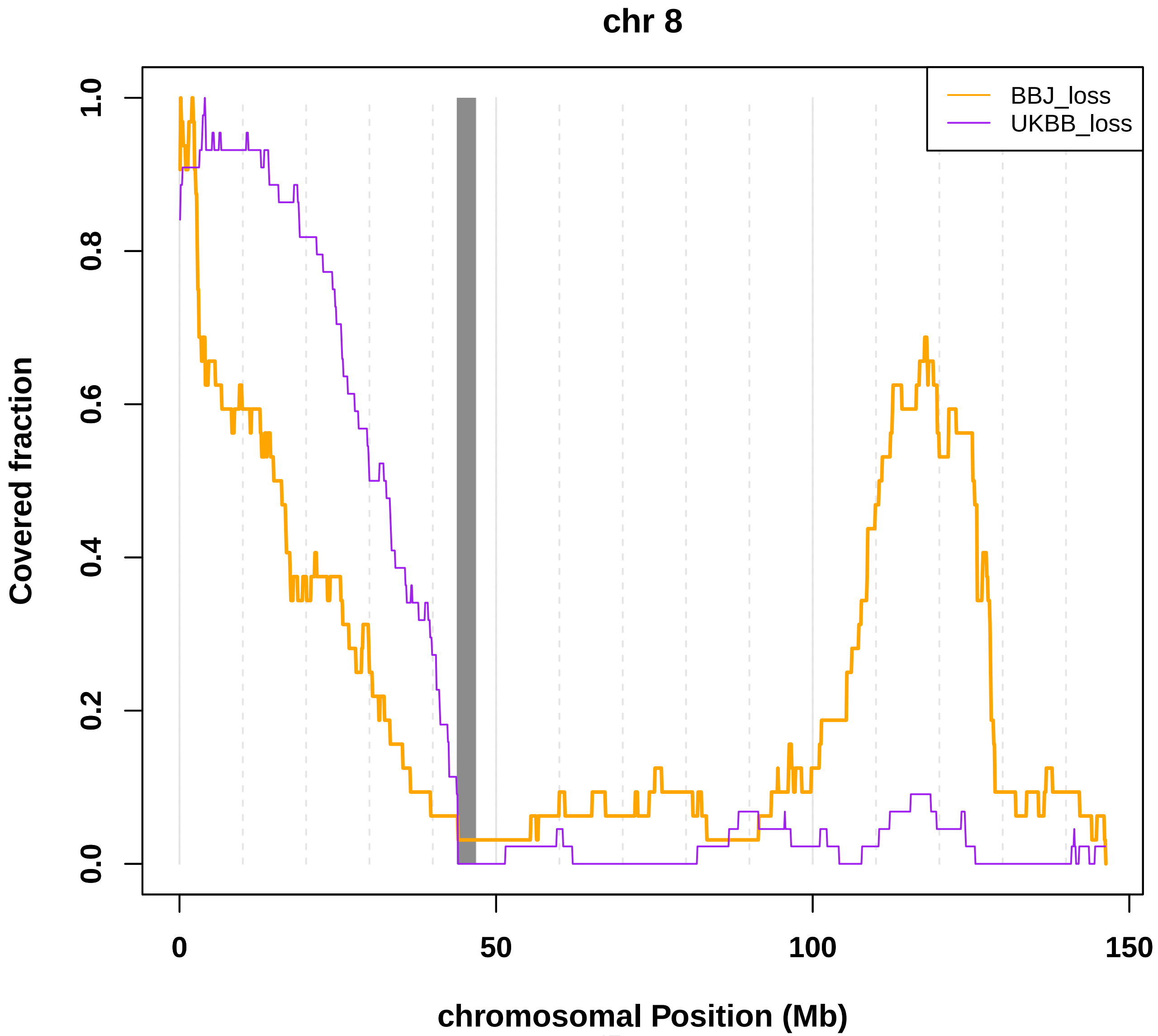

**Fig. S33 Coverage of mosaic loss in chromosome 9.**

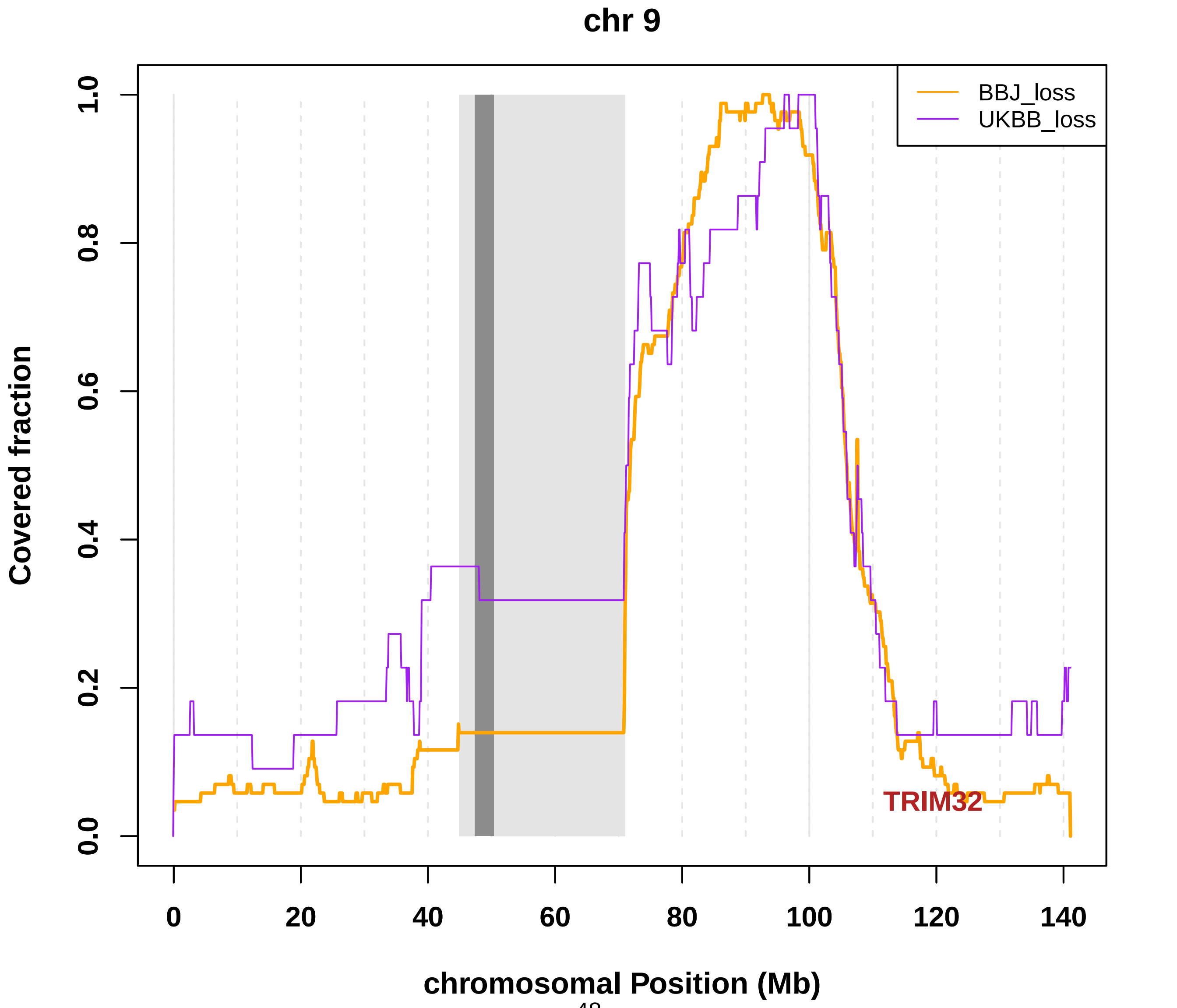

**Fig. S34 Coverage of mosaic loss in chromosome 10.**

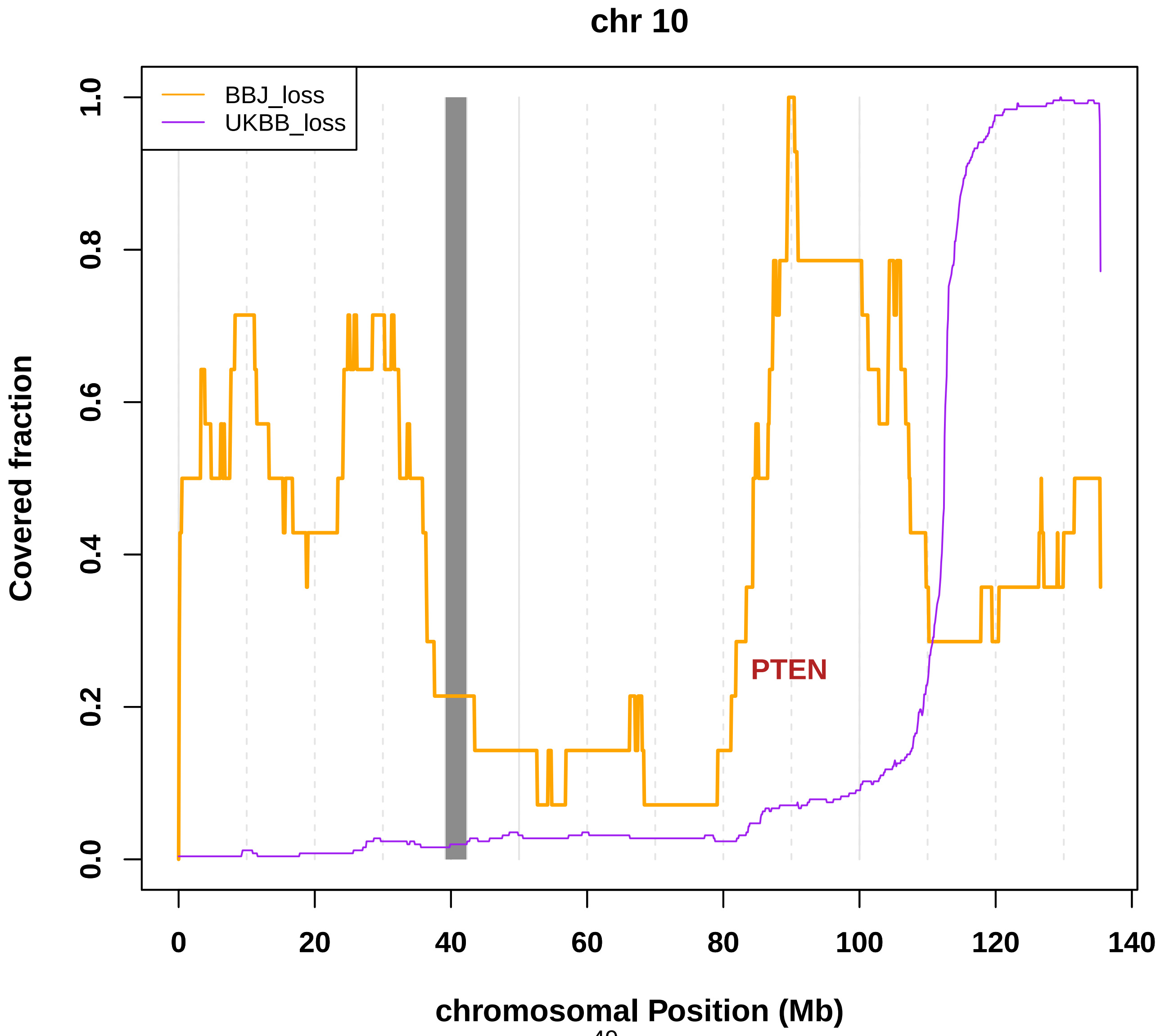

**Fig. S35 Coverage of mosaic loss in chromosome 11.**

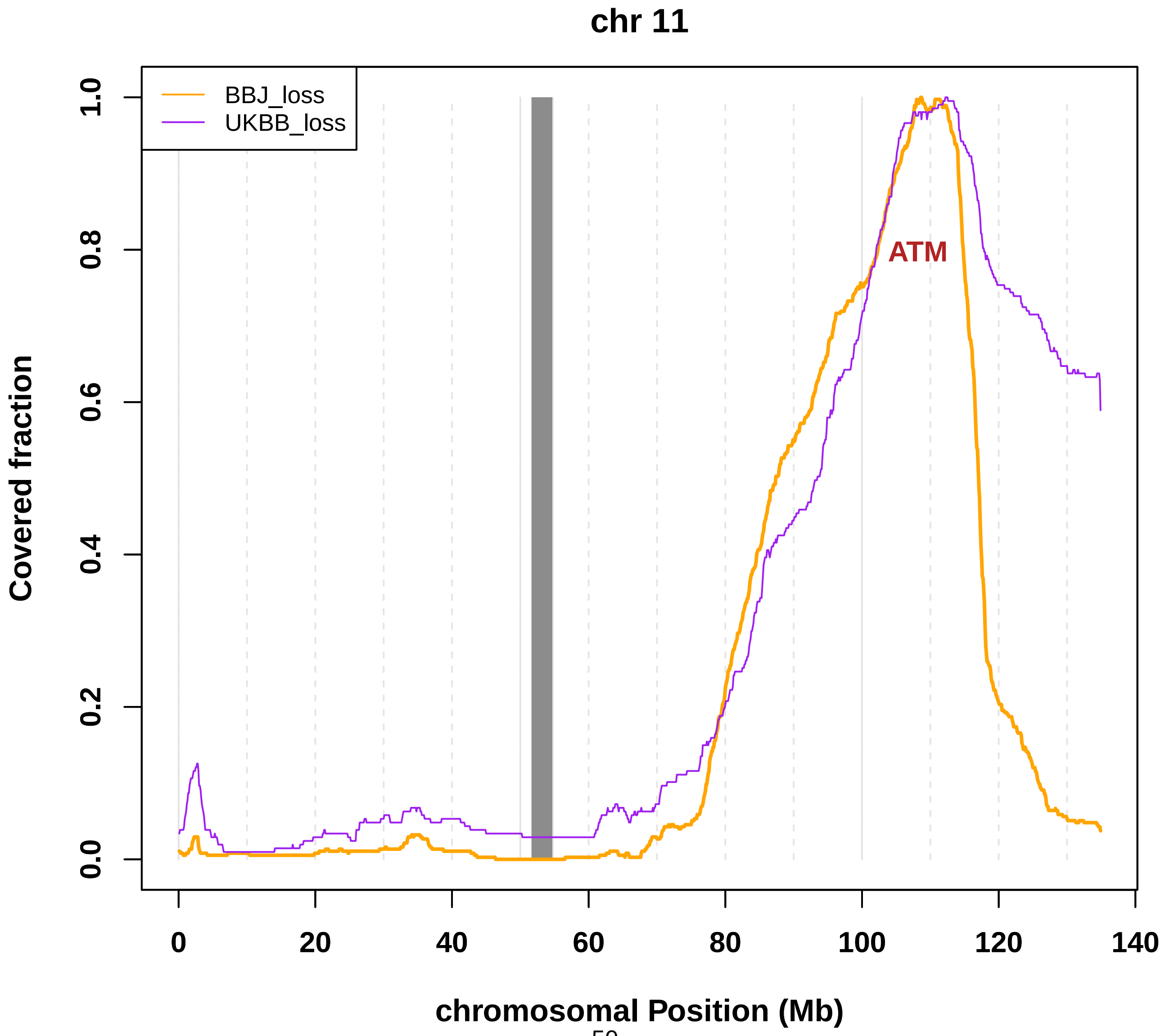

**Fig. S36 Coverage of mosaic loss in chromosome 12.**

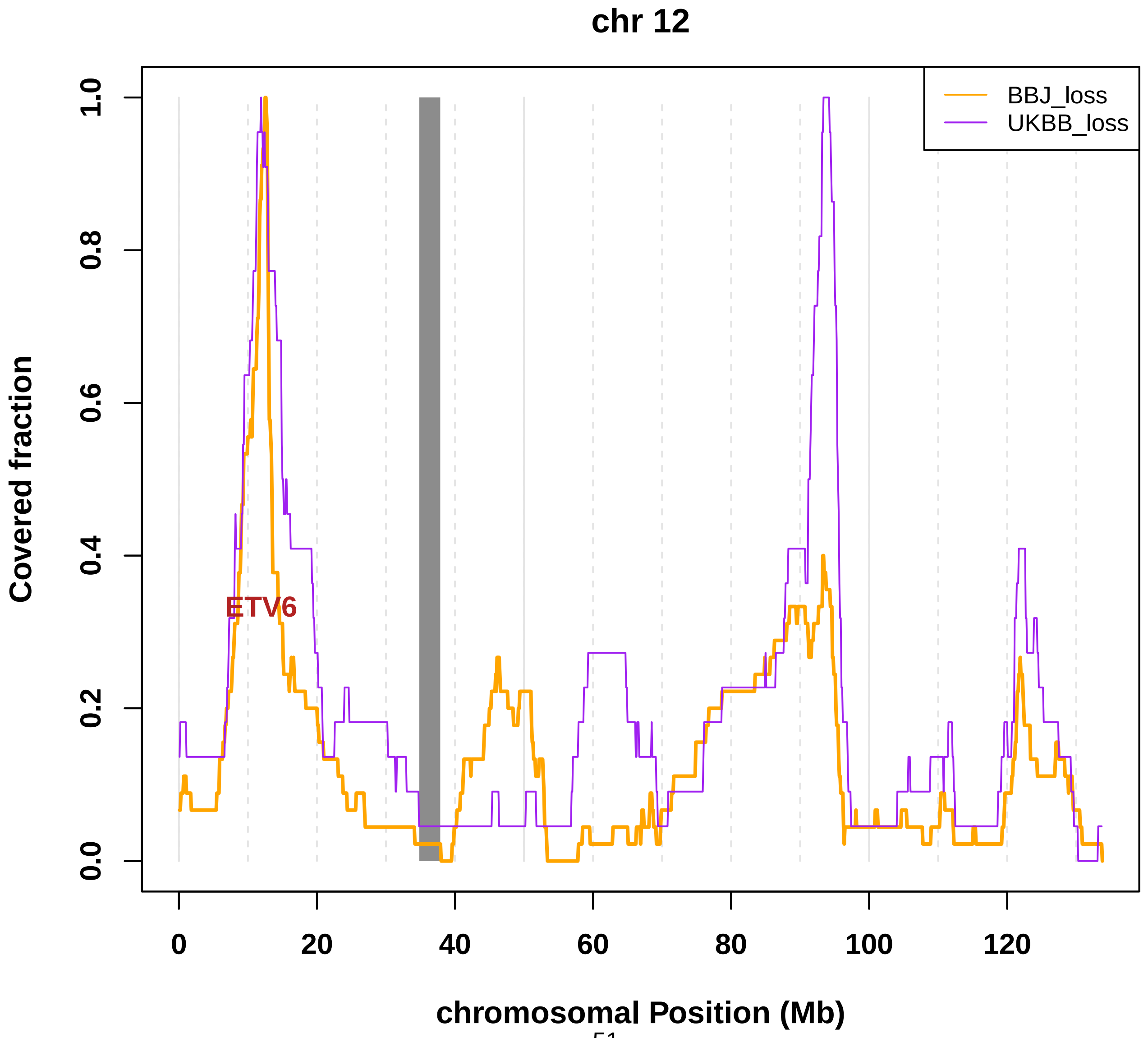

**Fig. S37 Coverage of mosaic loss in chromosome 13.**

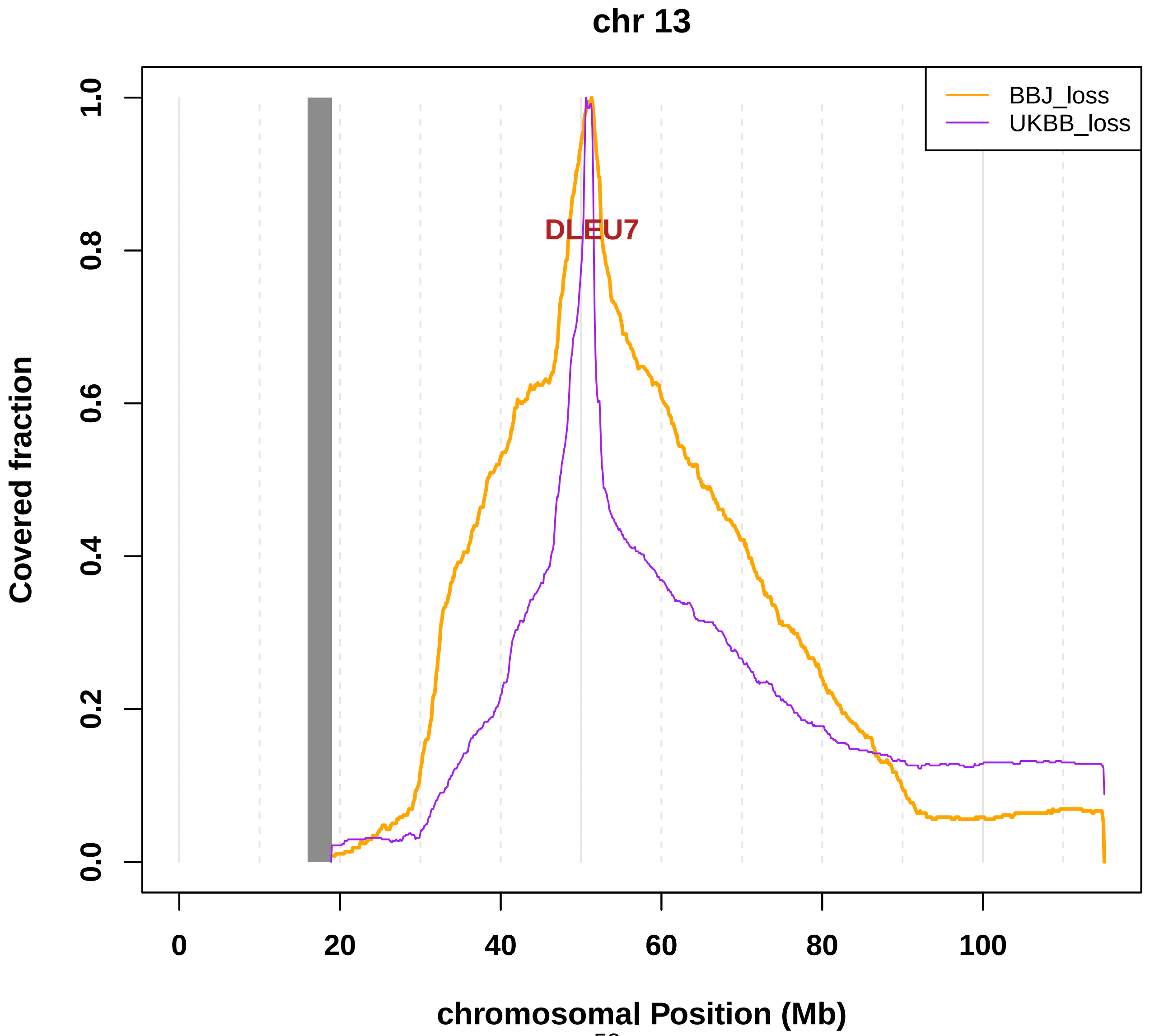

**Fig. S38 Coverage of mosaic loss in chromosome 14.**

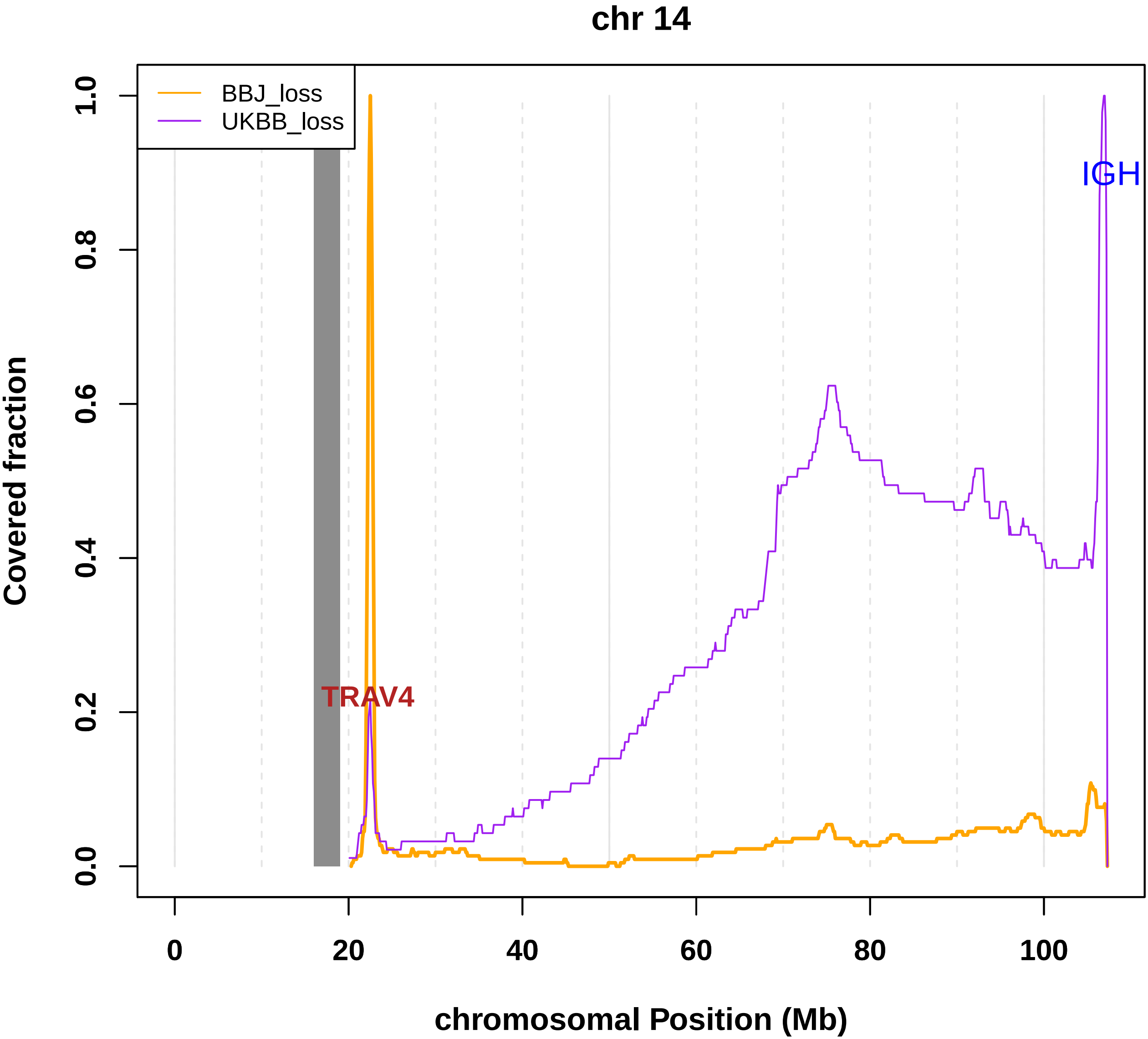

**Fig. S39 Coverage of mosaic loss in chromosome 15.**

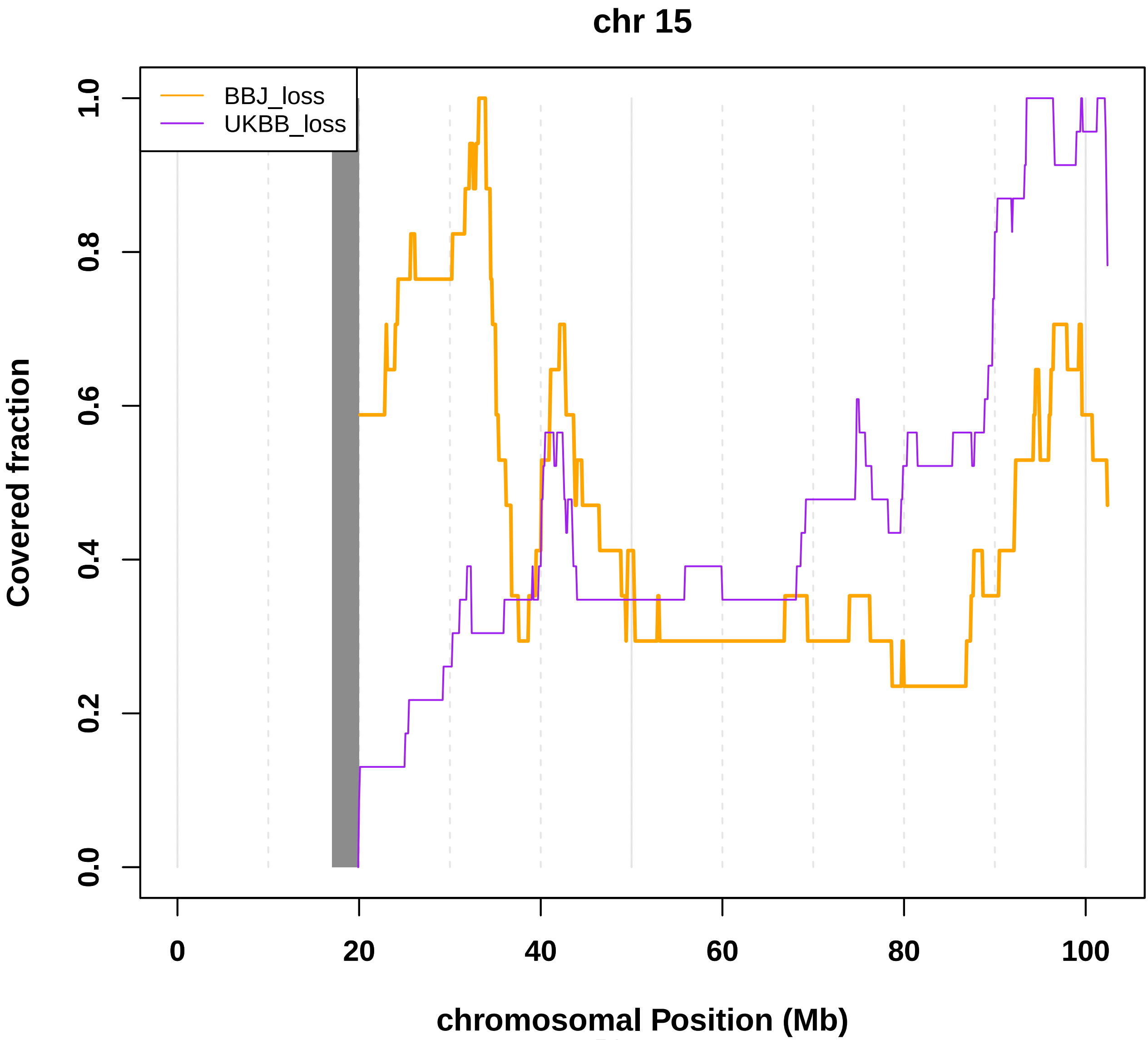

### Fig. S40 Coverage of mosaic loss in chromosome 16.

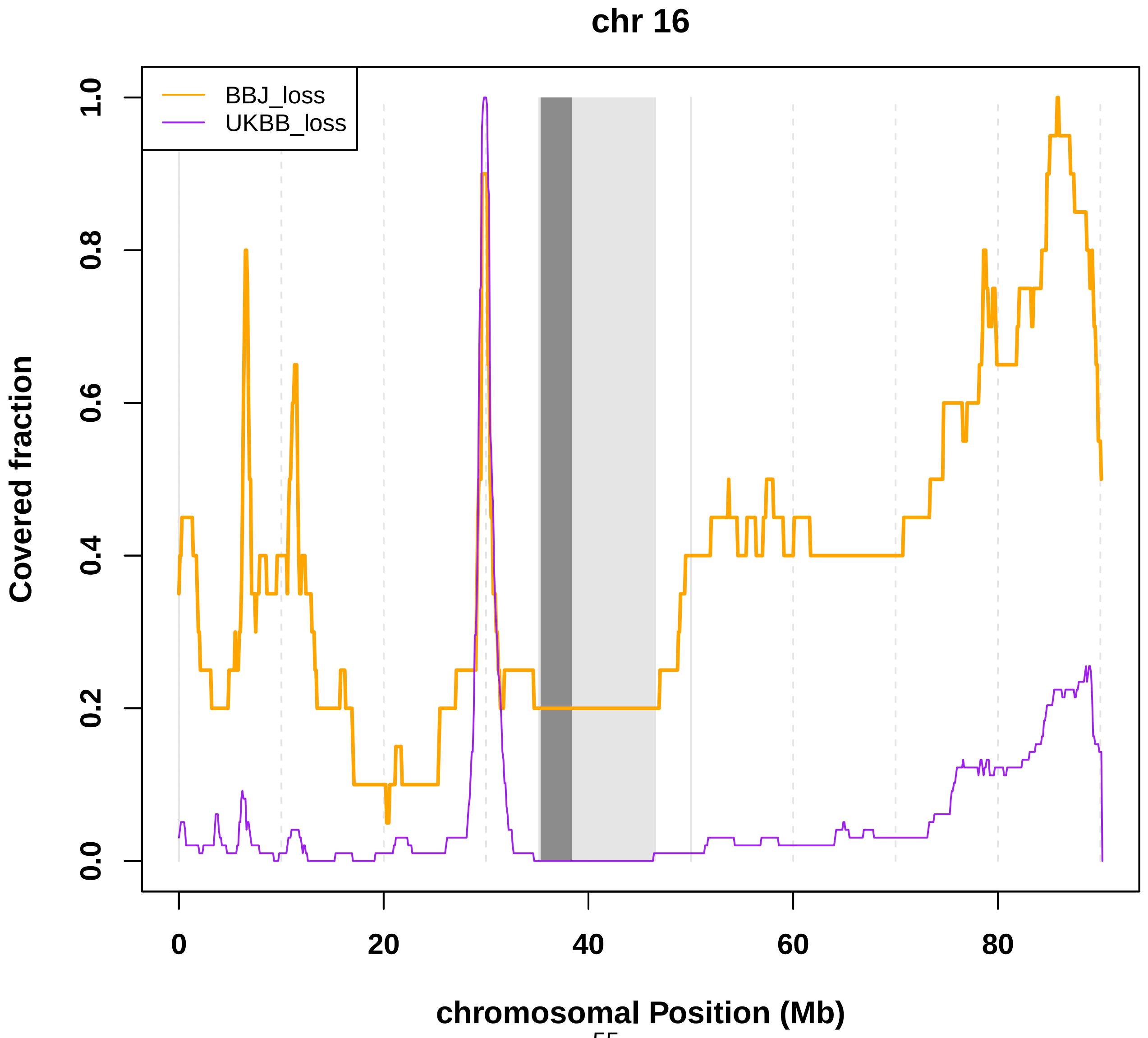

**Fig. S41 Coverage of mosaic loss in chromosome 17.**

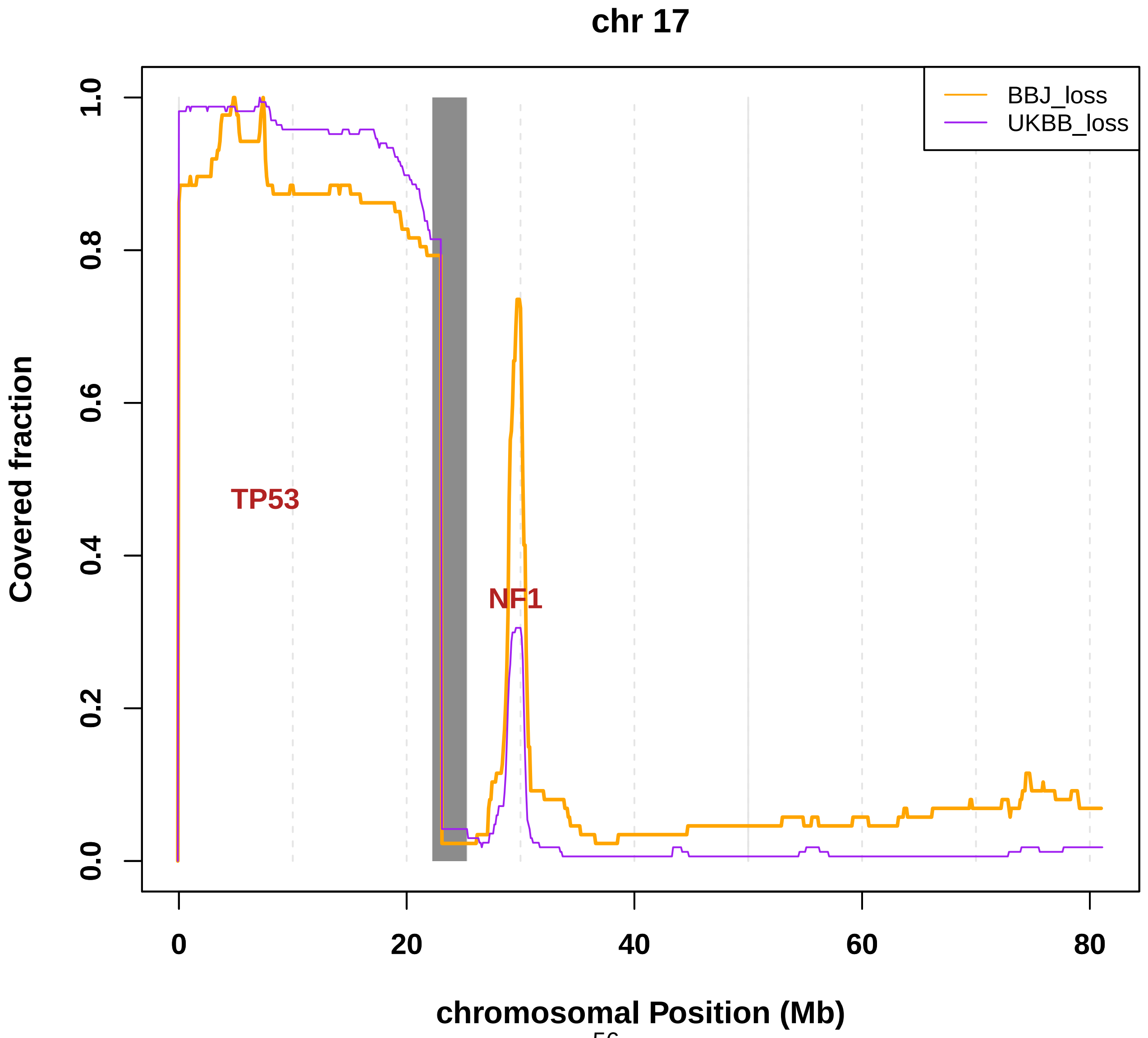

**Fig. S42 Coverage of mosaic loss in chromosome 18.**

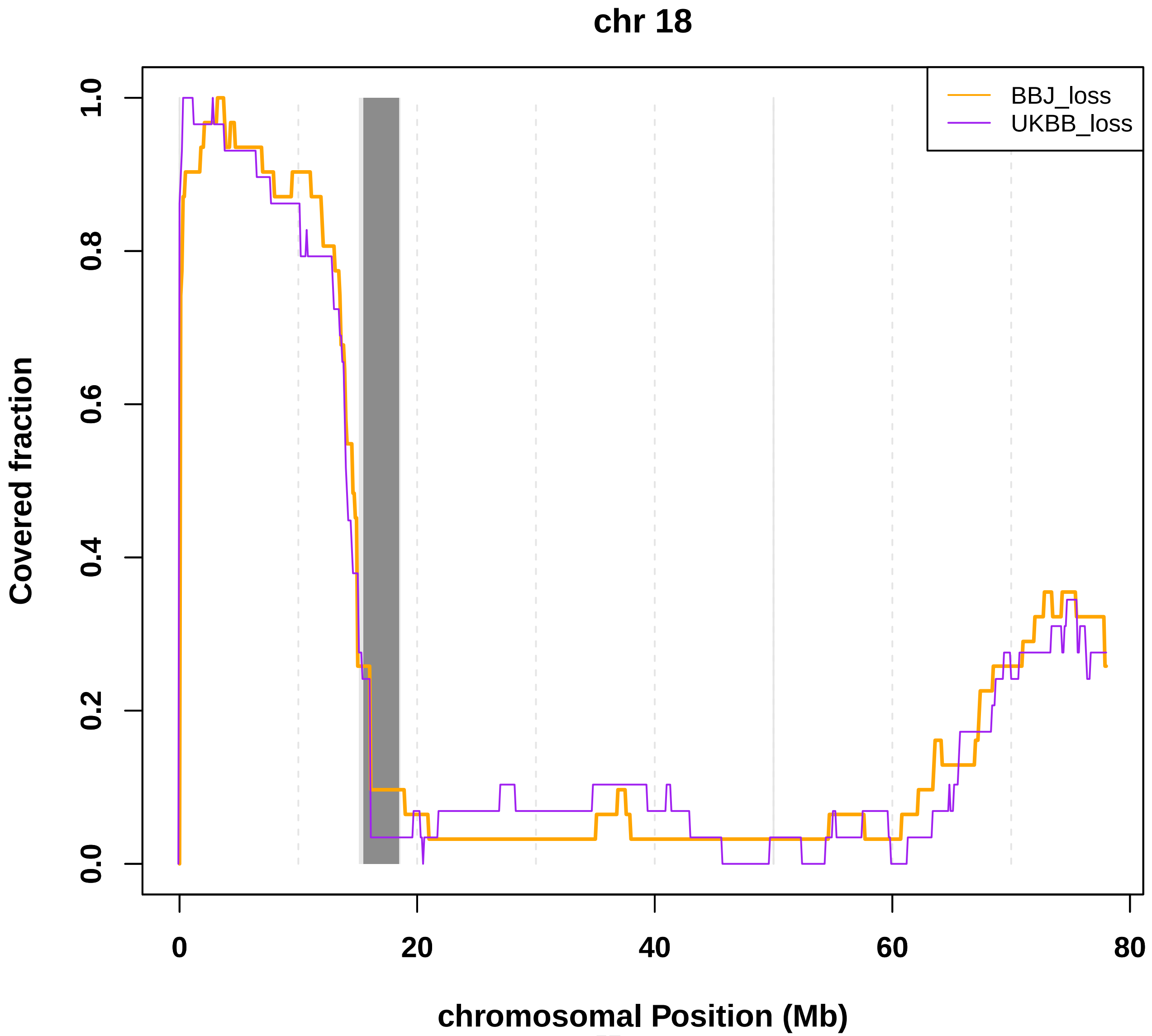

**Fig. S43 Coverage of mosaic loss in chromosome 19.**

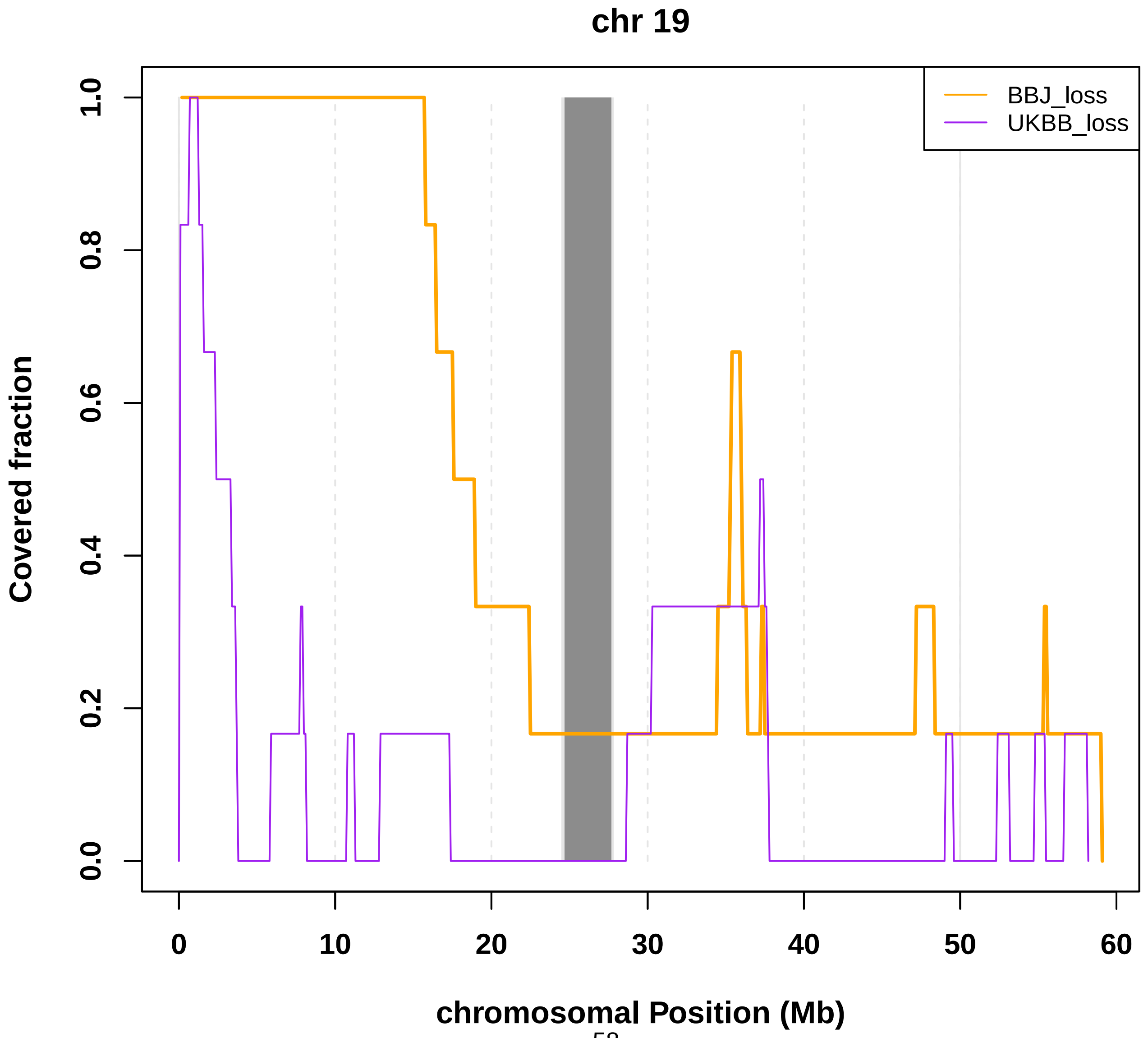

#### Fig. S44 Coverage of mosaic loss in chromosome 20.

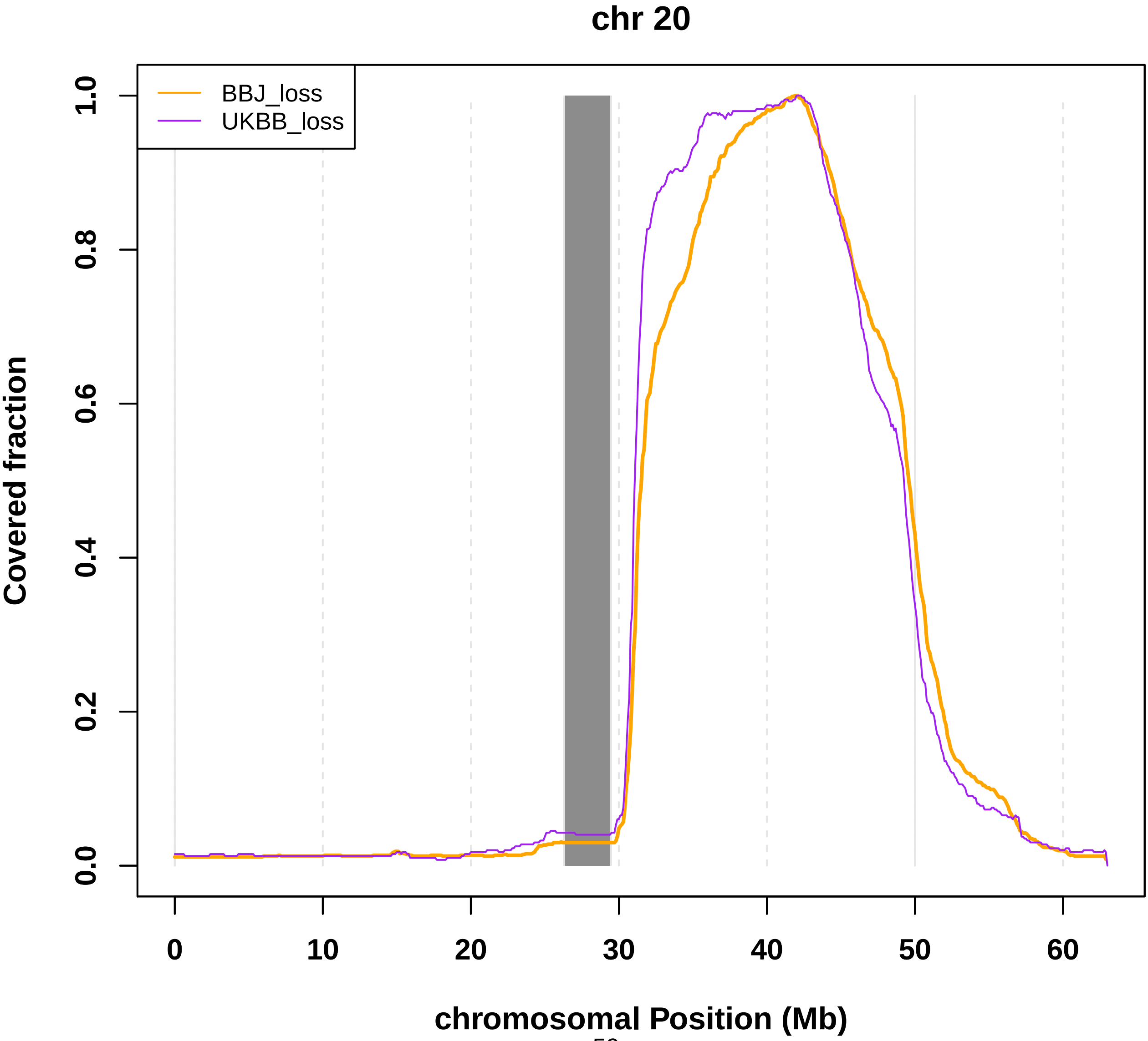

**Fig. S45 Coverage of mosaic loss in chromosome 21.**

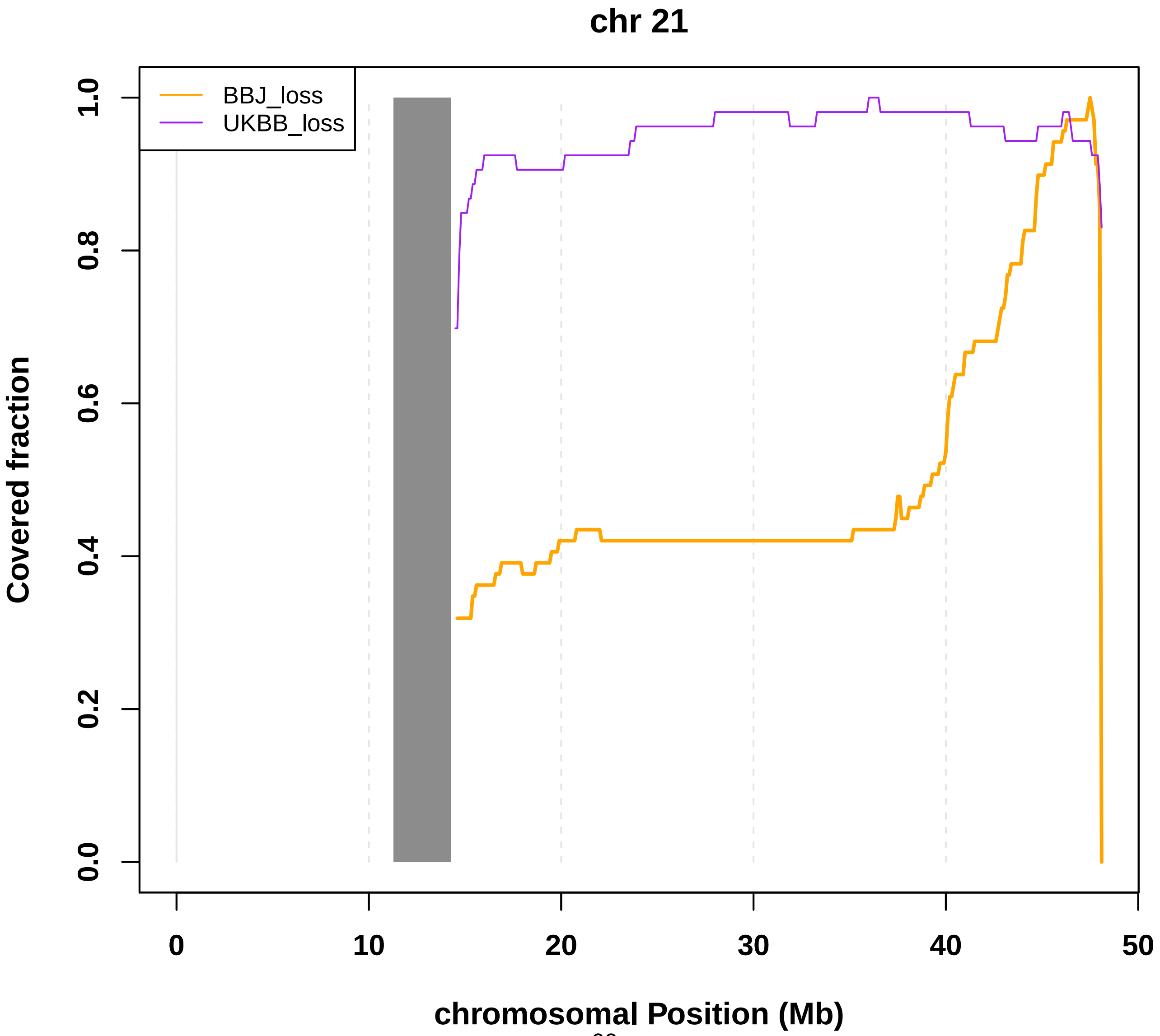

**Fig. S46 Coverage of mosaic loss in chromosome 22.**

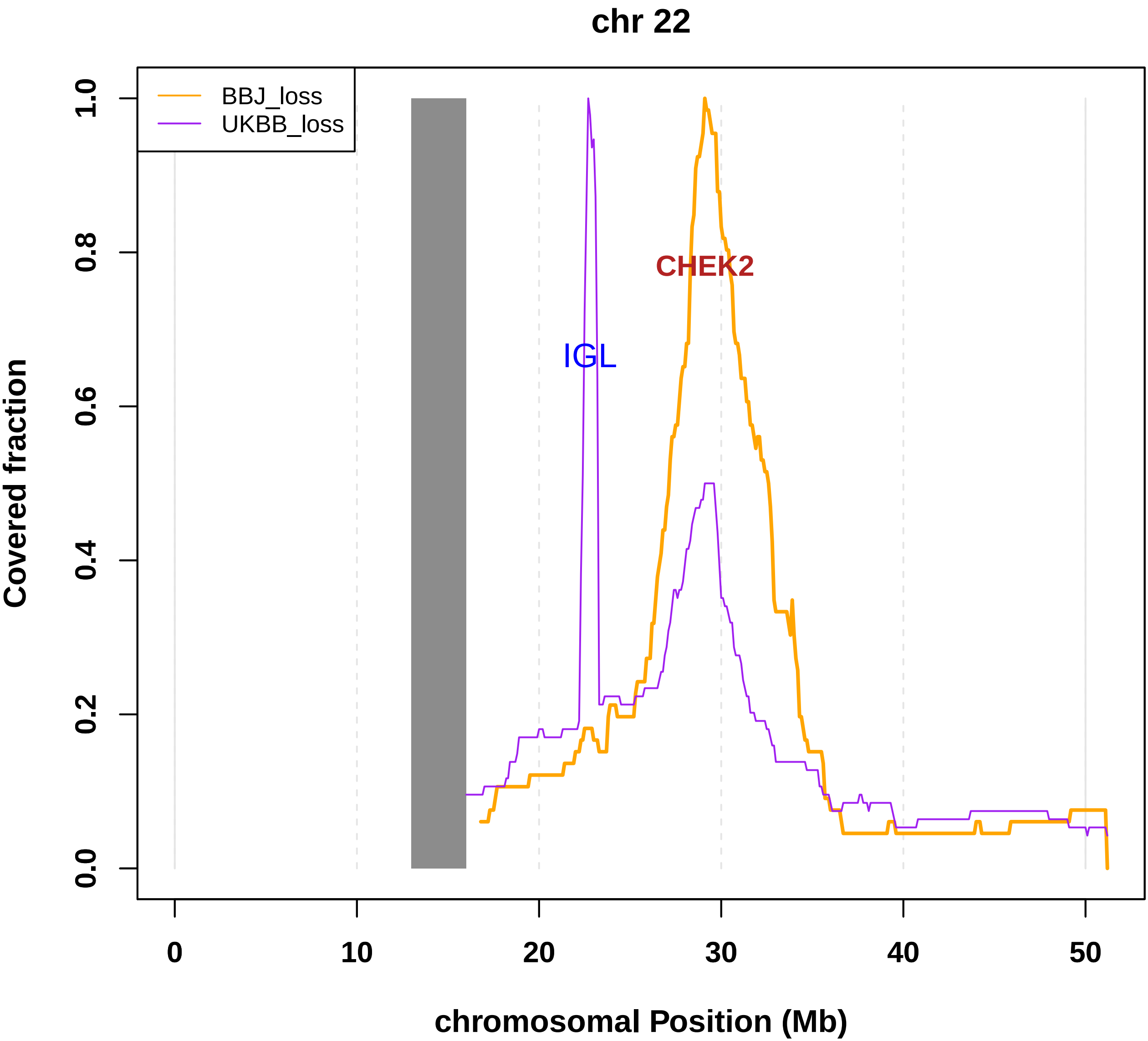

Fig S47. Comparable chromosomal coverage by heterozygous genotypes in BBJ and UKB.

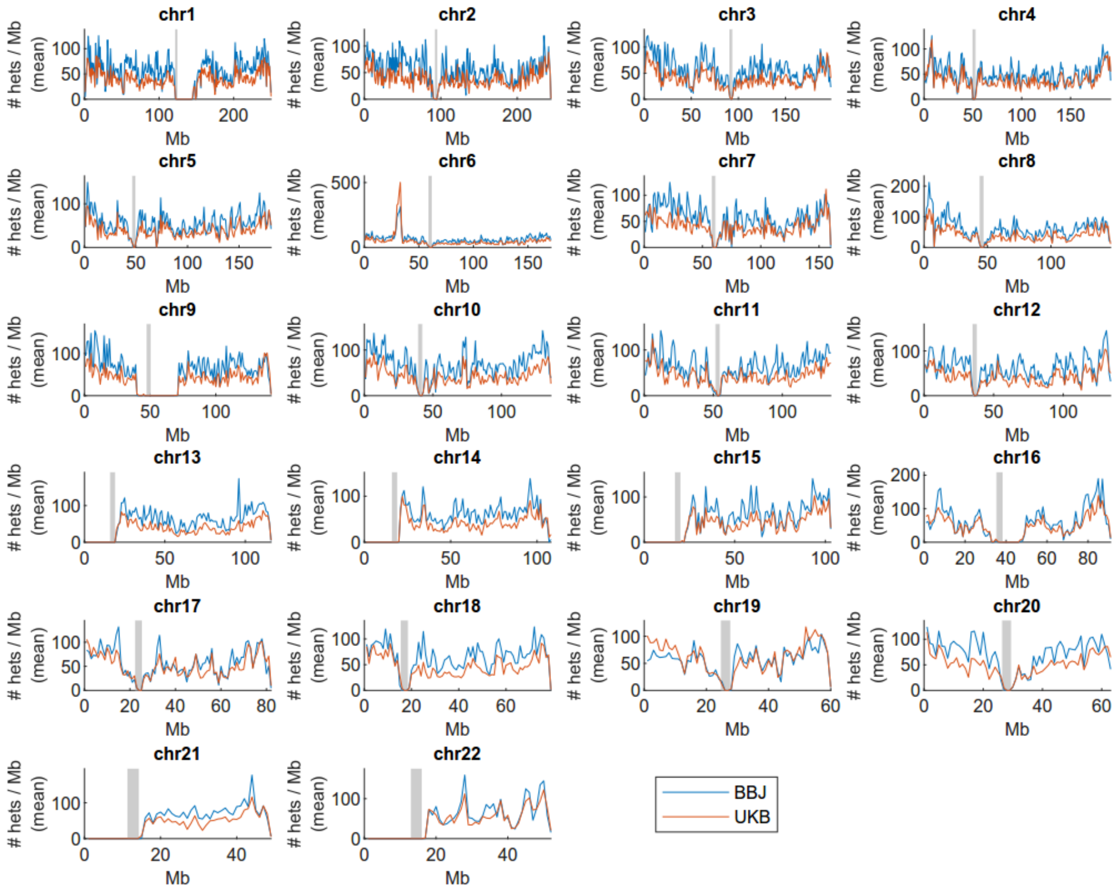

Average numbers of heterozygous genotyped sites (averaged across individuals) in each 1Mb region of the genome for the BioBank Japan and UK Biobank genotyping arrays.

Fig S48. Similar breakpoint distributions of CNN-LOH events in the BBJ and UKB.

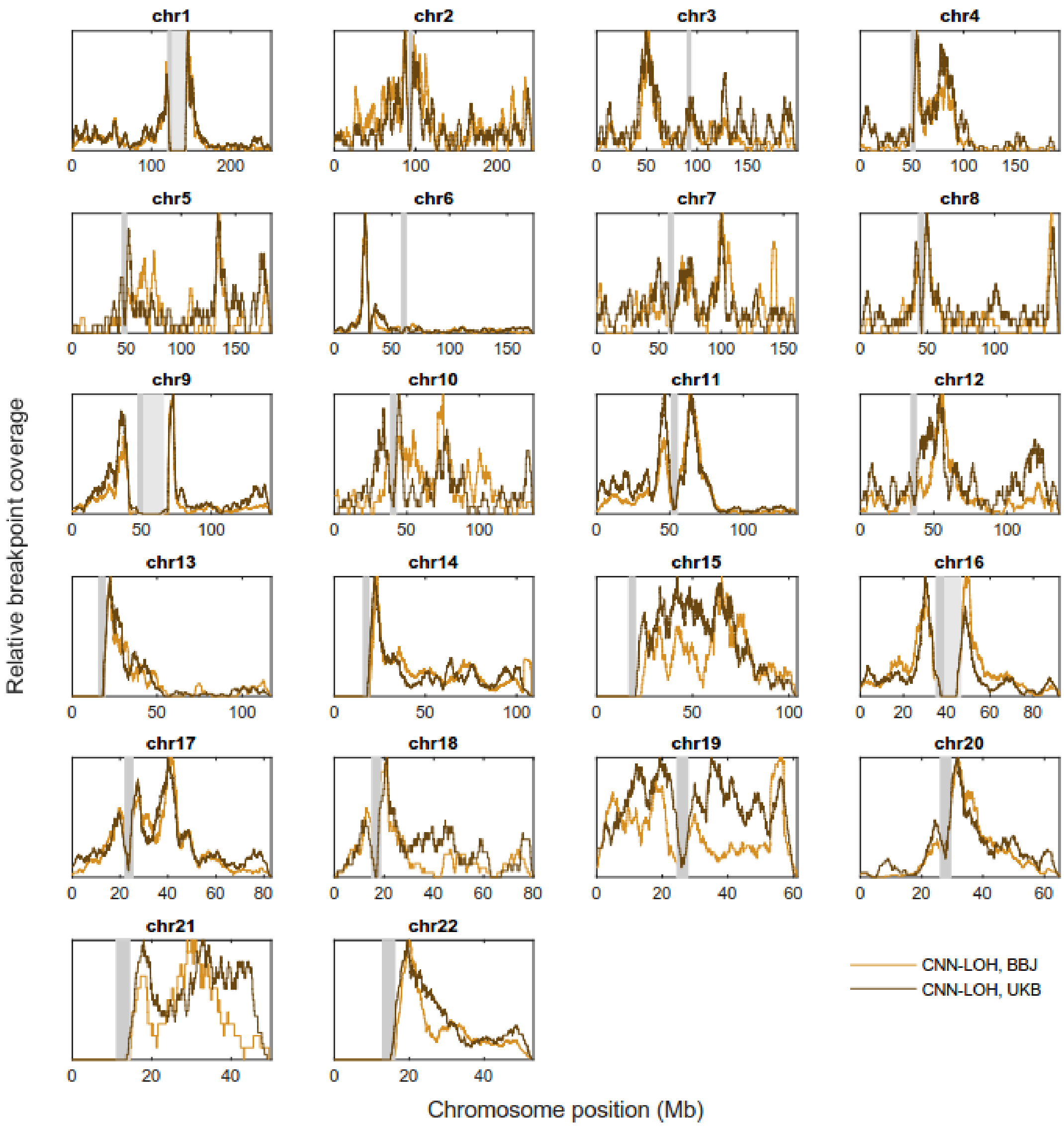

Relative frequencies of estimated CNN-LOH breakpoint locations in BioBank Japan and UK Biobank. Breakpoints were smoothed over +/-2Mb to allow plotting of frequency curves, which were rescaled to 1.

Fig S49. Quantile-quantile plots of mosaic events with significant associations demonstrating no inflation of association statistics.

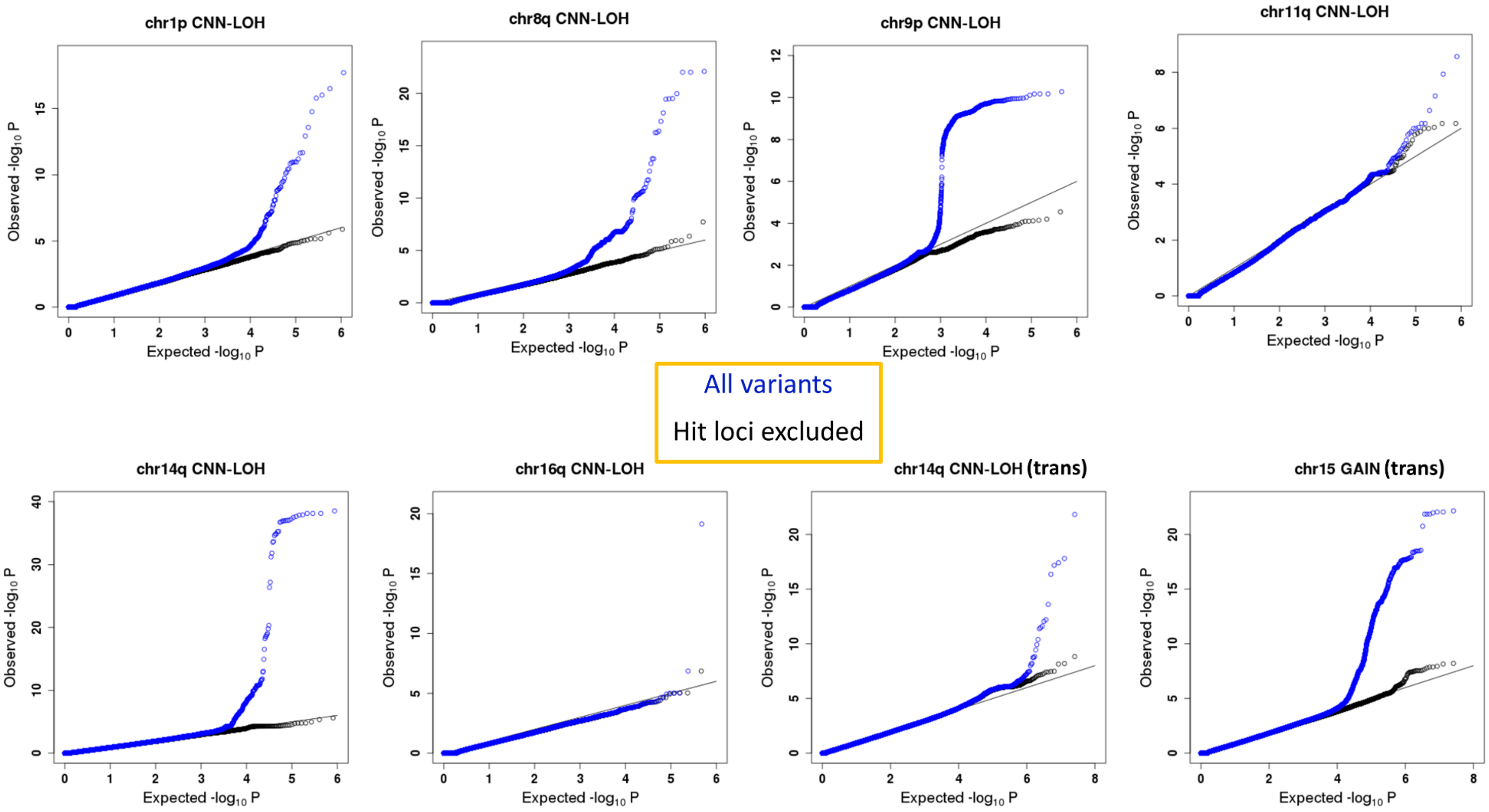

Quantile-quantile plots of results for mosaic events with significant associations are indicated. We defined as hit loci 42-49M at chr1 (1p CNN-LOH), 88-94M at chr8 (8q CNN-LOH), 92-96M at chr11 (11q CNN-LOH), 88-90M at chr16 (16q CNN-LOH), 23-26M and 100-103M at chr14 (cis association of 14q CNN-LOH), 4-6M at chr9 (9p CNN-LOH), 0-2M at chr5 (trans association of 14q CNN-LOH) and 1-3M at chr7 (trans association of chr15 gain).

Fig. S50. A local plot for the *MPL* region associated with chr1p CNN-LOH.

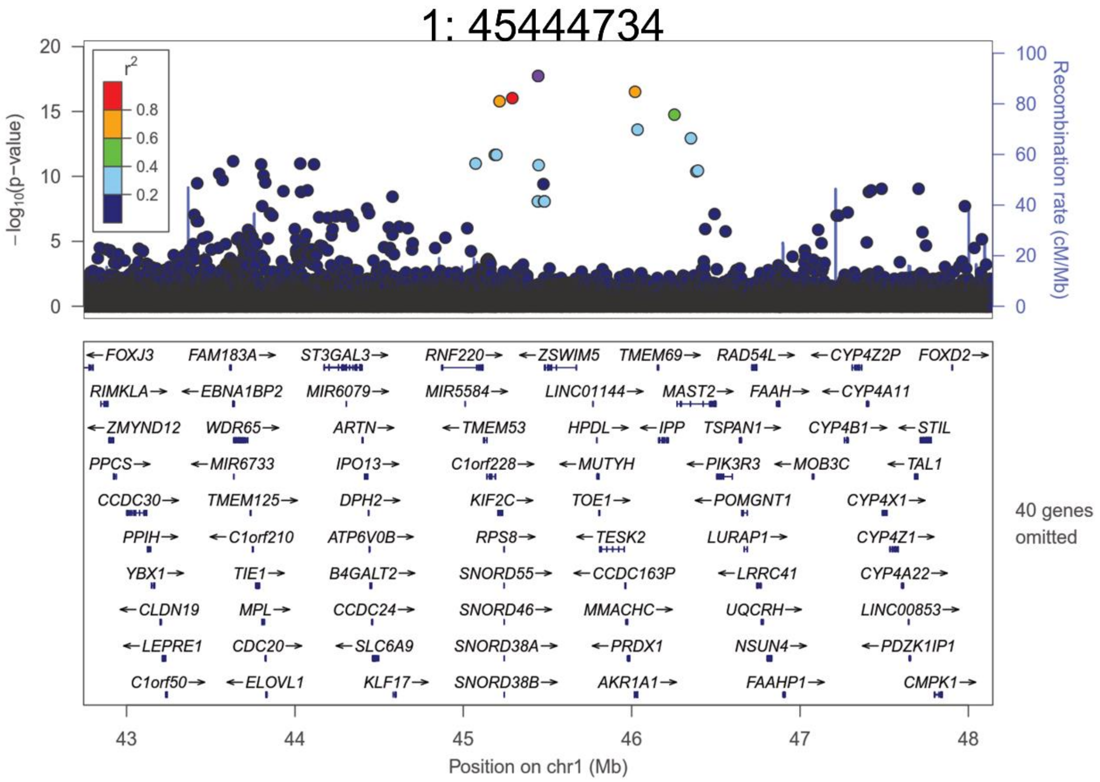

Associations of chromosome 1p CNN-LOH are shown according to chromosomal positions in a region containing the *MPL* locus.

Fig. S51. A local plot for the *JAK2* region associated with chr9p CNN-LOH.

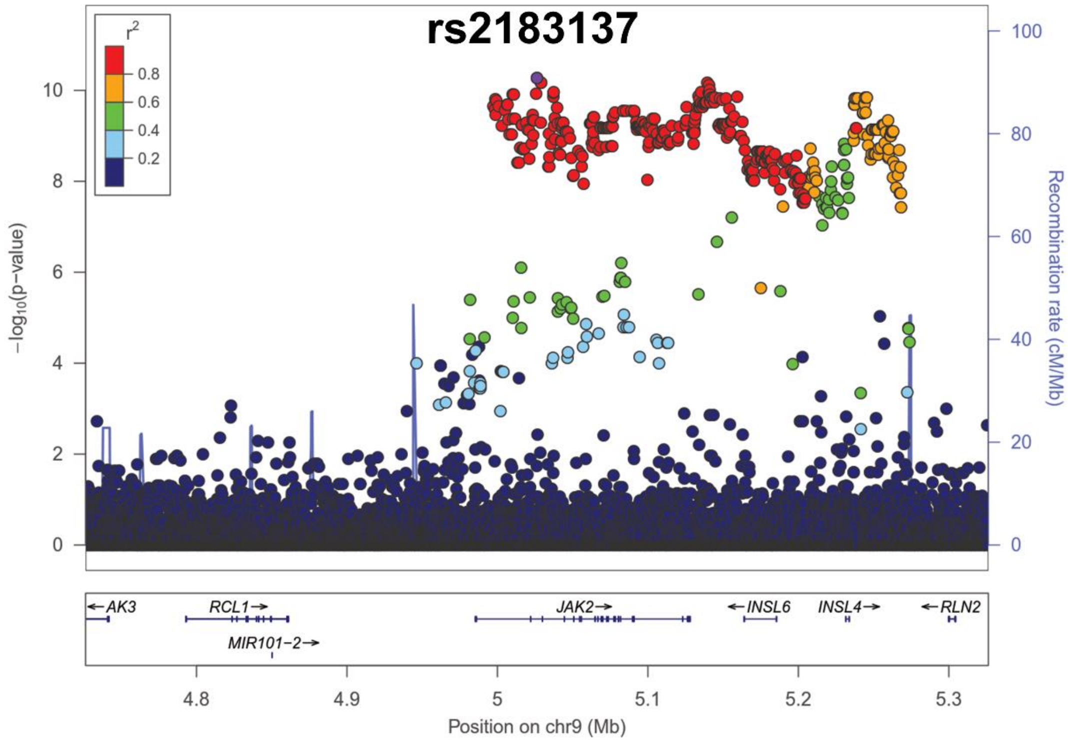

Associations of chromosome 9p CNN-LOH are shown according to chromosomal positions in a region containing the *JAK2* locus.

Fig. S52. Two *cis* and one *trans* association with chr14q CNN-LOH.

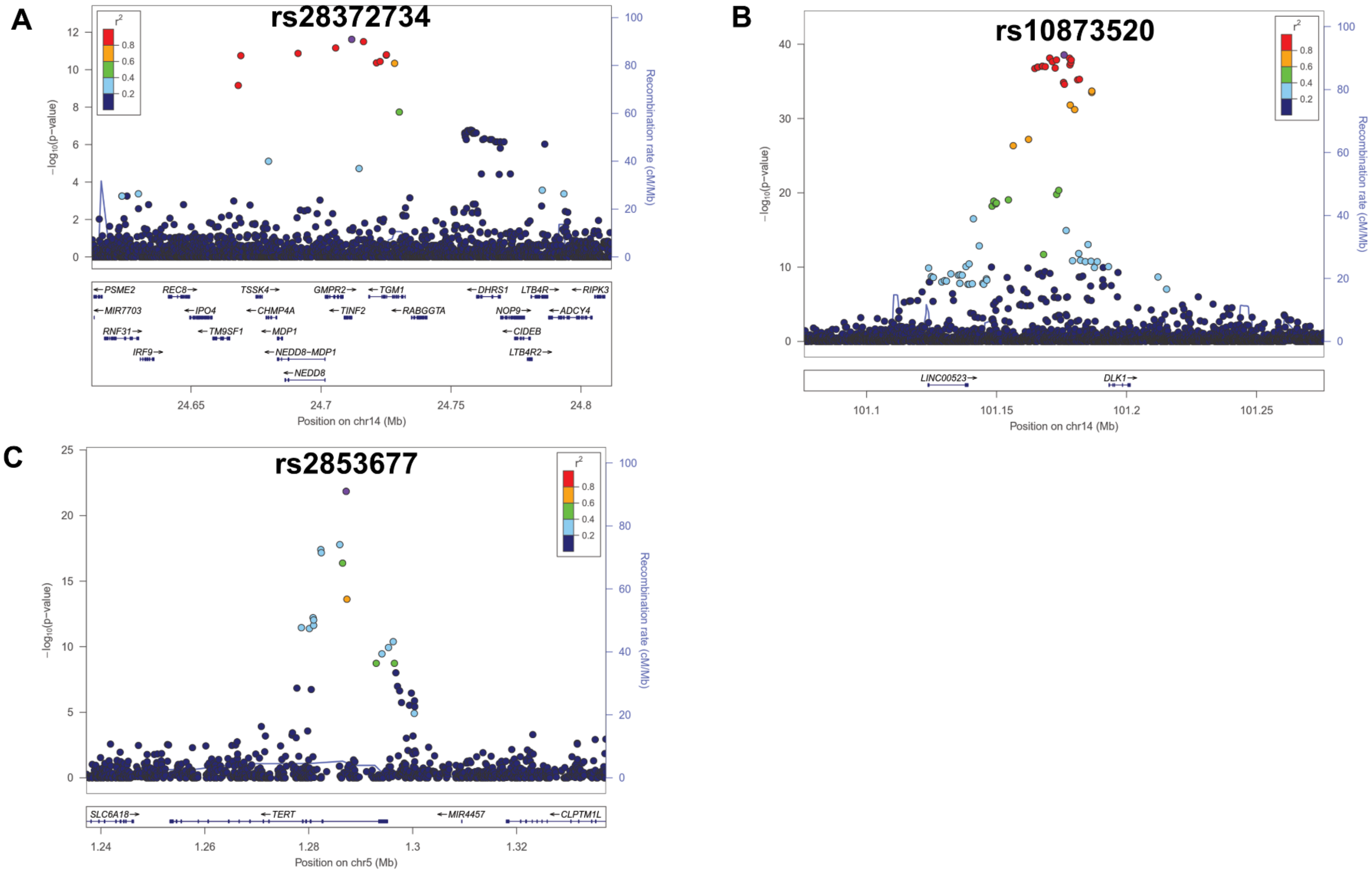

Associations of chromosome 14q CNN-LOH are shown according to chromosomal positions. Two *cis*-associations in the A) *NEDD8*/*TINF2* and B) *DLK1* regions, and C) one *trans*-association in the *TERT* region are indicated.

Fig S53 **Mortality risk conferred by mosaic chromosomal alterations.**

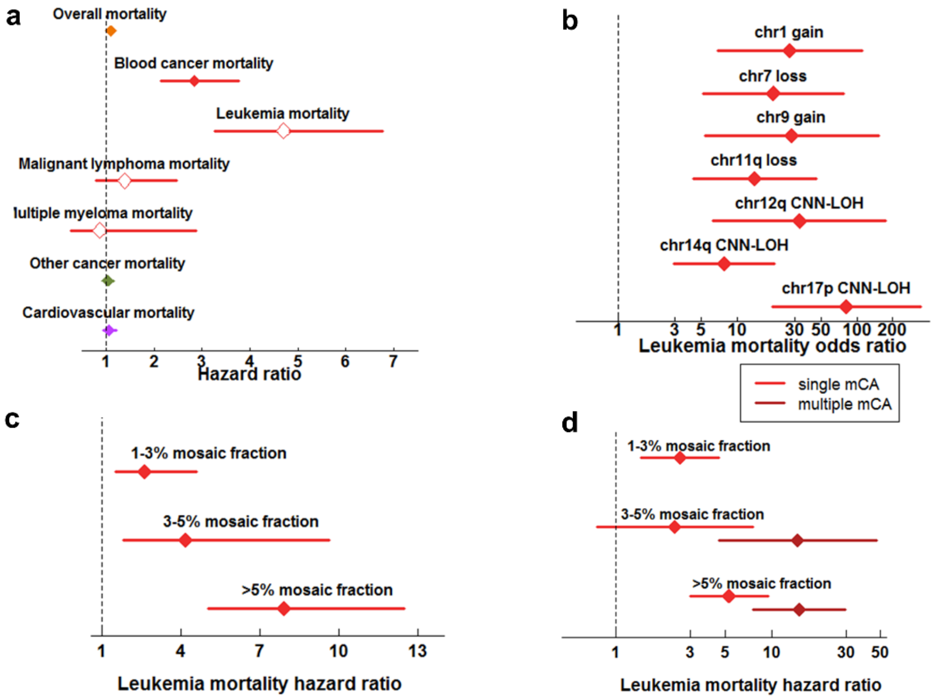

**a.** Risk of mortality from various causes conferred by presence of an mCA at >1% cell fraction. Leukemia, malignant lymphoma, and multiple myeloma are subdivisions of blood cancer. Cardiovascular mortality includes deaths from coronary artery disease (CAD) and ischemic stroke (IS).

**b.** Risk of leukemia mortality conferred by specific mCAs (grouped by chromosomal location and copy-number change) reaching Bonferroni significance.

**c.** Risk of leukemia mortality conferred by mosaic status stratified by mosaic cell fraction.

**d.** Risk of leukemia mortality conferred by mosaic status stratified by mosaic cell fraction and number of mosaic events detected (one vs. two or more).

All analyses were restricted to individuals with no previous cancer diagnosis and were corrected for age, sex, smoking status, and genotyping array (Methods). Error bars, 95% CIs. Numeric data are provided in Tables S19-22.

Fig S54. Co-occurrence patterns of mosaic events in different chromosomes in BBJ and UKB.

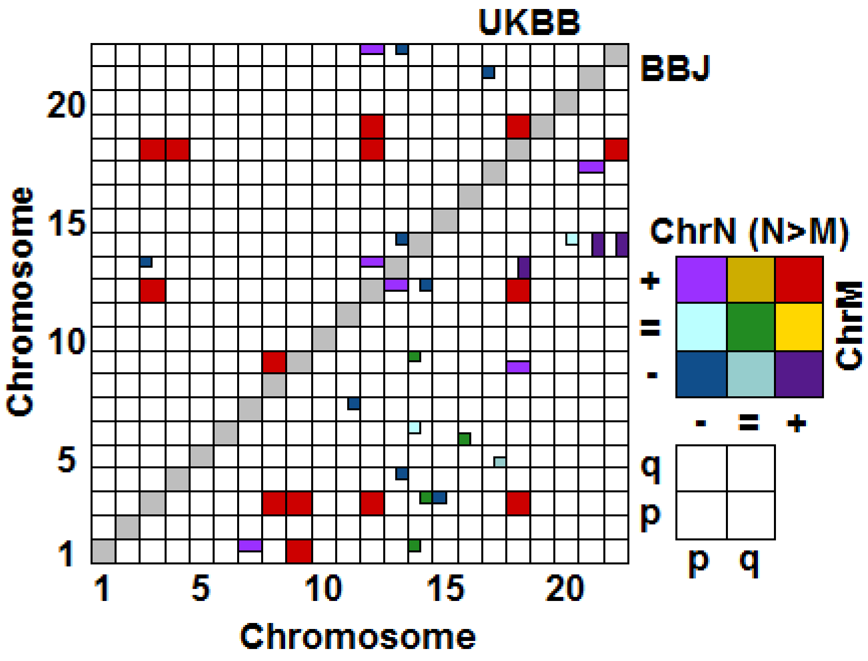

+, =, and – indicates gain, CNN-LOH and loss, respectively. UKB reports co-occurrence of the same chromosomal combinations (1) 13q- and 3p- or 3+ (2) 14q- and 13q- or 13q=, and (3) 22q- and 13q- or 13q=. If there are multiple co-occurrences in the same chromosomal combinations in a single population, we show one of two co-occurrence as a representative.

Fig. S55. Functional analyses suggesting alteration of gene expression in subjects carrying risk variants of *MPL* and *MRE11*.

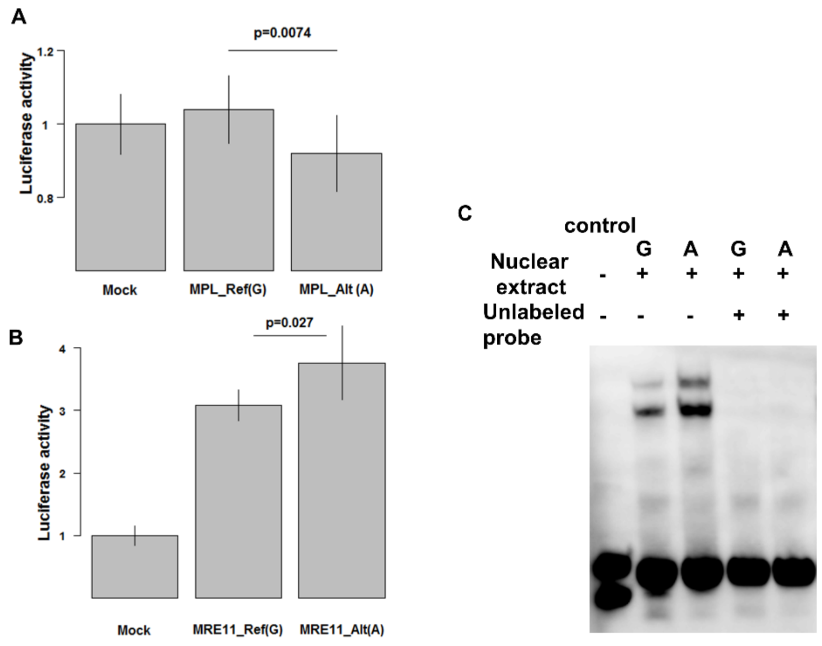

A.The *MPL* variant is suggestively associated with decreased expression of *MPL*.
A result of luciferase assay is indicated with the use of synthesized oligonucleotide centering the associated *MPL* variant.

B and C. *MRE11* variant is suggested to be associated with increased expression of *MRE11*.
A result of luciferase assay is indicated with the use of synthesized oligonucleotide centering the associated *MRE11* variant in B. A result of EMSA is indicated in C.

Table S1. The 179,417 subjects used to call mosaic events in the current study.

| | Set1 | Set2 | Set3 |
|---|---|---|---|
| Samples | 34,256 | 111,422 | 33,739 |
| Arrays | OmniExpressExome v1.0 | OmniExpressExome v1.2 | OmniExpress v1.0 and HumanExome v1.0, 1.1 |
| Age | 69.4+/-10.3 | 61.6+/-14.8 | 59.9+/-15.4 |
| Female ratio | 0.46 | 0.46 | 0.45 |

Table. S2. Chromosomal distribution of classified mosaic events.

| CHR | LOSS | p-arm LOSS | q-arm LOSS | CNN-LOH | p-arm CNN-LOH | q-arm CNN-LOH | GAIN |
|---|---|---|---|---|---|---|---|
| chr1 | 156 | 123 | 28 | 1489 | 813 | 652 | 221 |
| chr2 | 293 | 219 | 69 | 388 | 163 | 210 | 52 |
| chr3 | 175 | 125 | 47 | 270 | 158 | 105 | 91 |
| chr4 | 212 | 34 | 164 | 260 | 16 | 205 | 24 |
| chr5 | 460 | 12 | 443 | 102 | 7 | 93 | 86 |
| chr6 | 413 | 22 | 377 | 485 | 380 | 100 | 63 |
| chr7 | 263 | 70 | 179 | 152 | 33 | 104 | 43 |
| chr8 | 95 | 50 | 44 | 92 | 20 | 67 | 370 |
| chr9 | 169 | 19 | 136 | 690 | 317 | 366 | 198 |
| chr10 | 56 | 22 | 31 | 163 | 28 | 134 | 21 |
| chr11 | 466 | 35 | 428 | 907 | 362 | 535 | 23 |
| chr12 | 123 | 51 | 68 | 254 | 33 | 218 | 122 |
| chr13 | 453 | 0 | 451 | 224 | 0 | 157 | 13 |
| chr14 | 335 | 0 | 332 | 2688 | 0 | 2478 | 108 |
| chr15 | 48 | 0 | 47 | 225 | 0 | 199 | 886 |
| chr16 | 92 | 57 | 29 | 555 | 267 | 284 | 9 |
| chr17 | 177 | 95 | 80 | 567 | 133 | 428 | 95 |
| chr18 | 50 | 32 | 15 | 99 | 22 | 75 | 195 |
| chr19 | 12 | 5 | 6 | 217 | 112 | 105 | 16 |
| chr20 | 1029 | 16 | 984 | 332 | 20 | 304 | 8 |
| chr21 | 73 | 0 | 73 | 57 | 0 | 37 | 1098 |
| chr22 | 83 | 0 | 83 | 215 | 1 | 186 | 302 |
| Sum | 5233 | 987 | 4114 | 10431 | 2885 | 7042 | 4044 |

Since p and q indicates whether an event extends to a telomere, the sum of p and q mosaics is not always the same as LOSS or CNN-LOH in the corresponding chromosomes.

Table. S3. Number of mosaic events in each individual.

| Mosaic N | Individual |
|---|---|
| 0 | 151507 |
| 1 | 23754 |
| 2 | 3376 |
| 3 | 554 |
| 4 | 144 |
| 5 | 39 |
| 6 | 23 |
| 7 | 7 |
| 8 | 3 |
| 9 | 6 |
| 10 | 1 |
| 11 | 2 |
| 14 | 1 |

Table S4. Mosaic detection rate across batches

| | N | P | OR (95%CI) |
|---|---|---|---|
| Set1 | 32,959 | 0.33 | 1.02 (0.98-1.05) |
| Set2 | 107,767 | Reference | 1 |
| Set3 | 32,873 | $1.1x10^{-14}$ | 0.86 (0.83-0.9) |

OR: odds ratio, CI: confidence interval

OR was calculated by logistic regression with mosaic detection as a dependent variable and age, sex, smoking and arrays as independent variables.

Since we conducted association studies for 173,599 subjects after exclusion of samples based on kinship and evidence of contamination, the number of subjects in the three sets were slightly different from those in Table S1.

Table. S5. Associations between disease status and mosaic events

| Disease group | Disease | N | P | OR (95%CI) |
|---|---|---|---|---|
| Malignant tumors | Lung cancer | 3397 | 0.91 | 0.99 (0.91-1.09) |
| | Esophageal cancer | 1141 | 0.85 | 1.02 (0.86-1.19) |
| | Gastric cancer | 5613 | $1.5x10^{-5}$ | 1.17 (1.09-1.26) |
| | Colorectal cancer | 5909 | 0.014 | 1.09 (1.02-1.17) |
| | Liver cancer | 1334 | 0.25 | 1.09 (0.94-1.25) |
| | Pancreas cancer | 279 | 0.82 | 1.04 (0.75-1.44) |
| | Gallbladder/Cholangiocarcinoma | 224 | 0.69 | 0.93 (0.65-1.33) |
| | Prostate cancer | 4519 | 0.42 | 0.97 (0.89-1.05) |
| | Breast cancer | 4933 | 0.74 | 0.98 (0.89-1.09) |
| | Cervical cancer | 546 | 0.12 | 1.25 (0.94-1.66) |
| | Uterine cancer | 917 | 0.31 | 0.88 (0.7-1.12) |
| | Ovarian cancer | 663 | 0.81 | 1.03 (0.79-1.35) |
| | Hematopoietic tumor | 1075 | $2.0x10^{-17}$ | 1.93 (1.66-2.24) |
| Cerabral diseases | Cerebral infarction | 14862 | 0.43 | 1.02 (0.97-1.06) |
| | Cerebral aneurysm | 2473 | 0.012 | 0.85 (0.75-0.97) |
| | Epilepsy | 1908 | 0.36 | 0.93 (0.78-1.09) |
| Respiratory diseases | Bronchial asthma | 7304 | 0.26 | 0.96 (0.89-1.03) |
| | Pulmonary tuberculosis | 386 | 0.20 | 1.17 (0.92-1.5) |
| | Chronic obstructive pulmonary disease | 2525 | 0.49 | 1.03 (0.94-1.14) |
| | Interstitial lung disease/Pulmonary fibrosis | 450 | 0.77 | 0.96 (0.76-1.23) |
| Cardiovascular diseases | Myocardial infarction | 11881 | 0.015 | 0.94 (0.89-0.99) |
| | Unstable angina | 3875 | 0.43 | 0.97 (0.89-1.05) |
| | Stable angina | 13402 | 0.18 | 1.03 (0.99-1.08) |
| | Arrhythmia | 14447 | 0.72 | 0.99 (0.95-1.04) |
| | Heart failure | 6799 | 0.31 | 0.97 (0.91-1.03) |
| | Peripheral arterial diseases | 2424 | 0.0054 | 1.15 (1.04-1.27) |
| Liver diseases | Chronic hepatitis B | 1197 | 0.028 | 1.21 (1.02-1.43) |

| | | | | |
|---|---|---|---|---|
| | Chronic hepatitis C | 5144 | 0.39 | 1.03 (0.96-1.12) |
| | Liver cirrhosis | 1413 | 0.75 | 1.02 (0.89-1.18) |
| Urologic diseases | Nephrotic syndrome | 863 | 0.91 | 0.99 (0.79-1.24) |
| | Urolithiasis | 5800 | 0.0018 | 0.87 (0.8-0.95) |
| Metabolic diseases | Osteoporosis | 5906 | 0.069 | 0.94 (0.87-1.01) |
| | Diabetes mellitus | 35856 | 0.23 | 0.98 (0.95-1.01) |
| | Dyslipidemia | 39459 | 0.18 | 1.02 (0.99-1.06) |
| Endocrine diseases | Graves' disease | 1973 | $1.3x10^{-7}$ | 0.58 (0.47-0.71) |
| Connective tissue diseases | Rheumatoid arthritis | 3777 | 0.00015 | 0.82 (0.74-0.91) |
| Allergic diseases | Hay fever | 5062 | 0.020 | 0.88 (0.79-0.98) |
| Dermatologic diseases | Drug eruption | 213 | 0.16 | 1.28 (0.9-1.83) |
| | Atopic dermatitis | 2507 | 0.45 | 0.93 (0.77-1.12) |
| | Keloid | 735 | 0.47 | 1.09 (0.86-1.4) |
| Gynecologic diseases | Uterine fibroid | 5473 | 0.69 | 1.02 (0.91-1.15) |
| | Endometriosis | 656 | 0.43 | 0.87 (0.61-1.23) |
| Pediatric diseases | Febrile seizure | 15 | 0.82 | 0 (0-Inf) |
| Ophthalmologic diseases | Glaucoma | 4205 | 0.71 | 0.98 (0.91-1.07) |
| | Cataract | 18010 | 0.30 | 1.03 (0.98-1.08) |
| Dental diseases | Periodontitis | 2773 | 0.25 | 1.07 (0.95-1.2) |
| Other | Amyotrophic lateral sclerosis | 10 | 0.29 | 2.09 (0.53-8.28) |

OR: odds ratio, CI: confidence interval

Results of logistic regression analysis with disease presence as a dependent variable and presence of mosaic events, age, sex and genotyping array as independent variables. Significant level was set at 0.05/1,034 ($p=4.8x10^{-5}$)

Table S6. Fraction of mosaic presence according to age and sex

| Age range | % of males with autosomal event (s.e.) | % of females with autosomal event (s.e.) |
|---|---|---|
| <30 | 4.6% (0.4%) | 4.2% (0.4%) |
| 30-39 | 5.7% (0.4%) | 4.7% (0.3%) |
| 40-49 | 7.6% (0.3%) | 6.5% (0.3%) |
| 50-59 | 11.4% (0.2%) | 8.9% (0.2%) |
| 60-69 | 16.3% (0.2%) | 12.5% (0.2%) |
| 70-79 | 24.7% (0.3%) | 17.6% (0.3%) |
| 80-89 | 31.8% (0.6%) | 24.1% (0.5%) |
| 90+ | 40.7% (2.3%) | 31.5% (1.7%) |

This table contains numeric data plotted in Fig. 2b. s.e.:standard error

Table. S7. Average age and sex of carriers of mosaic event types

| | p-arm LOSS | | q-arm LOSS | | p-arm CNN-LOH | | q-arm CNN-LOH | | GAIN | |
|---|---|---|---|---|---|---|---|---|---|---|
| CHR | AGE | Male ratio | AGE | Male ratio | AGE | Male ratio | AGE | Male ratio | AGE | Male ratio |
| 1 | 72.7 (0.07) | 0.64 (0.04) | 72.9 (0.3) | 0.61 (0.09) | 68.6 (0.01) | 0.6 (0.01) | 69.3 (0.01) | 0.62 (0.01) | 67.5 (0.06) | 0.62 (0.03) |
| 2 | 71.2 (0.04) | 0.52 (0.03) | 69.3 (0.13) | 0.61 (0.06) | 69.2 (0.05) | 0.6 (0.03) | 66.6 (0.04) | 0.55 (0.03) | 70.3 (0.24) | 0.42 (0.07) |
| 3 | 73.6 (0.08) | 0.73 (0.04) | 67.8 (0.22) | 0.55 (0.07) | 67.7 (0.04) | 0.54 (0.03) | 69.7 (0.06) | 0.55 (0.03) | 70.1 (0.11) | 0.68 (0.05) |
| 4 | 66.7 (0.4) | 0.59 (0.08) | 71.3 (0.07) | 0.64 (0.04) | 66.8 (0.16) | 0.64 (0.05) | 71.9 (0.04) | 0.62 (0.03) | 69.8 (0.43) | 0.83 (0.08) |
| 5 | 65.2 (0.86) | 0.67 (0.14) | 72.1 (0.02) | 0.53 (0.02) | 68.2 (0.13) | 0.67 (0.05) | 67.4 (0.05) | 0.61 (0.03) | 68.5 (0.13) | 0.72 (0.05) |
| 6 | 73 (0.48) | 0.64 (0.1) | 69.8 (0.03) | 0.68 (0.02) | 67.3 (0.02) | 0.6 (0.02) | 67.1 (0.04) | 0.64 (0.03) | 68 (0.17) | 0.74 (0.05) |
| 7 | 65 (0.18) | 0.66 (0.06) | 70.7 (0.06) | 0.61 (0.04) | 66.4 (0.08) | 0.57 (0.04) | 66.5 (0.05) | 0.57 (0.03) | 65.8 (0.34) | 0.53 (0.08) |
| 8 | 67.7 (0.28) | 0.63 (0.07) | 70.4 (0.24) | 0.57 (0.07) | 66.3 (0.08) | 0.59 (0.04) | 69.8 (0.05) | 0.61 (0.03) | 74 (0.03) | 0.63 (0.02) |
| 9 | 73.3 (0.51) | 0.63 (0.11) | 71.7 (0.07) | 0.7 (0.04) | 68.2 (0.03) | 0.64 (0.02) | 68.9 (0.01) | 0.59 (0.02) | 71.5 (0.05) | 0.67 (0.03) |
| 10 | 69.8 (0.65) | 0.59 (0.1) | 69 (0.4) | 0.61 (0.09) | 66.1 (0.12) | 0.6 (0.05) | 68.6 (0.05) | 0.61 (0.03) | 63.6 (0.71) | 0.48 (0.11) |
| 11 | 67.3 (0.31) | 0.57 (0.08) | 71.3 (0.02) | 0.63 (0.02) | 65.8 (0.02) | 0.6 (0.02) | 69.9 (0.01) | 0.67 (0.01) | 64 (0.58) | 0.65 (0.09) |
| 12 | 71.8 (0.21) | 0.71 (0.06) | 69 (0.16) | 0.63 (0.06) | 68.5 (0.14) | 0.61 (0.05) | 67.4 (0.04) | 0.62 (0.02) | 71.4 (0.07) | 0.54 (0.05) |
| 13 | | | 71 (0.02) | 0.65 (0.02) | | | 68.2 (0.06) | 0.56 (0.03) | 64.9 (1.35) | 0.77 (0.12) |
| 14 | | | 68.7 (0.04) | 0.66 (0.03) | | | 71.8 (0) | 0.64 (0.01) | 69.7 (0.1) | 0.69 (0.04) |
| 15 | | | 66.7 (0.29) | 0.64 (0.07) | | | 66.9 (0.06) | 0.52 (0.03) | 74.7 (0.01) | 0.77 (0.01) |
| 16 | 67 (0.23) | 0.63 (0.06) | 69.1 (0.48) | 0.57 (0.09) | 68.1 (0.03) | 0.58 (0.02) | 68.5 (0.02) | 0.63 (0.02) | 68.4 (1.56) | 0.67 (0.16) |
| 17 | 72.4 (0.11) | 0.62 (0.05) | 69 (0.14) | 0.46 (0.06) | 70.5 (0.04) | 0.67 (0.03) | 67.4 (0.02) | 0.56 (0.02) | 70.5 (0.11) | 0.58 (0.05) |
| 18 | 71.7 (0.29) | 0.66 (0.08) | 72 (0.4) | 0.53 (0.13) | 68.5 (0.15) | 0.6 (0.06) | 68.1 (0.06) | 0.55 (0.03) | 69.1 (0.06) | 0.59 (0.04) |
| 19 | 60 (4.4) | 1 (0) | 64.7 (2.66) | 0.33 (0.19) | 70.2 (0.05) | 0.59 (0.03) | 67.9 (0.03) | 0.59 (0.03) | 63.2 (0.85) | 0.63 (0.12) |
| 20 | 73.9 (0.6) | 0.63 (0.12) | 72.7 (0.01) | 0.73 (0.01) | 65.4 (0.09) | 0.58 (0.04) | 68 (0.02) | 0.61 (0.02) | 72.8 (1.78) | 0.67 (0.16) |

| | | | | | | | | |
|---|---|---|---|---|---|---|---|---|
| 21 | 65.5 (0.16) | 0.63 (0.06) | | | 67.4 (0.21) | 0.46 (0.07) | 68.5 (0.01) | 0.66 (0.01) |
| 22 | 71.7 (0.13) | 0.77 (0.05) | | | 69.8 (0.06) | 0.65 (0.03) | 70.5 (0.03) | 0.56 (0.03) |

CHR:chromosome, mean (s.e.m) is indicated

Table. S8. Average age and sex of carriers of focal mosaic deletions.

| Focal deletion | Mean AGE (s.e.) | Male ratio (s.e.) |
|---|---|---|
| chr2:20-30Mb | 71.6 (0.06) | 0.52 (0.04) |
| chr3:55-95Mb | 74.7 (0.08) | 0.73 (0.04) |
| chr4:100-110Mb | 73.4 (0.17) | 0.57 (0.06) |
| chr6:125-150Mb | 69.7 (0.09) | 0.69 (0.04) |
| chr12:7-14Mb (*ETV6*) | 73 (0.35) | 0.85 (0.07) |
| chr13:40-60Mb | 71.6 (0.07) | 0.67 (0.04) |
| chr14:20-24Mb | 68.9 (0.04) | 0.65 (0.03) |
| chr16:24-31Mb | 69.7 (0.63) | 0.54 (0.14) |
| chr17:24-31Mb | 68.5 (0.22) | 0.4 (0.07) |
| chr22:24-35Mb (*CHEK2*) | 72.6 (0.18) | 0.81 (0.05) |

s.e.:standard error

Table S9. Associations between mosaic events and hematopoietic traits.

| pheno | Mosaic | Beta | SE | P |
|---|---|---|---|---|
| RBC | chr15_GAIN | -0.20486 | 0.041834 | $9.73 \times 10^{-7}$ |
| Plt | chr20q_LOSS | -0.22059 | 0.045935 | $1.57 \times 10^{-6}$ |
| RBC | chr9_GAIN | -0.40415 | 0.085778 | $2.46 \times 10^{-6}$ |
| Lym | chr14q_LOSS | 0.492607 | 0.104618 | $2.49 \times 10^{-6}$ |
| MCV | chr1_GAIN | 0.418216 | 0.089025 | $2.63 \times 10^{-6}$ |
| RBC | chr8_GAIN | -0.29461 | 0.063182 | $3.12 \times 10^{-6}$ |
| Lym | chr21_GAIN | 0.2582 | 0.056293 | $4.50 \times 10^{-6}$ |
| Neutro | chr14q_LOSS | -0.51383 | 0.113592 | $6.08 \times 10^{-6}$ |
| MCHC | chr9_GAIN | -0.39666 | 0.08978 | $9.96 \times 10^{-6}$ |
| RBC | chr20q_LOSS | -0.16794 | 0.039067 | $1.72 \times 10^{-5}$ |
| Lym | chr22_GAIN | 0.494356 | 0.11672 | $2.28 \times 10^{-5}$ |
| Neutro | chr22_GAIN | -0.53061 | 0.126898 | $2.90 \times 10^{-5}$ |

Lym: lymphocyte count, RBC: red blood cell count, MCV: mean corpuscular volume, MCHC: mean corpuscular hemoglobin concentration, Plt: platelet count, Neutro neutrophil count. Significant associations beyond Bonferroni's correction ($p<0.05/13/88$) are indicated

Table S10. Increased lymphocyte counts associated with presence of mosaic of V(D)J deletion in *TRA*.

| | *TRA* del (+) (N=84) | *TRA* del (-) (N=61992) | P |
|---|---|---|---|
| lymphocyte count | 2078hocy | 1791hocy | 0.0017 |

Mean±an (standard deviation) is indicated

Table S11. Significant correlation between cell fraction of V(D)J deletion in *TRA* and lymphocyte counts.

| Spearman rho (95%CI) | P |
|---|---|
| 0.33 (0.12-0.51) | 0.0037 |

CI: confidence interval

Table S12. **Distribution of mCAs by chromosome and copy number in BioBank Japan and UK Biobank.**

| Chromsome | Loss freq, BBJ | Loss freq, UKB | CNN-LOH freq, BBJ | CNN-LOH freq, UKB | Gain freq, BBJ | Gain freq, UKB |
|---|---|---|---|---|---|---|
| 1 | 0.8% | 0.7% | 7.6% | 8.2% | 1.1% | 0.5% |
| 2 | 1.5% | 1.5% | 2.0% | 1.8% | 0.3% | 0.2% |
| 3 | 0.9% | 0.5% | 1.4% | 1.6% | 0.5% | 1.2% |
| 4 | 1.1% | 1.0% | 1.3% | 1.7% | 0.1% | 0.2% |
| 5 | 2.3% | 1.2% | 0.5% | 0.9% | 0.4% | 0.6% |
| 6 | 2.1% | 0.8% | 2.5% | 2.2% | 0.3% | 0.2% |
| 7 | 1.3% | 1.2% | 0.8% | 1.2% | 0.2% | 0.2% |
| 8 | 0.5% | 0.5% | 0.5% | 0.9% | 1.9% | 1.1% |
| 9 | 0.9% | 0.4% | 3.5% | 4.9% | 1.0% | 0.8% |
| 10 | 0.3% | 2.0% | 0.8% | 0.9% | 0.1% | 0.1% |
| 11 | 2.4% | 2.1% | 4.6% | 6.1% | 0.1% | 0.0% |
| 12 | 0.6% | 0.6% | 1.3% | 1.8% | 0.6% | 3.7% |
| 13 | 2.3% | 4.2% | 1.1% | 3.1% | 0.1% | 0.1% |
| 14 | 1.7% | 1.1% | 13.6% | 4.9% | 0.5% | 1.1% |
| 15 | 0.2% | 0.3% | 1.1% | 2.8% | 4.5% | 1.6% |
| 16 | 0.5% | 1.3% | 2.8% | 3.3% | 0.0% | 0.1% |
| 17 | 0.9% | 1.6% | 2.9% | 3.0% | 0.5% | 0.9% |
| 18 | 0.3% | 0.4% | 0.5% | 0.7% | 1.0% | 1.4% |
| 19 | 0.1% | 0.1% | 1.1% | 2.4% | 0.1% | 0.3% |
| 20 | 5.2% | 3.2% | 1.7% | 1.4% | 0.0% | 0.1% |
| 21 | 0.4% | 0.4% | 0.3% | 1.0% | 5.6% | 1.1% |

| | | | | | | |
|---|---|---|---|---|---|---|
| 22 | 0.4% | 1.0% | 1.1% | 2.3% | 1.5% | 1.3% |

This table provides numeric data plotted in Fig. 3a,b. Frequencies indicate the contribution of each event type to the total number of mCAs classified as loss, CNN-LOH, or gain in each data set. Data for UK Biobank events are from parallel work on the UK Biobank cohort[17].

Table. S13. Concordance of positional coverage of mosaic events between Japanese and UK population

| Chr | All mosaic | Loss | CNN-LOH | Gain |
|---|---|---|---|---|
| 1 | 0.97 (0.97-0.97) | 0.41 (0.38-0.45) | 0.98 (0.97-0.98) | 0.91 (0.9-0.92) |
| 2 | 0.99 (0.99-0.99) | 0.96 (0.96-0.96) | 0.97 (0.97-0.97) | 0.23 (0.19-0.27) |
| 3 | 0.8 (0.79-0.82) | 0.92 (0.91-0.93) | 0.97 (0.97-0.97) | 0.98 (0.98-0.98) |
| 4 | 0.99 (0.99-0.99) | 0.72 (0.69-0.74) | 0.99 (0.99-0.99) | 0.8 (0.78-0.82) |
| 5 | 0.85 (0.84-0.86) | 0.99 (0.99-0.99) | 0.96 (0.95-0.96) | 0.99 (0.99-1) |
| 6 | 0.91 (0.91-0.92) | 0.86 (0.84-0.87) | 0.99 (0.99-0.99) | 0.89 (0.88-0.9) |
| 7 | 0.98 (0.98-0.98) | 0.9 (0.9-0.91) | 0.97 (0.96-0.97) | 0.83 (0.81-0.84) |
| 8 | 0.88 (0.87-0.89) | 0.63 (0.6-0.66) | 0.96 (0.95-0.96) | 0.95 (0.94-0.95) |
| 9 | 0.84 (0.82-0.85) | 0.97 (0.96-0.97) | 0.91 (0.9-0.92) | 0.95 (0.94-0.96) |
| 10 | 0.79 (0.77-0.81) | -0.03 (-0.08-0.03) | 0.94 (0.93-0.94) | 0.09 (0.04-0.15) |
| 11 | 0.87 (0.86-0.89) | 0.82 (0.8-0.84) | 0.8 (0.78-0.82) | -0.21 (-0.26--0.16) |
| 12 | 0.99 (0.99-0.99) | 0.79 (0.77-0.81) | 0.98 (0.98-0.98) | 0.9 (0.88-0.91) |
| 13 | 0.87 (0.86-0.89) | 0.91 (0.89-0.92) | 0.98 (0.98-0.98) | 0.86 (0.84-0.87) |
| 14 | 0.99 (0.99-0.99) | 0.13 (0.06-0.19) | 0.99 (0.99-0.99) | 0.73 (0.7-0.76) |
| 15 | 0.96 (0.95-0.96) | -0.04 (-0.11-0.03) | 0.98 (0.98-0.98) | 0.99 (0.99-0.99) |
| 16 | 0.92 (0.91-0.93) | 0.56 (0.51-0.6) | 0.93 (0.92-0.94) | 0.02 (-0.04-0.09) |
| 17 | 0.99 (0.99-0.99) | 0.99 (0.99-0.99) | 1 (1-1) | 1 (1-1) |
| 18 | 0.94 (0.93-0.94) | 0.99 (0.98-0.99) | 0.98 (0.98-0.98) | 0.77 (0.74-0.8) |
| 19 | 0.91 (0.89-0.92) | 0.31 (0.24-0.38) | 0.9 (0.89-0.92) | -0.42 (-0.48--0.35) |
| 20 | 0.99 (0.99-0.99) | 0.99 (0.99-0.99) | 0.97 (0.96-0.97) | 0.7 (0.65-0.73) |
| 21 | 0.62 (0.55-0.69) | 0.16 (0.05-0.26) | 0.94 (0.92-0.95) | 0.76 (0.71-0.8) |
| 22 | 0.93 (0.92-0.94) | 0.61 (0.54-0.68) | 0.97 (0.96-0.98) | 0.85 (0.82-0.88) |
| Mean | 0.91±0.09 | 0.66±0.35 | 0.96±0.04 | 0.66±0.42 |

Pearson's correlation coefficients between BBJ and UKB are indicated.

Mean indicates mean correlation across chromosomes (not concatenating all chromosomes).

Chr: chromosome

Table. S14. Genes most frequently involved in focal deletions in BBJ.

| Chr | Top gene |
|---|---|
| 1 | PTPN22, LOC100287722, BCL2L15, AP4B1, DCLRE1B, HIPK1, OLFML3, RPL13AP10, SYT6, MRP63P1, LOC100421116 |
| 2 | DNMT3A |
| 3 | FOXP1 |
| 4 | TET2 |
| 5 | FBXL17 |
| 6 | LOC100130476, TNFAIP3 |
| 7 | LOC136157, RPS2P31, GPR37, LOC154872, POT1 |
| 8 | FAM87A, FBXO25, C8orf42, DLGAP2, LOC100130321, LOC100507448, CLN8, MIR3674, MIR596, ARHGEF10, LOC100131395, KBTBD11, MYOM2 |
| 9 | LOC286370, MIR4290, IL6RP1, OR7E31P, OR7E116P, LOC340515, DIRAS2, OR7E109P, OR7E108P, SYK |
| 10 | PTEN, RPL11P3, LOC100128990, VN1R55P, RNLS, LIPJ, RPL7P34 |
| 11 | DDX10, CYCSP29 |
| 12 | LOH12CR1, DUSP16 |
| 13 | DLEU7 |
| 14 | TRAV24 |
| 15 | FMN1, LOC100421433, LOC100652815, LOC100652857, RYR3 |
| 16 | RPL10AP12, IRF8 |
| 17 | PFN1, ENO3, SPAG7, CAMTA2, INCA1, KIF1C |
| 18 | DLGAP1 |
| 20 | PTPRT |
| 21 | COL6A2, FTCD |
| 22 | CHEK2, CCDC117, XBP1, ZNRF3 |

Since chr19 has fewer than 20 loss events, we do not show results in chr 19.

Table S15. Genes frequently involved with focal deletions in Japanese and not in UK population.

| Chr | Genes |
|---|---|
| 1 | LOC100533666,ST13P20,LPHN2,CDK4PS |
| 3 | LOC100421672,FHIT,LOC100421670 |
| 4 | RPL21P46,SCFD2,FIP1L1,LNX1,LOC100129728,RPL21P44,CHIC2,RPL22P13,PDGFRA,LOC100421808,MIR548AG1,LOC100421630,VEGFC,NEIL3,AGA |
| 6 | PEX7,SLC35D3,RPL35AP3,NHEG1,IL20RA,IL22RA2,IFNGR1,OLIG3,LOC391040,LOC442263,LOC100507406,LOC100507429,LOC100130476,TNFAIP3,RPSAP42,PERP,KIAA1244,PBOV1,HEBP2,NHSL1,MIR3145 |
| 7 | NXPH1,RPL9P19,LOC100287551,NDUFA4,DGKB,EEF1A1P26,LOC100533714,VWC2,MAGI2,MAGI2-AS3,RPL10P11,GNAI1,LOC100420647,IMMP2L,LRRN3 |
| 8 | RPL23AP53,ZNF596,FAM87A,FBXO25,C8orf42,CSMD3,LOC100289099,MIR2053,EXT1 |
| 9 | LOC100128505 |
| 10 | CTNNA3,LOC100533794 |
| 11 | LOC729790 |
| 14 | LOC100288613,TRA@,TRAV1-1,OR10G2,TRAV1-2,ARL6IP1P1,OR4E2,OR4E1,TRAV2,TRAV3,TRAV4,TRAV5,RPL4P1,TRAV6,TRAV7,TRAV8-1,TRAV9-1,TRAV10,TRAV11,TRAV12-1,TRAV8-2,TRAV8-3,TRAV13-1,TRAV12-2,TRAV8-4,TRAV8-5,TRAV13-2,TRAV14DV4,TRAV9-2,TRAV15,TRAV12-3,TRAV8-6,TRAV16,TRAV17,TRAV18,TRAV19,TRAV20,TRAV21,TRAV22,TRAV23DV6,TRDV1,TRAV24,TRAV25,TRAV26-1,TRAV8-7,TRAV27,TRAV28,TRAV29DV5,TRAV30,TRAV31,TRAV32,TRAV33,TRAV26-2,TRAV34,TRAV35,TRAV36DV7,TRAV37,TRAV38-1,TRAV38-2DV8,TRAV39,TRAV40,TRAV41,TRD@,TRDV2,TRDD1,TRDD2,TRDD3,TRDJ1,TRDJ4,TRDJ2,TRDJ3 |
| 15 | KIAA1370,LINC00052,NTRK3 |
| 16 | LOC100131080 |
| 18 | LOC100422496 |
| 22 | LARGE,MIR4764,LOC100506195 |

Table S16. No associations between chr5q CNN-LOH and variants in *RAD50*.

| Chr:Pos | Ref | Alt | Case freq | Cont freq | P | OR (95%CI) |
|---|---|---|---|---|---|---|
| 5:131954134 | A | G | 0.006466 | 0.000782 | 0.006164 | 8.3 (2.7-26) |
| 5:131977046 | C | T | 0.06681 | 0.04117 | 0.009516 | 1.7 (1.2-2.4) |
| 5:131874403 | T | G | 0.01078 | 0.002778 | 0.01029 | 3.9 (1.6-9.5) |
| 5:131916654 | G | A | 0.006466 | 0.001083 | 0.01471 | 6 (1.9-18.8) |
| 5:131991821 | C | A | 0.01078 | 0.003476 | 0.02432 | 3.1 (1.3-7.6) |
| 5:131875296 | G | C | 0.01078 | 0.003496 | 0.02485 | 3.1 (1.3-7.5) |
| 5:131940799 | T | C | 0.002155 | 5.50E-05 | 0.0265 | 39.2 (5.2-293.8) |
| 5:131891706 | G | A | 0.006466 | 0.001422 | 0.02959 | 4.6 (1.5-14.3) |
| 5:131962216 | A | G | 0.002155 | 6.66E-05 | 0.03172 | 32.4 (4.4-240.6) |
| 5:131893543 | T | C | 0.01724 | 0.007925 | 0.03374 | 2.2 (1.1-4.4) |
| 5:131926388 | G | A | 0.006466 | 0.001605 | 0.03994 | 4 (1.3-12.6) |
| 5:131993276 | T | TCTGAA | 0.002155 | 9.56E-05 | 0.04464 | 22.6 (3.1-165.5) |

Variants showing p-values less than 0.05 in the region of *RAD50* are indicated.

Chr: chromosome, Pos: position, Freq: frequency, OR: odds ratio, CI: confidence interval

Table. S17. Associations of *MAD1L1* variant rs12699483 with gain events.

| Chr | Events N | P | OR (95%CI) |
|---|---|---|---|
| chr1 | 5 | 0.34 | 2.05 (0.58-7.25) |
| chr2 | 6 | 0.38 | 1.91 (0.61-6.03) |
| **chr3** | **59** | **0.025** | **1.51 (1.05-2.17)** |
| chr4 | 18 | 0.027 | 2.15 (1.1-4.2) |
| chr5 | 3 | 0.7 | 1.37 (0.28-6.77) |
| chr6 | 9 | 0.34 | 1.71 (0.67-4.33) |
| chr7 | 9 | 0.63 | 1.37 (0.54-3.44) |
| **chr8** | **320** | **0.15** | **1.13 (0.96-1.31)** |
| **chr9** | **129** | **0.05** | **1.29 (1.01-1.64)** |
| chr10 | 4 | 1 | 0.82 (0.2-3.43) |
| chr11 | 5 | 0.11 | 3.19 (0.82-12.33) |
| **chr12** | **105** | **0.4** | **1.13 (0.86-1.48)** |
| chr13 | 9 | 0.24 | 0.53 (0.19-1.47) |
| **chr14** | **96** | **0.12** | **1.26 (0.95-1.67)** |
| **chr15** | **855** | **$4.3x10^{-22}$** | **1.6 (1.46-1.76)** |
| chr16 | 2 | 1 | 1.37 (0.19-9.7) |
| chr17 | 71 | 0.87 | 1.03 (0.74-1.43) |
| **chr18** | **182** | **0.96** | **0.99 (0.8-1.22)** |
| chr19 | 8 | 0.13 | 2.28 (0.83-6.27) |
| chr20 | 0 | 1 | NA |
| **chr21** | **1060** | **0.085** | **1.08 (0.99-1.18)** |
| **chr22** | **264** | **0.17** | **1.13 (0.95-1.34)** |

Gain events covering >50% of the genotyped span of the chromosome were tested for association.

OR: odds ratio, CI: confidence interval

Table S18. Allele frequencies of BBJ mosaicism risk variants in European population

| Mosaic | Gene | Chr | Pos | Variant | Ref | Var | Jpn AF | Eur AF |
|---|---|---|---|---|---|---|---|---|
| Cis | | | | | | | | |
| chr1p_CNN | MPL | 1 | 45444734 | 1: 45444734 | G | A | 0.00016 | 0.00033* |
| chr8q_CNN | NBN | 8 | 90949282 | rs756831345 | C | A | 0.00061 | 0 |
| chr9p_CNN | JAK2 | 9 | 5026293 | rs2183137 | A | G | 0.24 | 0.29 |
| chr11q_CNN | MRE11 | 11 | 94160189 | 11:94160189 | G | A | 0.00011 | 0 |
| chr14q_CNN | NEDD8/ TINF2 | 14 | 24711798 | rs28372734 | C | G | 0.073 | 0.0020 |
| chr14q_CNN | TCL1A | 14 | 96180242 | rs1122138 | C | A | 0.05 | 0.15 |
| chr14q_CNN | DLK1 | 14 | 101175967 | rs10873520 | G | A | 0.30 | 0.20 |
| chr16q_CNN | CTU2 | 16 | 88781475 | rs200779411 | C | T | 0.00065 | 0.00014 |
| trans | | | | | | | | |
| chr14q_CNN | TERT | 5 | 1287194 | rs2853677 | A | G | 0.31 | 0.41 |
| Chr15_gain | MAD1L1 | 7 | 1975624 | rs12699483 | C | G | 0.42 | 0.35 |

Chr:chromosome, Pos: chromosomal base pair position, Ref: reference allele, Var: variant (tested) allele, Jpn AF: Japanese allele frequency calculated by control subjects, Eur MAF: European (non-Finnish) allele frequency obtained from gnomAD;
*present in gnomAD European data, but absent in UKBB.

Table S19. Associations between mosaic events and mortality.

| | HR (95%CI) | P |
|---|---|---|
| Overall mortality | 1.10 (1.05-1.16) | $2.7x10^{-5}$ |
| All cancer mortality | 1.13 (1.03-1.25) | 0.014 |
| Blood cancer mortality | 2.85 (2.15-3.78) | $4.1x10^{-13}$ |
| Leukemia mortality | 4.70 (3.26-6.78) | $1.0x10^{-16}$ |
| Malignant lymphoma mortality | 1.39 (0.78-2.47) | 0.26 |
| Multiple myeloma mortality | 0.87 (0.26-2.88) | 0.82 |
| Other cancer mortality | 1.04 (0.94-1.16) | 0.46 |
| Cardiovascular mortality | 1.07 (0.94-1.22) | 0.32 |
| Coronary heart disease mortality | 1.10 (0.93-1.30) | 0.27 |
| Ischemic stroke mortality | 1.03 (0.83-1.27) | 0.79 |

CI:confidence interval, HR: hazard ratio

Table S20. Associations between mosaic events and leukemia mortality

| | LOSS_CNN_GAIN | | LOSS | | CNN-LOH | | GAIN | |
|---|---|---|---|---|---|---|---|---|
| | P | OR (95%CI) | P | OR (95%CI) | P | OR (95%CI) | P | OR (95%CI) |
| chr1 | 0.00020 | 7.8 (2.8-18.1) | 1.0 | 0 (0-36.6) | 0.098 | 3.9 (0.5-14.8) | $2.6x10^{-5}$ | 27.4 (6.8-79.8) |
| chr2 | 1.0 | 0 (0-9) | 1.0 | 0 (0-16.4) | 1.0 | 0 (0-24.4) | 1.0 | 0 (0-129.4) |
| chr3 | 0.032 | 7.5 (0.9-29.1) | 1.0 | 0 (0-31.2) | 0.0061 | 18.6 (2.1-75) | 1.0 | 0 (0-154.8) |
| chr4 | 0.040 | 6.6 (0.8-25.3) | 0.011 | 13.7 (1.6-54.3) | 1.0 | 0 (0-26.5) | 1.0 | 0 (0-571.1) |
| chr5 | 0.080 | 4.4 (0.5-16.8) | 0.048 | 6 (0.7-23) | 1.0 | 0 (0-71.6) | 1.0 | 0 (0-68.5) |
| chr6 | 0.0046 | 6.3 (1.7-17.1) | 0.00056 | 11.5 (3-31.7) | 1.0 | 0 (0-15.1) | 1.0 | 0 (0-146) |
| chr7 | $1.8x10^{-5}$ | 17.2 (5.3-43.3) | $7.3x10^{-5}$ | 20 (5.2-56.1) | 1.0 | 0 (0-51.5) | 0.020 | 56.5 (1.3-422.4) |
| chr8 | 0.0058 | 8.8 (1.7-27.8) | 0.062 | 16.8 (0.4-111.4) | 0.062 | 16.9 (0.4-111.2) | 0.21 | 4.4 (0.1-26.6) |
| chr9 | 0.024 | 5.1 (1-15.9) | 1.0 | 0 (0-24.3) | 1.0 | 0 (0-12) | 0.00024 | 28.8 (5.4-97.6) |
| chr10 | 0.13 | 7.7 (0.2-47.1) | 0.066 | 15.8 (0.4-106.7) | 1.0 | 0 (0-95.4) | 1.0 | 0 (0-238.5) |
| chr11 | 0.00013 | 8.5 (3-19.7) | $5.8x10^{-5}$ | 13.5 (4.2-34) | 0.29 | 3 (0.1-17.8) | 1.0 | 0 (0-426.7) |
| chr12 | $1.3x10^{-5}$ | 18.8 (5.8-47.8) | 0.082 | 12.3 (0.3-78.2) | 0.00028 | 26.6 (5.1-87.1) | 0.076 | 13.7 (0.3-90.5) |
| chr13 | 0.0017 | 8.5 (2.2-23.4) | 0.00060 | 11.4 (3-31.6) | 1.0 | 0 (0-35) | | |
| chr14 | $1.3x10^{-5}$ | 7.1 (3.1-14.4) | 0.24 | 3.8 (0.1-22.5) | $6.8x10^{-5}$ | 7.8 (3-17.2) | 0.087 | 11.6 (0.3-73.9) |
| chr15 | 1.0 | 1 (0-5.9) | 1.0 | 0 (0-104.9) | 1.0 | 0 (0-31.1) | 0.57 | 1.2 (0-7.1) |
| chr16 | 0.0094 | 7.3 (1.5-22.9) | 0.096 | 10.5 (0.2-66.5) | 0.040 | 6.6 (0.8-25.3) | | |
| chr17 | $6.6x10^{-7}$ | 16.1 (6.2-35.8) | 0.0079 | 16.1 (1.9-64.2) | $1.8x10^{-5}$ | 17.5 (5.4-44.3) | 1.0 | 0 (0-60.9) |
| chr18 | 0.019 | 10 (1.2-39.2) | 0.050 | 21.6 (0.5-149.9) | 1.0 | 0 (0-80.9) | 0.12 | 8.5 (0.2-52.3) |
| chr19 | 0.18 | 5.3 (0.1-31.6) | 1.0 | 0 (0-599.3) | 1.0 | 0 (0-22.6) | | |
| chr20 | 0.00098 | 5.8 (2-13.4) | 0.0028 | 5.6 (1.7-14.1) | 0.13 | 7.2 (0.2-43.5) | | |

| | | | | | | | | |
|---|---|---|---|---|---|---|---|---|
| chr21 | 0.017 | 4.3 (1.1-11.5) | 0.063 | 16.3 (0.4-103.6) | 1.0 | 0 (0-123) | 0.058 | 3.5 (0.7-10.9) |
| chr22 | 1.0 | 0 (0-7.2) | 1.0 | 0 (0-75.6) | 1.0 | 0 (0-22) | 1.0 | 0 (0-13.1) |
| chr1p | | | | | 0.026 | 8.4 (1-32.4) | | |
| chr1q | | | | | 1.0 | 0 (0-14.3) | | |
| chr2p | | | | | 1.0 | 0 (0-55.6) | | |
| chr2q | | | | | 1.0 | 0 (0-51.9) | | |
| chr3p | | | | | 1.0 | 0 (0-72.5) | | |
| chr3q | | | | | 0.0015 | 40.6 (4.4-176.3) | | |
| chr4q | | | | | 1.0 | 0 (0-36.7) | | |
| chr5q | | | | | 1.0 | 0 (0-77.4) | | |
| chr6p | | | | | 1.0 | 0 (0-18.7) | | |
| chr6q | | | | | 1.0 | 0 (0-89.6) | | |
| chr7p | | | | | 1.0 | 0 (0-145.6) | | |
| chr7q | | | | | 1.0 | 0 (0-108.7) | | |
| chr8p | | | | | 1.0 | 0 (0-821.4) | | |
| chr8q | | | | | 0.054 | 19.8 (0.5-134.7) | | |
| chr9p | | | | | 1.0 | 0 (0-24.6) | | |
| chr9q | | | | | 1.0 | 0 (0-24.5) | | |
| chr10p | | | | | 1.0 | 0 (0-318.3) | | |
| chr10q | | | | | 1.0 | 0 (0-177.1) | | |
| chr11p | | | | | 0.12 | 8.4 (0.2-50.8) | | |
| chr11q | | | | | 1.0 | 0 (0-18.6) | | |
| chr12p | | | | | 1.0 | 0 (0-264.9) | | |

| | | |
|---|---|---|
| chr12q | 0.00015 | 33.1 (6.3-111) |
| chr16p | 1.0 | 0 (0-41.2) |
| chr16q | 0.019 | 10 (1.2-39.2) |
| chr17p | $4.5x10^{-7}$ | 82 (19.5-259) |
| chr17q | 0.21 | 4.2 (0.1-25.2) |
| chr18p | 1.0 | 0 (0-692.8) |
| chr18q | 1.0 | 0 (0-109.5) |
| chr19p | 1.0 | 0 (0-40.2) |
| chr19q | 1.0 | 0 (0-57.3) |
| chr20p | 1.0 | 0 (0-507.1) |
| chr20q | 0.12 | 8.1 (0.2-49.3) |

CI:confidence interval, OR:odds ratio

Table S21. Associations between leukemia mortality and cell fraction of mosaic events.

| cell fraction | case subjects | coeff | SE | HR (95%CI) | P |
|---|---|---|---|---|---|
| 1-3% | ALL | 0.97 | 0.28 | 2.64 (1.51-4.6) | $6.2\times10^{-4}$ |
| 3-5% | ALL | 1.43 | 0.43 | 4.19 (1.82-9.64) | $7.5\times10^{-4}$ |
| 5%- | ALL | 2.07 | 0.23 | 7.95 (5.06-12.48) | $<2.2\times10^{-16}$ |

Subjects having hematopoietic malignancy are excluded. Results in Cox proportional hazard model with age, age^2, sex, smoking ,disease status and genotyping arrays in covariates. Coeff: coefficient, SE: standard error, CI:confidence interval, HR: hazard ratio

Table S22. Associations between leukemia mortality and presence of multiple mosaic events.

| cell fraction | case subjects | coeff | SE | HR (95%CI) | P |
|---|---|---|---|---|---|
| 1-3% | multiple | - | - | - | - |
| 1-3% | single | 0.95 | 0.29 | 2.58 (1.45-4.57) | 0.0012 |
| 3-5% | multiple | 2.69 | 0.59 | 14.74 (4.59-47.32) | $6.1\times10^{-6}$ |
| 3-5% | single | 0.87 | 0.59 | 2.4 (0.76-7.62) | 0.14 |
| 5%- | multiple | 2.71 | 0.35 | 15.02 (7.61-29.63) | $5.6\times10^{-15}$ |
| 5%- | single | 1.67 | 0.29 | 5.33 (3-9.47) | $1.1\times10^{-8}$ |

Subjects having hematopoietic malignancy are excluded. Results in Cox proportional hazard model with age, age^2, sex, smoking ,disease status and genotyping arrays in covariates. Coeff: coefficient, SE: standard error, CI:confidence interval, HR: hazard ratio

Table S23. Associations between mosaic events and overall mortality.

| | LOSS_CNN_GAIN | | LOSS | | CNN | | GAIN | |
|---|---|---|---|---|---|---|---|---|
| | P | OR (95%CI) | P | OR (95%CI) | P | OR (95%CI) | P | OR (95%CI) |
| chr1 | 0.0078 | 1.3 (1.1-1.6) | 0.035 | 1.7 (1-2.9) | 0.060 | 1.3 (1-1.6) | 0.41 | 1.2 (0.7-1.9) |
| chr2 | 0.24 | 1.2 (0.9-1.6) | 0.62 | 1.1 (0.7-1.7) | 0.72 | 1.1 (0.7-1.7) | 0.17 | 1.8 (0.7-4.5) |
| chr3 | 0.023 | 1.4 (1-2) | 0.24 | 1.4 (0.8-2.3) | 0.37 | 1.3 (0.7-2.1) | 0.040 | 2.3 (1-5) |
| chr4 | 0.16 | 1.3 (0.9-1.7) | 0.090 | 1.5 (0.9-2.3) | 0.90 | 1 (0.6-1.7) | 1.0 | 1.2 (0.1-10) |
| chr5 | 0.016 | 1.4 (1.1-1.8) | 0.018 | 1.4 (1.1-1.9) | 0.85 | 1.1 (0.5-2.3) | 0.40 | 1.3 (0.7-2.6) |
| chr6 | 0.72 | 1 (0.7-1.2) | 0.87 | 1 (0.7-1.3) | 0.51 | 0.9 (0.6-1.3) | 0.34 | 1.6 (0.6-4) |
| chr7 | 0.80 | 1.1 (0.7-1.5) | 0.92 | 1 (0.7-1.6) | 0.87 | 0.9 (0.4-1.8) | 0.40 | 1.6 (0.4-4.8) |
| chr8 | 0.042 | 1.4 (1-1.8) | 0.69 | 0.8 (0.3-1.8) | 0.47 | 1.3 (0.6-2.7) | 0.018 | 1.5 (1.1-2.2) |
| chr9 | 0.00078 | 1.5 (1.2-1.9) | 0.71 | 0.9 (0.5-1.5) | 0.0016 | 1.6 (1.2-2.2) | 0.0094 | 2 (1.2-3.4) |
| chr10 | 0.89 | 0.9 (0.5-1.6) | 0.53 | 0.7 (0.2-1.8) | 0.43 | 1.4 (0.6-3.1) | 0.70 | 0.4 (0-2.8) |
| chr11 | 0.10 | 1.2 (1-1.5) | 0.40 | 1.1 (0.8-1.5) | 0.21 | 1.2 (0.9-1.7) | 0.30 | 2.1 (0.5-8.1) |
| chr12 | 0.096 | 1.3 (0.9-1.9) | 0.23 | 1.5 (0.8-2.6) | 0.60 | 1.2 (0.7-1.9) | 0.26 | 1.5 (0.7-3.2) |
| chr13 | 0.25 | 1.2 (0.9-1.5) | 0.28 | 1.2 (0.9-1.6) | 0.88 | 1.1 (0.6-1.9) | | |
| chr14 | 0.00017 | 1.3 (1.1-1.6) | 0.18 | 1.3 (0.9-1.8) | 0.00082 | 1.4 (1.1-1.6) | 0.29 | 1.4 (0.7-2.5) |
| chr15 | 0.66 | 1 (0.9-1.3) | 0.18 | 0.4 (0.1-1.5) | 0.18 | 0.7 (0.3-1.2) | 0.20 | 1.1 (0.9-1.4) |
| chr16 | 0.19 | 0.8 (0.6-1.1) | 0.28 | 0.6 (0.3-1.4) | 0.54 | 0.9 (0.6-1.3) | | |
| chr17 | 0.054 | 1.3 (1-1.7) | 0.22 | 1.4 (0.8-2.2) | 0.25 | 1.2 (0.9-1.7) | 0.31 | 1.4 (0.7-2.8) |
| chr18 | 0.62 | 1.1 (0.7-1.6) | 0.22 | 0.4 (0.1-1.4) | 0.22 | 1.5 (0.7-2.9) | 1.0 | 1 (0.6-1.8) |
| chr19 | 1.0 | 1 (0.6-1.6) | 0.65 | 0.4 (0-4.6) | 0.81 | 1.1 (0.7-1.7) | | |
| chr20 | 0.0067 | 1.3 (1.1-1.5) | 0.016 | 1.3 (1-1.5) | 0.17 | 1.4 (0.8-2.2) | | |

| | | | | | | | | |
|---|---|---|---|---|---|---|---|---|
| chr21 | 0.52 | 0.9 (0.8-1.1) | 0.32 | 0.6 (0.2-1.4) | 0.81 | 1.1 (0.3-3) | 0.67 | 1 (0.8-1.2) |
| chr22 | 0.25 | 0.8 (0.6-1.1) | 0.29 | 1.5 (0.7-3) | 0.55 | 0.8 (0.5-1.4) | 0.10 | 0.7 (0.5-1.1) |
| chr1p | | | | | 0.12 | 1.3 (0.9-1.8) | | |
| chr1q | | | | | 0.27 | 1.2 (0.9-1.7) | | |
| chr2p | | | | | 0.72 | 1.1 (0.5-2.4) | | |
| chr2q | | | | | 1.0 | 1 (0.5-1.9) | | |
| chr3p | | | | | 0.71 | 1.2 (0.5-2.4) | | |
| chr3q | | | | | 0.58 | 1.3 (0.6-2.7) | | |
| chr4q | | | | | 0.78 | 1.1 (0.6-1.9) | | |
| chr5q | | | | | 0.71 | 1.2 (0.5-2.5) | | |
| chr6p | | | | | 0.18 | 0.7 (0.4-1.1) | | |
| chr6q | | | | | 0.36 | 1.4 (0.7-2.9) | | |
| chr7p | | | | | 1.0 | 0.8 (0.2-2.8) | | |
| chr7q | | | | | 0.83 | 1.1 (0.4-2.6) | | |
| chr8p | | | | | 0.66 | 1.3 (0.1-9.7) | | |
| chr8q | | | | | 0.54 | 1.3 (0.6-3.1) | | |
| chr9p | | | | | 0.061 | 1.5 (1-2.4) | | |
| chr9q | | | | | 0.024 | 1.6 (1-2.4) | | |
| chr10p | | | | | 0.72 | 1.2 (0.2-5.9) | | |
| chr10q | | | | | 0.35 | 1.5 (0.6-3.9) | | |
| chr11p | | | | | 0.61 | 1.1 (0.7-1.9) | | |
| chr11q | | | | | 0.31 | 1.2 (0.8-1.8) | | |
| chr12p | | | | | 0.77 | 1.3 (0.3-4.3) | | |

| | | |
|---|---|---|
| chr12q | 0.77 | 1.1 (0.6-2) |
| chr16p | 0.57 | 0.8 (0.4-1.5) |
| chr16q | 0.91 | 1 (0.6-1.5) |
| chr17p | 0.0028 | 2.3 (1.3-4.2) |
| chr17q | 0.56 | 0.9 (0.6-1.3) |
| chr18p | 0.037 | 3.6 (0.9-14.4) |
| chr18q | 0.68 | 1.2 (0.5-2.7) |
| chr19p | 0.62 | 1.2 (0.6-2.3) |
| chr19q | 1.0 | 1 (0.5-1.9) |
| chr20p | 0.70 | 1.4 (0.2-6.4) |
| chr20q | 0.49 | 1.2 (0.7-2) |

CI:confidence interval, OR:odds ratio

Table. S24. Oligonucleotide sequences used for luciferase assay.

| oligonucleotide | sequence |
|---|---|
| MRE11_C allele | TTCGTAGAATGAGTTAGGGAGGAGC**C**TCCCTTGATTTTTTTGGAATAATTT |
| MRE11_C allele_c | AAATTATTCCAAAAAAATCAAGGGA**G**GCTCCTCCCTAACTCATTCTACGAA |
| MRE11_T allele | TTCGTAGAATGAGTTAGGGAGGAGC**T**TCCCTTGATTTTTTTGGAATAATTT |
| MRE11_T allele_c | AAATTATTCCAAAAAAATCAAGGGA**A**GCTCCTCCCTAACTCATTCTACGAA |
| MPL_G allele | GGAAGTATTTACACCACAAAAAGCA**G**CAAATGCTACCAAACAGAACACCCT |
| MPL_G allele_c | AGGGTGTTCTGTTTGGTAGCATTTG**C**TGCTTTTTGTGGTGTAAATACTTCC |
| MPL_A allele | GGAAGTATTTACACCACAAAAAGCA**A**CAAATGCTACCAAACAGAACACCCT |
| MPL_A allele_c | AGGGTGTTCTGTTTGGTAGCATTTG**T**TGCTTTTTGTGGTGTAAATACTTCC |

_c indicates complementary sequences. The position of variants are shown in bold.

Table. S25. Combinations of significantly co-occurring mosaic events in different chromosomes

| mosaic1 | mosaic2 | P | OR (95%CI) | Shared by UK |
|---|---|---|---|---|
| chr3_GAIN | chr18_GAIN | $3.82x10^{-25}$ | 198 (98-365) | 1 |
| chr1_GAIN | chr7q_LOSS | $1.32x10^{-16}$ | 66 (32-123) | 0 |
| chr20q_LOSS | chr14q_CNN-LOH | $1.1x10^{-13}$ | 4 (3-5) | 0 |
| chr14q_LOSS | chr21_GAIN | $2.02x10^{-13}$ | 11 (6-17) | 0 |
| chr1_GAIN | chr9_GAIN | $8.31x10^{-11}$ | 40 (17-81) | 0 |
| chr12_GAIN | chr13q_LOSS | $7.0x10^{-10}$ | 30 (13-62) | 1 |
| chr1p_CNN-LOH | chr14q_CNN-LOH | $2.11x10^{-9}$ | 3 (2-4) | 0 |
| chr3_GAIN | chr12_GAIN | $2.26x10^{-9}$ | 109 (34-274) | 1 |
| chr6p_CNN-LOH | chr16p_CNN-LOH | $2.95x10^{-9}$ | 10 (5-17) | 0 |
| chr14q_LOSS | chr22_GAIN | $4.18x10^{-9}$ | 18 (8-35) | 0 |
| chr3_GAIN | chr9_GAIN | $3.12x10^{-8}$ | 63 (20-157) | 0 |
| chr18_GAIN | chr22_GAIN | $3.86x10^{-8}$ | 24 (9-50) | 0 |
| chr17_GAIN | chr21q_LOSS | $5.86x10^{-8}$ | 123 (32-338) | 0 |
| chr12q_LOSS | chr14q_LOSS | $1.49x10^{-7}$ | 46 (14-115) | 0 |
| chr12_GAIN | chr18_GAIN | $1.85x10^{-7}$ | 44 (14-107) | 1 |
| chr3p_LOSS | chr9p_CNN-LOH | $4.78x10^{-7}$ | 23 (8-51) | 0 |
| chr3_GAIN | chr8_GAIN | $7.19x10^{-7}$ | 33 (10-81) | 0 |
| chr3q_CNN-LOH | chr14q_CNN-LOH | $7.99x10^{-7}$ | 5 (3-9) | 0 |
| chr1_GAIN | chr3_GAIN | $2.46x10^{-6}$ | 47 (12-127) | 0 |
| chr6p_CNN-LOH | chr14q_LOSS | $2.8x10^{-6}$ | 8 (4-16) | 0 |
| chr9_GAIN | chr18p_LOSS | $3.04x10^{-6}$ | 124 (24-419) | 0 |
| chr3p_LOSS | chr15q_LOSS | $3.43x10^{-6}$ | 116 (23-377) | 0 |
| chr7q_LOSS | chr11q_LOSS | $3.45x10^{-6}$ | 16 (6-36) | 0 |
| chr4q_LOSS | chr13q_LOSS | $3.67x10^{-6}$ | 16 (6-35) | 0 |
| chr5q_LOSS | chr17p_CNN-LOH | $5.04x10^{-6}$ | 11 (4-24) | 0 |
| chr9p_CNN-LOH | chr14q_CNN-LOH | $5.51x10^{-6}$ | 3 (2-5) | 0 |
| chr17p_LOSS | chr21q_LOSS | $6.51x10^{-6}$ | 92 (18-288) | 1 |
| chr7_GAIN | chr9p_LOSS | $6.58x10^{-6}$ | 627 (67-2791) | 0 |
| chr11q_LOSS | chr14q_CNN-LOH | $7.33x10^{-6}$ | 3 (2-5) | 0 |
| chr18_GAIN | chr13q_LOSS | $9.47x10^{-6}$ | 13 (5-30) | 0 |

We assess co-occurrence of mosaic events (more than 10 carriers) in different chromosome. Loss and CNN-LOH are evaluated in p,q arm-basis. As a result, 4,299 combinations remain for evaluation. Significance level is set of 0.05/4,299. OR: odds ratio, CI: confidence interval

Table S26. Variants associated with *JAK2* V617F demonstrating associations with chr9p CNN-LOH.

| Gene | SNP | Chrband | Pos | ref | risk | Freqcont | OR (95%CI) | P | shared risk |
|---|---|---|---|---|---|---|---|---|---|
| [JAK2] | rs59384377 | 9p24.1 | 5005034 | A | T | 0.35/0.24 | 1.65 (1.43-1.90) | $3.2x10^{-11}$ | Yes |
| [TERT] | rs7705526 | 5p15.33 | 1285974 | C | A | 0.43/0.36 | 1.32 (1.14-1.53) | $1.9x10^{-4}$ | Yes |
| [TERT] | rs2853677 | 5p15.33 | 1287194 | A | G | 0.36/0.31 | 1.27 (1.10-1.46) | 0.0012 | Yes |
| [SH2B3] | rs7310615 | 12q24.12 | 111865049 | G | C | NA | NA | NA | NA |
| CXXC4—[]—TET2 | rs1548483 | 4q24 | 105749895 | C | T | NA | NA | NA | NA |
| [CHEK2] | rs555607708 | 22q12.1 | 29091857 | I | D | NA | NA | NA | NA |
| [ATM] | rs1800056 | 11q22.3 | 108138003 | T | C | NA | NA | NA | NA |
| [PINT] | rs58270997 | 7q32.3 | 130729394 | C | T | 0.84/0.80 | 1.27 (1.05-1.52) | 0.011 | Yes |
| GFI1B-[]–GTF3C5 | rs621940 | 9q34.13 | 135870130 | C | G | 0.058/0.050 | 1.17 (0.87-1.56) | 0.29 | Yes |

Chrband: chromosome band, Pos: position, ref: reference allele, risk: risk allele, OR: odds ratio, CI: confidence interval, shared risk: shared risk allele (increasing presence of mosaic) between chromosome 9p CNN-LOH and *JAK2* V617F in the Hinds et al.

Table. S27.Candidate analyses of variants associated with MPN, CLL and mLOY in the current study.

| Variant | Chr:Pos | trait | gene | loss | CNN-LOH | gain | ANY |
|---|---|---|---|---|---|---|---|
| rs2736609 | 1:156202640 | mLOY | PMF1,SEMA4A | 0.3994 | 0.5068 | 0.06674 | 0.08883 |
| rs11125529 | 2:54475866 | telo | ACYP2 | 0.48 | 0.01974 | 0.5716 | 0.002883 |
| rs13401811 | 2:111616104 | CLL | ACOXL,BCL2L11 | 0.1493 | 0.6137 | 0.2077 | 0.6464 |
| rs17483466 | 2:111797458 | CLL | ACOXL,BCL2L11 | 0.7407 | 0.3332 | 0.8743 | 0.8827 |
| rs58055674 | 2:111831793 | CLL | ACOXL | 0.5313 | 0.5326 | 0.9672 | 0.7056 |
| rs1439287 | 2:111871897 | CLL | ACOXL,BCL2L11 | 0.4944 | 0.2843 | 0.2888 | 0.5048 |
| rs9308731 | 2:111908262 | CLL | BCL2L11 | 0.3116 | 0.7945 | 0.4168 | 0.2454 |
| rs13015798 | 2:201909515 | CLL | FAM126B,CASP8 | 0.5161 | 0.003709 | 0.4157 | 0.1776 |
| rs3769825 | 2:202111380 | CLL | CASP8,CASP10 | 0.2221 | 0.01237 | 0.1645 | 0.4406 |
| rs13397985 | 2:231091223 | CLL | SP140 | 0.2245 | 0.02097 | 0.7345 | 0.01227 |
| rs9880772 | 3:27777779 | CLL | EOMES | 0.4504 | 0.0824 | 0.06091 | 0.000791 |
| rs115854006 | 3:48388170 | mLOY | TREX1,PLXNB1 | 0.8675 | 0.35 | 0.713 | 0.555 |
| rs13088318 | 3:101242751 | mLOY | SENP7 | 0.2752 | 0.05705 | 0.1488 | 0.7914 |
| rs59633341 | 3:150018880 | mLOY | TSC22D2 | 0.1433 | 0.006182 | 0.4014 | 0.001952 |
| rs2201862 | 3:168648039 | MPN | EGFEM1P,MECOM | 0.789 | 0.242 | 0.8756 | 0.7287 |
| rs10936599 | 3:169492101 | CLL,telo | MYNN | 0.4384 | 0.1494 | 0.252 | 0.009303 |
| rs9815073 | 3:188115682 | CLL | LPP | 0.1216 | 0.09923 | 0.1447 | 0.03627 |
| rs898518 | 4:109016824 | CLL | LEF1 | 0.1337 | 0.5649 | 0.7709 | 0.08433 |
| rs6858698 | 4:114683844 | CLL | CAMK2D | 0.2148 | 0.8426 | 0.9411 | 0.8943 |
| rs56084922 | 5:111061883 | mLOY | NR | 0.5866 | 0.08685 | 0.3625 | 0.9034 |
| rs9391997 | 6:409119 | CLL | IRF4 | 0.8326 | 0.9 | 0.1064 | 0.4043 |

| | | | | | | | |
|---|---|---|---|---|---|---|---|
| rs872071 | 6:411064 | CLL | IRF4 | 0.8762 | 0.9124 | 0.09851 | 0.4127 |
| rs926070 | 6:32257566 | CLL | HLA | 0.4525 | 0.6019 | 0.6346 | 0.5426 |
| rs674313 | 6:32578082 | CLL | HLA-DRB5 | 0.265 | 0.4723 | 0.3235 | 0.3812 |
| rs9273363 | 6:32626272 | CLL | HLA | 0.9069 | 0.4529 | 0.8402 | 0.1018 |
| rs210142 | 6:33546837 | CLL | BAK1 | 0.6582 | 0.1101 | 0.4377 | 0.4298 |
| rs9487023 | 6:109590004 | mLOY | C6orf183 | 0.012 | 0.42 | 0.68 | 0.95 |
| rs13191948 | 6:109634599 | mLOY | SMPD2,CCDC162P | 0.01944 | 0.3357 | 0.6073 | 0.9245 |
| rs2236256 | 6:154478440 | CLL | IPCEF1 | 0.2501 | 0.4082 | 0.4072 | 0.5381 |
| rs381500 | 6:164478388 | mLOY | QKI | 0.3421 | 0.1151 | 0.6336 | 0.4127 |
| rs17246404 | 7:124462661 | CLL | POT1 | 0.08878 | 0.601 | 0.4974 | 0.8552 |
| rs58270997 | 7:130729394 | MPN | PINT | 0.2214 | 0.6965 | 0.8257 | 0.7073 |
| rs2511714 | 8:103578874 | CLL | ODF1,KLF10 | 0.3717 | 0.1984 | 0.5268 | 0.5758 |
| rs2466035 | 8:128211229 | CLL | MYC | 0.02086 | 0.01859 | 0.5271 | 0.9881 |
| rs1679013 | 9:22206987 | CLL | AS1,CDKN2B | 0.9177 | 0.5288 | 0.3195 | 0.7511 |
| rs1359742 | 9:22336996 | CLL | DMRTA1,CDKN2B-AS1 | 0.3594 | 0.7839 | 0.4557 | 0.9172 |
| rs621940 | 9:135870130 | MPN | GFI1B | 0.9617 | 0.293 | 0.6485 | 0.3854 |
| rs1800682 | 10:90749963 | CLL | ACTA,FAS | 0.0329 | 0.1123 | 0.0377 | 0.1962 |
| rs4406737 | 10:90759724 | CLL | ACTA2,FAS | 0.0252 | 0.04858 | 0.1261 | 0.1701 |
| rs9420907 | 10:105676465 | telo | OBFC1 | 0.8338 | 0.4587 | 0.05934 | 0.9254 |
| rs7944004 | 11:2311152 | CLL | TSPAN32 | 0.3873 | 0.1941 | 0.008026 | 0.7445 |
| rs4754301 | 11:108048541 | mLOY | NPAT,ATM,ACAT1 | 0.03653 | 0.005498 | 0.3504 | 0.00014* |
| rs35923643 | 11:123355391 | CLL | GRAMD1B | 0.008576 | 0.09729 | 0.1087 | 0.01234 |
| rs735665 | 11:123361397 | CLL | SCN3B,GRAMD1B | 0.009671 | 0.1022 | 0.1134 | 0.01733 |

| | | | | | | | |
|---|---|---|---|---|---|---|---|
| rs2953196 | 11:123368333 | CLL | NR | 0.7498 | 0.9521 | 0.9432 | 0.2529 |
| rs4251697 | 12:12874462 | mLOY | CDKN1B | 0.029 | 0.67 | 0.88 | 0.86 |
| rs8024033 | 15:40403657 | CLL | BMF | 0.8124 | 0.05056 | 0.6588 | 0.897 |
| rs11636802 | 15:56775597 | CLL | MNS1,RFXDC2 | 0.6555 | 0.1894 | 0.5614 | 0.2737 |
| rs72742684 | 15:56780767 | CLL | MNS1,RFX7 | 0.6554 | 0.2171 | 0.5893 | 0.2809 |
| rs2052702 | 15:69989505 | CLL | PCAT29 | 0.4067 | 0.718 | 0.1411 | 0.6192 |
| rs7176508 | 15:70018990 | CLL | RPLP1 | 0.433 | 0.7141 | 0.1849 | 0.5891 |
| rs12448368 | 16:81044947 | mLOY | CENPN,ATMIN | 0.6864 | 0.9048 | 0.1129 | 0.5769 |
| rs391023 | 16:85927814 | CLL | IRF8 | 0.8238 | 0.1303 | 0.7048 | 0.5291 |
| rs391855 | 16:85928621 | CLL | IRF8 | 0.7976 | 0.2941 | 0.9619 | 0.6437 |
| rs391525 | 16:85944439 | CLL | IRF8 | 0.05825 | 0.49 | 0.87 | 0.09857 |
| rs1044873 | 16:85955671 | CLL | IRF8 | 0.6189 | 0.6304 | 0.6378 | 0.3543 |
| rs77522818 | 17:47817373 | mLOY | FAM117A | 0.201 | 0.1088 | 0.02506 | 0.3552 |
| rs11082396 | 18:42080720 | mLOY | SETBP1 | 0.01109 | 0.1867 | 0.4464 | 0.03494 |
| rs8088824 | 18:42151261 | mLOY | LINC01601;SETBP1 | $1.4x10^{-5}$* | $8.3x10^{-5}$* | 0.21 | $2.1x10^{-5}$* |
| rs4368253 | 18:57622287 | CLL | PMAIP1 | 0.0923 | 0.6578 | 0.6852 | 0.7534 |
| rs4987852 | 18:60793921 | CLL | BCL2 | 0.3739 | 0.1067 | 0.3923 | 0.2347 |
| rs8105767 | 19:22215441 | telo | ZNF208 | 0.06189 | 0.8532 | 0.8549 | 0.4649 |
| rs755017 | 20:62421622 | telo | RTEL1 | 0.9915 | 0.2553 | 0.1523 | 0.8723 |

*indicates significant associations based on Bonferroni's correction (p≤0.00020 (0.05/63/4)).

Table. S28. The significant association between rs8088824 at LINC01601/SETBP1 and mosaic events is largely explained by an association of chr20q loss and chr14q CNN-LOH.

| | p_loss | q_loss | p_CNN-LOH | q_CNN-LOH | gain |
|---|---|---|---|---|---|
| chr1 | 0.6794 | 0.4784 | 0.4907 | 0.123 | 0.1172 |
| chr2 | 0.8038 | 0.8253 | 0.5236 | 0.9201 | 0.2523 |
| chr3 | 0.07043 | 0.416 | 0.8694 | 0.1481 | 0.07884 |
| chr4 | 0.7456 | 0.007841 | 0.2097 | 0.5731 | 1 |
| chr5 | 1 | 0.3354 | 0.6433 | 0.01124 | 1 |
| chr6 | 0.5502 | 0.6356 | 0.1936 | 0.2696 | 0.6435 |
| chr7 | 0.06276 | 1 | 0.3321 | 0.2086 | 0.3204 |
| chr8 | 0.6969 | 0.4065 | 0.773 | 0.315 | 1 |
| chr9 | 1 | 0.1759 | 0.4994 | 0.3613 | 0.4283 |
| chr10 | 1 | 0.09823 | 0.4839 | 0.08764 | 1 |
| chr11 | 0.7561 | 0.7894 | 0.4727 | 0.8373 | 0.8479 |
| chr12 | 0.00992 | 0.1486 | 0.5572 | 0.3042 | 0.3104 |
| chr13 | | 0.2593 | | 0.1381 | 0.3039 |
| chr14 | | 0.04808 | | $1.12 \times 10^{-7}$* | 0.8587 |
| chr15 | | 0.8936 | | 0.2107 | 0.7791 |
| chr16 | 0.01503 | 0.6096 | 0.163 | 0.1443 | 0.554 |
| chr17 | 0.11 | 0.1517 | 0.8683 | 0.106 | 0.1822 |
| chr18 | 0.329 | 0.8123 | 0.4029 | 0.8475 | 0.7412 |
| chr19 | 0.09068 | 0.1409 | 0.1583 | 0.4749 | 0.1653 |
| chr20 | 0.03462 | 0.000147* | 0.8786 | 0.8014 | 0.3342 |
| chr21 | | 0.2825 | | 0.3732 | 0.5544 |
| chr22 | | 0.417 | | 0.0926 | 0.2574 |

45 Bock, C., Walter, J., Paulsen, M. & Lengauer, T. CpG island mapping by epigenome prediction. PLoS Comput. Biol. 3, e110, doi:10.1371/journal.pcbi.0030110 (2007).

46 Satoh, S. et al. AXIN1 mutations in hepatocellular carcinomas, and growth suppression in cancer cells by virus-mediated transfer of AXIN1. Nat. Genet. 24, 245-250, doi:10.1038/73448 (2000).

47 Kagami, M. et al. Deletions and epimutations affecting the human 14q32.2 imprinted region in individuals with paternal and maternal upd(14)-like phenotypes. Nat. Genet. 40, 237-242, doi:10.1038/ng.2007.56 (2008).

48 Miyazato, A. et al. Identification of myelodysplastic syndrome-specific genes by DNA microarray analysis with purified hematopoietic stem cell fraction. Blood 98, 422-427 (2001).

49 Falix, F. A., Aronson, D. C., Lamers, W. H. & Gaemers, I. C. Possible roles of DLK1 in the Notch pathway during development and disease. Biochim. Biophys. Acta 1822, 988-995, doi:10.1016/j.bbadis.2012.02.003 (2012).

50 Soucy, T. A., Dick, L. R., Smith, P. G., Milhollen, M. A. & Brownell, J. E. The NEDD8 Conjugation Pathway and Its Relevance in Cancer Biology and Therapy. Genes Cancer 1, 708-716, doi:10.1177/1947601910382898 (2010).

51 Swords, R. T. et al. Pevonedistat, a first-in-class NEDD8-activating enzyme inhibitor, combined with azacitidine in patients with AML. Blood 131, 1415-1424, doi:10.1182/blood-2017-09-805895 (2018).

52 Jones, M. et al. The shelterin complex and hematopoiesis. J. Clin. Invest. 126, 1621-1629, doi:10.1172/JCI84547 (2016).

53 Paduano, F. et al. T-Cell Leukemia/Lymphoma 1 (TCL1): An Oncogene Regulating Multiple Signaling Pathways. Front. Oncol. 8, 317, doi:10.3389/fonc.2018.00317 (2018).
